# Limited, but potentially functional translation of non-coding transcripts in the HEK293T cellular cytosol

**DOI:** 10.1101/2021.12.23.473848

**Authors:** Annelies Bogaert, Daria Fijalkowska, An Staes, Tessa Van de Steene, Hans Demol, Kris Gevaert

**Affiliations:** VIB Center for Medical Biotechnology, VIB, Ghent, East Flanders, 9052, Belgium; Department of Biomolecular Medicine, Ghent University, Ghent, East Flanders, 9052, Belgium

## Abstract

Ribosome profiling has revealed translation outside of canonical coding sequences (CDSs) including translation of short upstream ORFs, long non-coding RNAs, overlapping ORFs, ORFs in UTRs or ORFs in alternative reading frames. Studies combining mass spectrometry, ribosome profiling and CRISPR-based screens showed that hundreds of ORFs derived from non-coding transcripts produce (micro)proteins, while other studies failed to find evidence for such types of non-canonical translation products. Here, we attempted to discover translation products from non-coding regions by strongly reducing the complexity of the sample prior to mass spectrometric analysis. We used an extended database as the search space and applied stringent filtering of the identified peptides to find evidence for novel translation events. Theoretically, we show that our strategy facilitates the detection of translation events of transcripts from non-coding regions, but experimentally only find 19 peptides (less than 1% of all identified peptides) that might originate from such translation events. Virotrap based interactome analysis of two N-terminal proteoforms originating from non-coding regions finally showed the functional potential of these novel proteins.

## Introduction

A single protein-coding gene may give rise to several protein variants, so-called proteoforms (1). Proteoforms can arise from the usage of different promoters during gene transcription and, in eukaryotes, by differences in processing of immature mRNA molecules (alternative splicing). Also translation events (in-frame alternative translation initiation, ribosomal frameshifting and stop codon read-through), protein modifications including processing by proteases give rise to proteoforms (1,2). Ribosome profiling (Ribo-seq), RNA-sequencing, sequence conservation analysis, bioinformatics prediction tools and proteogenomics have revealed that many transcripts contain more than one ORF (3–7). Such ORFs often do not resemble annotated ORFs, as they can be either situated within 5’ or 3’ untranslated regions (UTRs) or have alternative reading frames that overlap with annotated ORFs. In addition, several ORFs are derived from transcripts that are annotated as non-coding. The latter include, among others, long non-coding RNA (lncRNA), retained introns and transcribed pseudogenes. ORFs currently non annotated to code for proteins are often shorter than 100 codons and are therefore also referred to as small ORFs (sORFs) (6,8–11).

Although such unannotated ORFs gained increased attention over the years, their coding potential and possible biological functions remain a matter of debate. Targeted bioinformatics approaches and several ribosomal profiling approaches enabled the prediction, detection and discovery of thousands of novel ORFs possibly being translated to proteins (3,5,7,8,12–14). However, the peptide and protein products of only a fraction of these have been detected by mass spectrometry (7,15–20). The biases and shortcomings inherent to mass spectrometry were considered as potential causes for the lack of detection of protein products originating from non-coding RNA (21,22). However, in 2017 Verheggen *et al.* (15) showed that such technical aspects alone cannot explain this absence of long-non coding RNA-encoded proteins in mass spectrometry data. This discrepancy between the limited number of detected products from unannotated ORFs in mammalian cells and the large number of unannotated ORFs detected by ribosome profiling and computational methods leaves open the possibility that the protein products of such ORFs are rapidly degraded and therefore not detectable, or are not translated as predicted (7,10,15,23,24). It was also suggested that Ribo-seq overestimates the amount of translation events due to imperfect sequences matching the genome (24).

Some unannotated ORFs function as cis-acting translation controls of annotated ORFs such as upstream ORFs within the 5’UTR (25,26). In contrast, other studies have indicated that unannotated ORFs encode for small proteins with roles in muscle contraction, immune response and mitochondrial functions (8,27–31). Recently, Chen *et al*. (32) studied micropeptides originating from small ORFs on a large scale by combining Ribo-seq, mass spectrometry and CRISPR-Cas based screens. They detected stable expression, localization, knockout and rescue effects, as well as protein interactors of the translation products of six long non-coding RNA’s and seven upstream ORFs (uORFs). For example, a human lncRNA *RP11-84A1.3* was found to encode a 70 amino acid (AA) long protein that localizes to the plasma membrane and interacts with several cell surface proteins. Another study by Ruiz Cuevas *et al*. identified 1,529 peptide products from non-coding ORFs in B-cell lymphoma cells. Of note, these peptides were found to be associated with the MHCI complex and were only found upon analyzing the immunopeptidome. It was therefore predicted that the proteins from which they originated were more disordered and less stable, leading to their rapid degradation by the proteasome, which is the main source for generating MHC I complex-associated peptides (24). In general, such studies point to the protein-coding potential and functional importance of unannotated ORFs.

To improve the detection of novel protein products, proteogenomics approaches were developed that combine more comprehensive sequence databases with techniques to enrich small and/or low abundant proteins in complex samples (10,15,23). The latter because it was found that unannotated ORFs generally have lower transcription and translation rates (24). In this study, we aimed to detect and characterize protein products from annotated non-coding regions/transcripts in HEK293 cells. We created a database containing UniProtKB-SwissProt entries and UniProt isoforms (33) appended with a Ribo-seq based protein database. For this we used two publicly available Ribo-seq datasets from cultured HEK293 cells (4,34), which were processed with Proteoformer 2.0 (35) to derive translation products. We reduced the complexity of the studied proteome by focusing on cytosolic proteins and enriched their N-terminal peptides (36) as, in theory, every protein gets then represented by one peptide (its N-terminal one), allowing the detection of lower abundant proteins (37). Further, protein N-termini hold a lot of information, as most proteins can be identified by their N-terminus alone (38) and N-termini are ideal proxies for studying protein variants (39). We show, using *in silico* studies, that the proteome only contains 3.7% of unique peptides originating from non-coding genes. However, when focusing on N-terminal (Nt-) peptides this percentage raises to 25.4%, thus greatly improving the likelihood of detecting protein products from these non-coding genes. To increase proteome coverage, three different proteases to generate N-terminal peptides were used in parallel (40). Besides reducing sample complexity, enriching for cytosolic proteins comes with the benefit that Virotrap can be used to characterize the protein complexes in which the proteins reside (41).

Beside reducing proteome complexity, Nt-peptide enrichment offers other major advantages for proteoform discovery. The presence of an initiator methionine and the acetylation state of the N-terminus allows to verify if a proteoform originates from translation, as opposed to protein processing, and indicates the exact translation initiation site, allowing the distinction of related proteoforms. Indeed, only nascent proteins start with an initiator methionine (iMet) (42) that can be co-translationally removed by methionine aminopeptidases (MetAPs), exposing the second amino acid at the protein’s N-terminus, and providing a first level of evidence for a protein’s origin being due to a translation event. MetAPs remove iMet when the side-chain of the second amino acid has a small gyration radius (Ala, Cys, Gly, Pro, Ser, Val or Thr) (43,44). Next to the peptide sequence, we monitor protein N-terminal acetylation, a modification that carries another level of evidence for protein synthesis. Nascent polypeptides can be acetylated by N-terminal acetyltransferases (NATs), both on the iMet as well as on newly exposed residues upon iMet removal (45,46) (**Figure 1A**). In human cells, 80-90% of all cytosolic proteins are N-terminally acetylated in this way (39,45,47). After translation, protein processing may occur, including signal or transit peptide removal and many other types of modifications (48). Such post-translational processing events generate new N-termini that are typically not acetylated. COFRADIC allows to distinguish *in vivo* acetylated Nt-peptides originating from translation events from such non-acetylated neo-N-termini(36) (**Figure 1B**). In our study, we thus possess of three levels of evidence indicating if an Nt-peptide can be used as a proxy of translation (**Figure 2**). Based on these three levels of evidence we applied a stringent filtering approach on our cytosolic Nt-data to find high confident, peptide level evidence for the translation of non-coding transcripts. We obtained 2,896 distinct N-termini, with only 19 of them pointing to the translation of non-coding transcripts. Our study thus seems to prove that stringent filtering and careful inspection of proteomics data is required when one aims to identify novel proteins.

**Figure 1:**
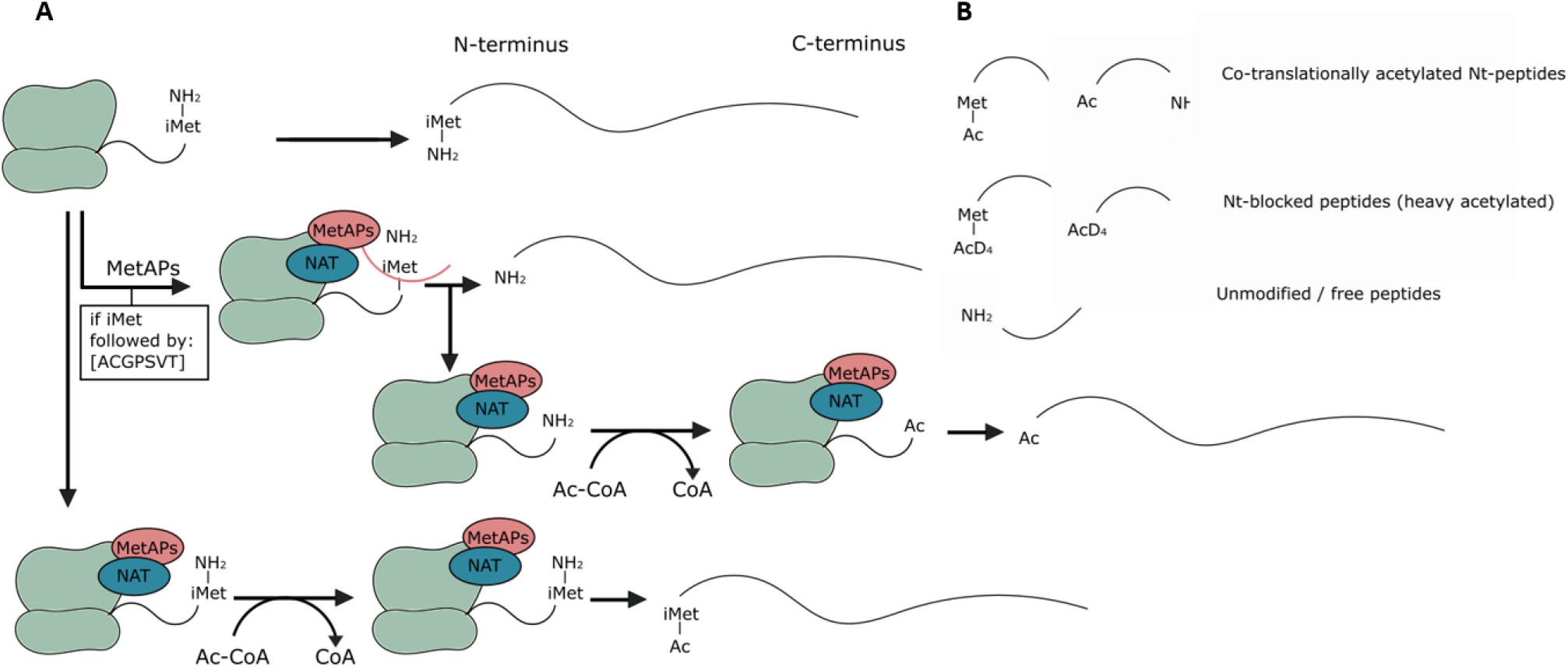
Overview of protein synthesis and co-translational modification, and the types of N-terminal peptides expected. A) Overview of protein synthesis. The first translated amino acid is normally iMet and can be co-translationally removed by MetAPs (their specificity is indicated), exposing the second amino acid as the new protein’s N-terminus. Nascent polypeptides can also be acetylated by N-terminal acetyltransferases (NATs). Depending on the involved NAT, the acetyl group of acetyl coenzyme A is transferred to iMet or to the second residue after iMet removal. B) Overview of the types of peptides expected and their terminology used throughout this paper.

**Figure 2:**
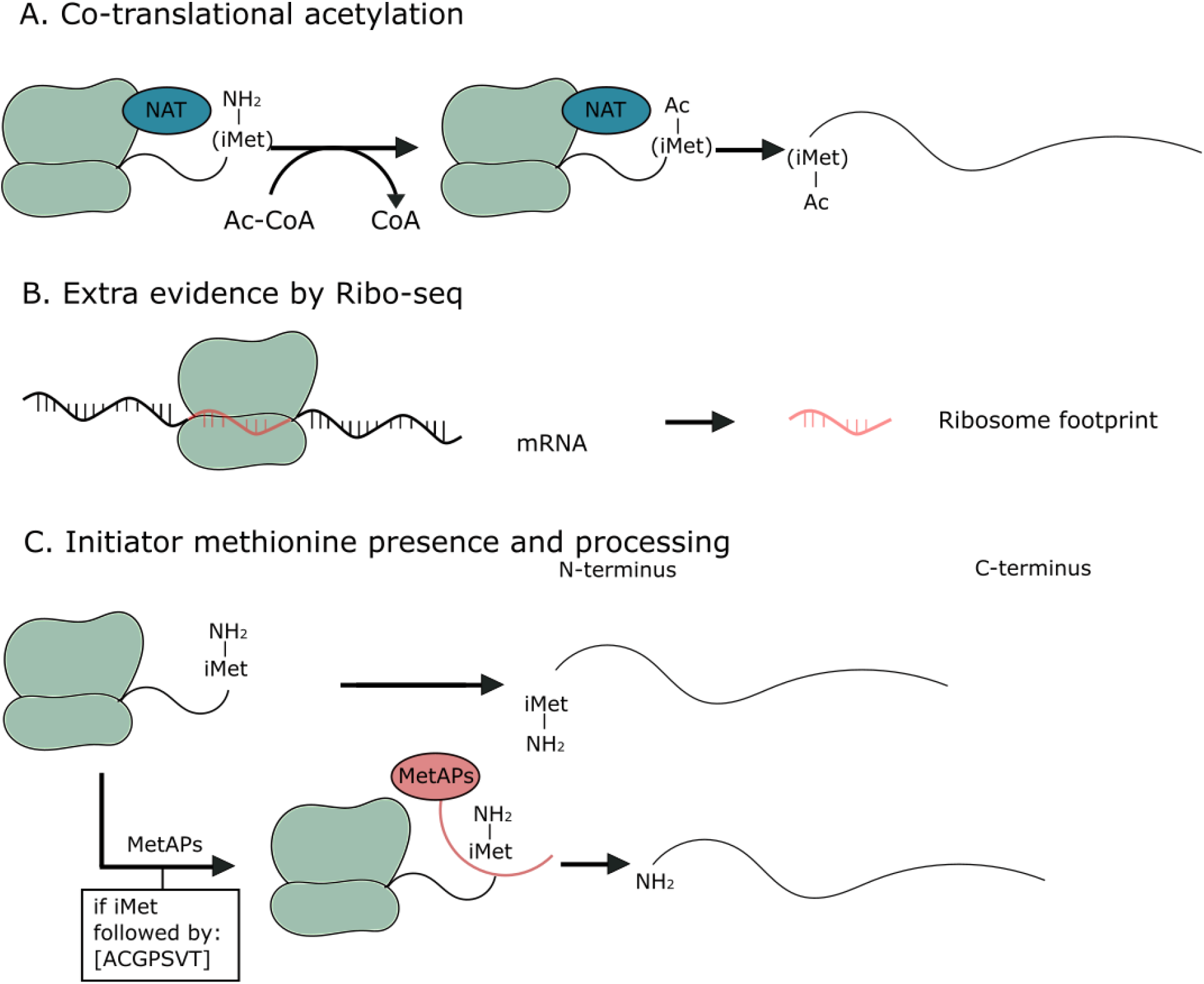
Overview of the different forms of evidence that indicate if an identified peptide contains an N-terminus originating from a translation event. A) In human cells, acetylation of a protein’s N-terminus occurs on 80-90% of all intracellular proteins during translation. Thus, the presence of such an acetyl group is direct translational evidence pointing to a protein’s N-terminus. B) In Ribo-seq, fragments of RNA molecules that are translated are protected by ribosomes (ribosome footprints). Thus, peptides for which there is also Ribo-seq evidence that translation occurred have extra translational evidence. C) Peptides starting with or preceded by a methionine (in accordance with the specificity of the MetAPs) can contain an extra layer of information indicating translation events.

## Materials and Methods

### Custom protein database generation

Ribo-seq reads were downloaded from European Nucleotide Archive (ENA), including the Lee *et al*. (4) dataset collected in HEK293 cells grown under standard conditions (identifiers: SRR618770-SRR618773) as well as the Gao *et al*. (34) datasets of control (SRR1630828, SRR1630831) and amino-acid starved (SRR1630830, SRR1630833) HEK293 cells. Subsequently, Ribo-seq reads pointing to translation initiation with lactimidomycin (LTM) or translation elongation with cycloheximide (CHX) were subjected to the PROTEOFORMER 2.0 pipeline (35) (https://github.com/Biobix/proteoformer) in a pairwise fashion for the corresponding LTM and CHX experiments. Human genome assembly GRCh38.p13 with Ensembl annotation 98 was used to generate indexes for the splice-aware mapper STAR (v. 2.5.4b) with the following settings: --genomeSAindexNbases 14 --sjdbOverhang 35. Common contaminating sequences were retrieved: PhiX bacteriophage genome (NC_001422.1) and human rRNA sequences (search term: ‘(biomol_rrna) AND “Homo sapiens” [porgn:_txid9606]’) were obtained from NCBI, human sn-RNA and sn-o-RNA sequences were obtained from Biomart (v. 98, using gene type filter), whereas human tRNA sequences (v. hg38) were downloaded from gtrnadb.ucsc.edu. The mapping.pl suite of PROTEOFORMER 2.0 allowed integrating several subsequent steps of data processing. Read quality filtering and trimming by fastx toolkit (v. 0.0.14) was performed for the Lee *et al*. dataset to remove polyA adaptors (48x A), whereas no adaptor removal was necessary for the Gao *et al*. dataset. STAR was subsequently used to filter out reads mapping to PhiX, rRNA, sn-(o-)RNA and tRNA, before the remaining reads were mapped non-uniquely to the human genome with the following settings: --readlength 36 --unique N --rpf_split Y --suite plastid. Lastly, Plastid was used to determine P-site offsets. Rule-based transcript calling was performed using Ensembl release 98, resulting in the elimination of transcripts without any reads and classification of other transcripts based on exon coverage. Translation initiation site (TIS) calling was performed in a rule-based manner, jointly using the results of the corresponding LTM and CHX experiments. SNP calling was omitted. Near-cognate start sites were decoded to methionine and known selenocysteines were included. Custom identifiers of proteoforms were composed of Ensembl transcript ID, TIS genomic position and TIS annotation. The following TIS annotations were distinguished: aTIS (proteoform TIS corresponding to the Ensembl annotated TIS); CDS (proteoform TIS falls within the Ensembl annotated coding sequence – CDS); 5UTR (proteoform TIS is located in the 5’UTR); ntr (proteoform TIS is located on a noncoding transcript (NTR) based on Ensembl biotype, such as pseudogene) and 3UTR (proteoform TIS is located in the 3’UTR). A FASTA file of candidate proteoforms was generated for each Ribo-seq experiment, without removing subsequence redundancy (--mflag 5), retaining all Ensembl annotated TIS that did not pass the TIS calling algorithm (--tis_call Y). FASTA files from the Lee *et al*. and Gao *et al*. datasets were subsequently combined (combine_dbs.py) and merged with the UniProt FASTA file (human canonical proteins and isoforms, version 2019_04, 42,425 entries) using the combine_with_uniprot.py module. The origin of the proteoforms is clear from the structure of the FASTA headers: redundant sequences are reduced to a single entry with one main ID and all other IDs kept in the description line, between square brackets. The UniProt ID is preferentially selected as the main ID, and otherwise, the main ID is selected based on the following order of importance: aTIS, CDS, 5UTR, ntr, 3UTR. The last six characters of the main ID are reserved for bincodes denoting the Ribo-seq dataset of origin, with the following order: 1) Lee *et al*. dataset, 2) Gao *et al*. dataset of normal conditions and 3) Gao *et al*. dataset of starved cells. The combined FASTA file was used as a custom protein database for identifying MS/MS spectra.

### Detectability analysis

UniProt and NTR proteoform sequences were selected from the custom database using FASTA headers and the R package Biostring. NTR entries matching completely any proteoform of another category (UniProt, Ensembl aTIS, 5UTR, CDS, 3UTR) were removed. *In silico* protein digestion was performed using the R package cleaver with the following enzyme settings: ArgC specificity “R”, up to 2 missed cleavages (MC); chymotrypsin specificity “[FLMWY]”, up to 2 MC; V8/GluC specificity c(”[DE]”, “[DE](?=P)”), up to 4 MC. Peptide mass and charge at a pH of 2 were calculated using the R package Peptides. *In silico* generated peptides longer than 6 amino acids, with a maximal charge of +4 and a mass to charge ratio (m/z) ≤ 1500 Th, were considered to be MS-detectable peptides. N-terminal peptides starting at position 1 were retrieved from the complete pool of peptides using the R cleaver and IRanges packages. Methionine cleavage was considered and processed N-terminal peptides starting at the second position in the protein sequence were additionally generated if the initiator methionine was followed by A, S, G, P, T or V. N-terminal peptides were also subjected to the MS suitability criteria indicated above. For the uniqueness analysis, the R package stringr was used and leucine residues were considered indistinguishable from isoleucine. MS-detectable peptides were subjected to the DeepMSPeptide Convolutional Neural Network detectability prediction tool using the Python Tensorflow package, version 1.13.1. Data concerning MS detectability and uniqueness were grouped by protein using the R package dplyr. Biotype information was taken from Ensembl Biomart v.98. R packages ggplot2, RColorBrewer, GeomSplitViolin, reshape2, scales, ggExtra, ggsci and GGally were used for plotting.

### Cell culture

Human embryotic kidney cells (HEK293T) were cultured at 37 °C and 8% CO_2_ in DMEM medium, supplemented with 10% fetal bovine serum, 25 units/ml penicillin and 25 µg/ml streptomycin.

### Cytosol extraction

Cytosolic extracts were prepared from 2.5 x10^7^ HEK293T cells similar as described in (49). The cell pellets were washed with ice-cold DPBS (Thermo Fisher Scientific cat. No. 14190250) and resuspended in 1.25 ml of cell-free systems (CFS) buffer (220 mM mannitol, 170 mM sucrose, 5 mM NaCl, 5 mM MgCl_2_, 10 mM HEPES pH 7.5, 2.5 mM KH_2_PO_4_, 0.02 % digitonin and Complete protease inhibitor cocktail (Roche Cat. No. 4693132001)) and kept on ice for 2 min. Lysates were cleared by centrifugation at 14,000 x g for 15 min at 4 °C. The supernatants, being the cytosolic extracts, were collected. The remaining pellets and a HEK293T total lysate serving as control were resuspended in 1 ml of RIPA buffer (50 mM Tris-HCl pH 7.4, 150 mM NaCl, 1% NP40, 0.5% sodium deoxycholate, 0.1% SDS and Complete protease inhibitor cocktail) followed by three freeze-thaw cycles. Lysates were cleared by centrifugation at 16,000 x g for 15 min at room temperature. 200 µl of each sample was used for Western blot analysis, while the rest of the sample was used for N-terminal peptide enrichment by COFRADIC.

### Western blot analysis

Proteins were denatured in XT Sample buffer (Bio-Rad cat. No. 1610791) and XT reducing agent (Bio-Rad cat. No. 1610792), heated at 99 °C for 10 min and centrifuged for 5 min at 16,000 x g (room temperature). 25 µg of each protein mixture was separated by SDS-PAGE (polyacrylamide gel electrophoresis) on a Criterion XT 4-12% Bis-Tris gel (Biorad Laboratories cat. No. 3450124). Proteins were transferred to a PVDF membrane (Merck Millipore cat. No. IPFL00010) after which the membrane was blocked using Odyssey Blocking buffer (PBS) (LI-COR cat. No. 927-4000) diluted once with TBS-T (TBS supplemented with 0.1% Tween 20). Immunoblots were incubated overnight with primary antibodies against GAPDH (Abcam, ab8245), HSP60 (Santa Cruz, sc-13115), Lamin B (Santa Cruz, sc-374015), RibophorinI (Santa Cruz, sc-12164) and γ-Tubulin (Thermo Fisher Scientific, MA1-850) in Odyssey Blocking buffer (PBS) diluted once with TBS-T. Blots were washed four times with TBS-T, incubated with fluorescent-labeled secondary antibodies (IRDye 800CW Goat Anti-Mouse IgG polyclonal 0.5 mg from LI-COR, Cat. No 926-32210) and IRDye 800CW Donkey Anti-Goat IgG polyclonal 0.5 mg from LI-COR cat. No. 926-32214) in Odyssey blocking buffer diluted once with TBS-T for 1 h. After three washes with TBS-T and an additional wash in TBS, immunoblots were imaged using the Odyssey infrared imaging system (LI-COR).

### N-terminal COFRADIC

N-terminal peptides were enriched by COFRADIC as described previously (50) however without the pyroglutamate removal and SCX steps. In the following, we only mention the main differences with the published protocol. In brief, 1 mg of cytosolic proteins was used. As digitonin was used for cytosolic extraction in combination with the RIPA lysis buffer, which interfere with LC-MS/MS analysis, the samples were cleaned-up using Pierce^®^ Detergent removal spin columns (Thermo Fisher Scientific, cat. No. 87777) according to the manufacturer’s instructions. Then, guanidinium hydrochloride was added to a final concentration of 4 M before proteins were reduced (with 15 mM f.c. (final concentration) TCEP) and alkylated (with 30 mM f.c. iodoacetamide) for 15 min at 37 °C. To enable the assignment of *in vivo* Nt-acetylation events, all primary protein amines were blocked using stable isotope encoded acetate, i.e. an NHS-ester of ^13^C_1_D_3_-acetate. This acetylation reaction was allowed to proceed for 1 h at 30 °C and was repeated once. Prior to digestion, the samples were desalted on a NAP-10 column in 50 mM freshly prepared ammonium bicarbonate pH 7.8. Samples were digested either with trypsin in a trypsin/protein ratio of 1/50 (w/w) (Promega Cat. No. V5111) and incubated overnight at 37 °C, chymotrypsin in a chymotrypsin/protein ratio of 1/20 (w/w) (Promega Cat. No. V1061) and incubated overnight at 25 °C or endoproteinase GluC in a GluC/protein ratio of 1/20 (w/w) (Promega Cat. No.V1651) and incubated overnight at 37 °C. After vacuum drying, the samples were re-dissolved in 80 µl loading solvent A (2% acetonitrile (ACN), 0.1% TFA in ddH_2_O) before isolating N-terminal peptides by two subsequent RP-HPLC fractionations with a TNBS reaction in between.

### LC-MS/MS analysis of N-terminal peptides and peptide identification

LC-MS/MS analysis was similar as reported before (50). Each COFRADIC fraction was solubilized in 20 µl loading solvent A and half of each fraction was injected for LC-MS/MS analysis on an Ultimate 3000 RSLCnano system in-line connected to an Orbitrap Fusion Lumos mass spectrometer (Thermo Scientific, Germany). Trapping was performed at 10 μl/min for 4 min in loading solvent A on a 20 mm trapping column (made in-house, 100 μm internal diameter (I.D.), 5 μm beads, C18 Reprosil-HD, Dr. Maisch, Germany). The peptides were separated on a 200 cm µPAC™ column (C18-endcapped functionality, 300 µm wide channels, 5 µm porous-shell pillars, inter pillar distance of 2.5 µm and a depth of 20 µm; Pharmafluidics, Belgium). The column was kept at a constant temperature of 50 °C. Peptides were eluted by a linear gradient reaching 33% MS solvent B (0.1% formic acid (FA) in water/acetonitrile (2:8, v/v)) after 42 min, 55% MS solvent B after 58 min and 99% MS solvent B at 60 min, followed by a 10-minutes wash at 99% MS solvent B and re-equilibration with MS solvent A (0.1% FA in water). The first 15 min, the flow rate was set to 750 nl/min after which it was kept constant at 300 nl/min.

The mass spectrometer was operated in data-dependent mode, automatically switching between MS and MS/MS acquisition. Full-scan MS spectra (300-1500 m/z) were acquired in 3 s acquisition cycles at a resolution of 120,000 in the Orbitrap analyzer after accumulation to a target AGC value of 200,000 with a maximum injection time of 250 ms. The precursor ions were filtered for charge states (2-7 required), dynamic range (60 s; ± 10 ppm window) and intensity (minimal intensity of 5E3). The precursor ions were selected in the ion routing multipole with an isolation window of 1.6 Da and accumulated to an AGC target of 10E3 or a maximum injection time of 40 ms and activated using CID fragmentation (35% NCE). The fragments were analyzed in the Ion Trap Analyzer at rapid scan rate.

The generated MS/MS peak lists were searched with Mascot using the Mascot Daemon interface (version 2.6.0, Matrix Science, Boston, MA). MS data were matched against our custom build database (containing UniProt, UniProt isoform entries appended with Ribo-seq derived protein sequences). The Mascot search parameters were as follows: heavy acetylation of lysine side chains (with ^13^C_1_D_3_-acetate), carbamidomethylation of cysteine and methionine oxidation to methionine-sulfoxide were set as fixed modifications. Variable modifications were acetylation of N-termini (both light and heavy due to the ^13^C_1_D_3_ label) and pyroglutamate formation of N-terminal glutamine (both at the peptide level). The enzyme settings were: endoproteinase semi-Arg-C/P (semi-Arg-C specificity with Arg-Pro cleavage allowed) allowing for two missed cleavages for the trypsin sample. For chymotrypsin and GluC, the enzyme settings were semi-Chymo and semi-GluC. For GluC, two missed cleavages were allowed while for chymotrypsin four missed cleavages were allowed. Mass tolerance was set to 10 ppm on the precursor ion and to 0.5 Da on fragment ions. In addition, the C13 setting of Mascot was set to 1. Peptide charge was set to 1+, 2+, 3+ and instrument setting was put to ESI-TRAP. Raw DAT-result files of MASCOT were further queried using ms_lims (51). Only peptides that were ranked first and scored above the threshold score set at 99% confidence were withheld. The FDR was estimated by searching a decoy database (a reversed version of the custom generated database), which resulted in an FDR of 0.44% for the trypsin sample, 0.14% for the chymotrypsin sample and 0.53% for the GluC sample.

### Selection of N-termini

From this dataset, N-terminal peptides were selected and classified. The selection workflow was built in KNIME (see https://www.knime.com/). Selection was done per protease and all identified peptides (co-translationally acetylated, heavy acetylated (blocked) peptides and N-terminally free peptides) were used as input. Peptides were grouped based on sequence and accession to get a list of distinct (unique) identified peptides. Information on multiple identifications of a given peptide was retained and, if possible, used to calculate an acetylation percentage. Internal (solely found as free, NH_2_-starting peptide) and C-terminal peptides were removed. The remaining potential N-terminal peptides were classified. High confident TIS/N-termini encompass: 1) all (partially) co-translationally (*in vivo* acetylated) N-termini and blocked (*in vitro* heavy acetylated) N-termini of which the start position corresponded with a UniProt, UniProt isoform or Ensembl (Ribo-seq) annotated TIS site; 2) co-translationally acetylated peptides with a start position higher than two, and for which the iMet is retained or removed; 3) N-termini matching TIS identified by ribosome profiling (either co-translationally acetylated or heavy acetylated (blocked)). Low confident TIS/N-termini encompass: 1) co-translationally acetylated peptides with a start position beyond position two that do not start nor are preceded by a Met, with no extra Ribo-seq evidence and that are not preceded by a cleavage site recognized by the proteases used; 2) heavy acetylated (blocked) peptides with a start position higher than two that start with or are preceded by a Met (according to the iMet processing rules), with no extra Ribo-seq evidence and that are not preceded by a proteolytic cleavage site. In a final step, the data from the three different proteases were merged to create a final list of distinct N-termini.

### Synthetic peptides

Two peptides (ADDAGAAGGPGGPGGPEMGNRGGFRGGF and MDGEEKTCGGCEGPDAMYVKLISSDGHEFIVKR) were made in-house, while all other peptides were obtained from Thermo Scientific (standard peptide custom synthesis service). In-house peptide synthesis was done using Fmoc-chemistry on an Applied Biosystems 433A Peptide Synthesizer. All required modifications, besides heavy acetylation of primary amines, were introduced during peptide synthesis. Primary amines were blocked after peptide synthesis by adding a 150 times molar excess of an NHS-ester of ^13^C_1_D_3_-acetate, and peptides were incubated for 1 h at 37 °C. This step was repeated once, after which the remainder of the NHS-ester was quenched by adding glycine to a final concentration of 30 mM and incubating the peptides for 10 min at room temperature. O-acetylation was reversed by adding hydroxylamine (75 mM f.c.) followed by an incubation for 10 min at room temperature. Next, peptides were purified on OMIX C18 Tips (Agilent) which were first washed with pre-wash buffer (0.1% TFA in water/acetonitrile (20:80, v/v) and pre-equilibrated with 0.1% TFA before sample loading. Tips were then washed with 0.1% TFA and peptides were eluted with 0.1% TFA in water/acetonitrile (40:60, v/v). Purified peptides were mixed and diluted to a final concentration of 100 fmol/µl (of each peptide).

1 pmol of the acetylated synthetic peptides was injected for LC-MS/MS analysis on an Ultimate 3000 RSLCnano system in-line connected to an Orbitrap Fusion Lumos mass spectrometer (Thermo). Trapping was performed at 10 μl/min for 4 min in loading solvent A on a 20 mm trapping column (made in-house, 100 μm internal diameter (I.D.), 5 μm beads, C18 Reprosil-HD, Dr. Maisch, Germany). The peptides were separated on a 200 cm µPAC™ column (C18-endcapped functionality, 300 µm wide channels, 5 µm porous-shell pillars, inter pillar distance of 2.5 µm and a depth of 20 µm; Pharmafluidics, Belgium). The column was kept at a constant temperature of 50 °C. Peptides were eluted by a linear gradient reaching 26.4% MS solvent B after 20 min, 44% MS solvent B after 25 min and 56% MS solvent B at 28 min, followed by a 5-minutes wash at 56% MS solvent B and re-equilibration with MS solvent A. The first 15 min, the flow rate was set to 750 nl/min after which it was kept constant at 300 nl/min. The mass spectrometer was operated in data-dependent mode, automatically switching between MS and MS/MS acquisition with the m/z-values of the precursors of the synthetic peptides as an inclusion list. Full-scan MS spectra (300-1500 m/z) were acquired in 3 s acquisition cycles at a resolution of 120,000 in the Orbitrap analyzer after accumulation to a target AGC value of 200,000 with a maximum injection time of 30 ms. The precursor ions not present in the inclusion list were filtered for charge states (2-7 required) and intensity (minimal intensity of 5E3). The precursor ions were selected in the ion routing multipole with an isolation window of 1.6 Da and accumulated to an AGC target of 10E3 or a maximum injection time of 40 ms and activated using CID fragmentation (35% NCE). The fragments were analyzed in the Ion Trap Analyzer at rapid scan rate.

The data analysis software Skyline ((52), Skyline-Daily V21.1.1.316, was used to compare the ranking of the fragment ions between the synthetic peptides and the possible NTR peptides. For each synthetic peptide, the top 10 most abundant fragment ions of the synthetic peptides were selected to perform the comparison. A previously identified N-terminal peptide was considered to matching a synthetic peptide if the ranking of the fragment ions was in line with the ranking of the fragment ions of the synthetic peptide.

### Generation of the NTR clones

Gag-bait fusion constructs were generated as described (41). The coding sequences for the full length and the proteoform of the selected gene (Ensembl accession: ACTBP8, ENSG00000220267) were ordered from IDT (gBlocks gene fragments) and transferred into the pMET7-GAG-sp1-RAS plasmid by classic cloning with restriction enzymes. The pMD2.G (expressing VSV-G), pcDNA3-FLAG-VSV-G plasmids (available at Addgene #12259 and #80606) and the GAG-eDHFR vector (serving as a control) were a gift from Sven Eyckerman (VIB-UGent Center for Medical Biotechnology, Ghent, Belgium).

### Protein complex purification by Virotrap

For full details on the Virotrap protocol we refer to (41). HEK293T cells were kept at low passage (<10) and cultured at 37 °C and 8% CO_2_ in DMEM, supplemented with 10% fetal bovine serum, 25 units/ml penicillin and 25 μg/ml streptomycin. Each construct was analyzed in triplicate and for every replicate, the day prior to transfection, a 75 cm^2^ falcon was seeded with 9×10^6^ cells. Cells were transfected using polyethylenemine (PEI), with a DNA mixture containing 6.43 μg of bait plasmid (pMET7-GAG-bait), 0.71 μg of pcDNA3-FLAG-VSV-G plasmid and 0.36 μg of pMD2.G plasmid. For the eDHFR control, cells were transfected with a DNA mixture containing 3.75 μg of eDHFR plasmid (pMET7-GAG-eDHFR), 2.68 µg of pSVsport plasmid, 0.71 μg of pcDNA3-FLAG-VSV-G plasmid and 0.36 μg of pMD2.G plasmid. The medium was refreshed after 6 h with 8 ml of supplemented DMEM.

The cellular supernatant was harvested after 46 h and centrifuged for 3 min at 1,250 x g to remove debris. The cleared supernatant was then filtered using 0.45 μm filters (Merck Millipore Cat. No. SLHV033RB). For every sample, 20 μl MyOne Streptavidin T1 beads in suspension (10 mg/ml, Thermo Fisher Scientific, Cat. No. 65601) were first washed with 300 μl wash buffer containing 20 mM Tris-HCl pH 7.5 and 150 mM NaCl, and subsequently pre-loaded with 2 μl biotinylated anti-FLAG antibody (BioM2, Sigma, cat. No. F9291). This was done in 500 μl wash buffer and the mixture was incubated for 10 min at room temperature. Beads were added to the samples and the viral-like particles were allowed to bind for 2 h at room temperature by end-over-end rotation. Bead-particle complexes were washed once with 200 µl washing buffer (20 mM Tris-HCl pH 7.5 and 150 mM NaCl) and subsequently eluted with FLAG peptide (30 min at 37 °C; 200 μg/ml in washing buffer, Sigma Cat. No. F3290) and lysed by addition of Amphipol A8–35 (Anatrace, cat. No. A835) (53) to a final concertation of 1 mg/ml. After 10 min the lysates were acidified (pH <3) by adding 2.5% formic acid (FA). Samples were centrifuged for 10 min at >20.000 x g to pellet the protein/Amphipol A8–35 complexes. The supernatant was removed and the pellet was resuspended in 20 µl 50 mM fresh triethylammonium bicarbonate (TEAB). Proteins were heated at 95 °C for 5 min, cooled on ice to room temperature for 5 min and digested overnight at 37 °C with 0.5 μg of sequencing-grade trypsin (Promega Cat. No. V5111). Peptide mixtures were acidified to pH 3 with 1.5 µl 5% FA. Samples were centrifuged for 10 min at 20,000 x g. 7.5 µl of the supernatant was injected for LC-MS/MS on an Ultimate 3000 RSLCnano system in-line connected to a Q Exactive HF Biopharma mass spectrometer (Thermo Scientific, Germany). Trapping was performed at 10 μl/min for 4 min in loading solvent A on a 20 mm trapping column (made in-house, 100 μm internal diameter (I.D.), 5 μm beads, C18 Reprosil-HD, Dr. Maisch, Germany). The peptides were separated on a 250 mm Waters nanoEase M/Z HSS T3 Column, 100 Å, 1.8 µm, 75 µm inner diameter (Waters Corporation, UK) kept at a constant temperature of 50 °C. Peptides were eluted by a non-linear gradient starting at 1% MS solvent B reaching 55% MS solvent B in 80 min, 97% MS solvent B in 90 minutes followed by a 5-minute wash at 97% MS solvent B and re-equilibration with MS solvent A. The mass spectrometer was operated in data-dependent mode, automatically switching between MS and MS/MS acquisition for the 12 most abundant ion peaks per MS spectrum. Full-scan MS spectra (375-1500 m/z) were acquired at a resolution of 60,000 in the Orbitrap analyzer after accumulation to a target value of 3,000,000. The 12 most intense ions above a threshold value of 13,000 were isolated with a width of 1.5 m/z for fragmentation at a normalized collision energy of 30% after filling the trap at a target value of 100,000 for maximum 80 ms. MS/MS spectra (200-2000 m/z) were acquired at a resolution of 15,000 in the Orbitrap analyzer.

The generated MS/MS spectra were processed with MaxQuant (version 1.6.17.0) using the Andromeda search engine with default search settings, including a false discovery rate set at 1% on both the peptide and protein level. The sequences of the human proteins in the Swiss-Prot database (release Jan 2021; complemented with GAG, VSV-G, eDHFR and the NTR (both FL and proteoform) sequences for Virotrap) were used as the search space. The enzyme specificity was set at trypsin/P, allowing for two missed cleavages. Variable modifications were set to oxidation of methionine residues and N-terminal protein acetylation. Standard settings were used. In the settings of advanced identification, match between runs was implemented (with standard settings). Only proteins with at least one unique or razor peptide were retained, leading to the identification of 1,569 proteins across all samples. Reverse proteins, proteins that are only identified by site and potential contaminants were removed. Differential analysis of the Virotrap data was conducted using the limma R package (version 3.48.0) (54). Proteins quantified with iBAQ values in all replicates of at least one condition were retained. Samples were log_2_ transformed and normalized to a common median. Missing values were imputed using imputeLCMD R package (version 2.0) from a truncated distribution with parameters estimated using quantile regression. Pairwise contrasts of interest between differentially treated samples were retrieved at a significance level alpha 0.01, corresponding to Benjamini-Hochberg (BH) adjusted p-value (FDR) cutoff. Z-score transformed iBAQ values were compared, clustered and presented as a heatmap using the pheatmap package (version 1.0.12). Other visualizations were generated using ggplot2 (version 3.3.3), ggrepel (0.9.1) and RColorBrewer (1.1-2).

## Results

### 1. A comprehensive sequence database for identifying novel proteoforms

A comprehensive database of known and putative protein sequences is essential for the mass spectrometry-based identification of (novel) proteoforms. We used publicly available ribosome profiling (Ribo-seq) datasets from HEK293 cells (4,34) as Ribo-seq involves deep sequencing of ribosome-protected transcripts (55) and, combined with drugs that halt initiating ribosomes, Ribo-seq allows to detect translation initiation sites (TIS) (4), including those of novel proteins and Nt-proteoforms (56–58).

Ribo-seq data were processed using Proteoformer 2.0 (35). The resulting (putative) protein sequences were stored in a database that was supplemented with annotated proteoforms from Ensembl and UniProt, both canonical sequences (a curated selection including one protein per gene) and sequences of annotated isoforms. Our final database contained 103,020 non-redundant protein sequences (**Figure 3A**), including 60,043 (58.4%) annotated sequences, 25,564 (24.8%) new sequences of proteoforms in known translated transcripts and 16,919 (16.4%) putative proteoforms originating from noncoding transcripts (NTR). We classified novel predicted proteoforms in protein-coding transcripts according to the position of the TIS, and most were found in 5’ untranslated regions (5’UTR; 21,008 entries), followed by TIS within coding sequences (CDS; 3,555 entries) and those in 3’ untranslated regions (3’UTR; 1,001 entries). NTRs were classified according to the Ensembl biotype and mostly originated from processed pseudogenes (pseudogenes generated through a genome insertion of reverse-transcribed mRNA, possibly with evidence of locus-specific transcription; 7,519 entries), transcripts with retained intron (5,887 entries) and long noncoding RNAs (lncRNA; 3,192 entries; see **Figure 3A**), with only 321 NTRs belonging to other biotypes.

**Figure 3:**
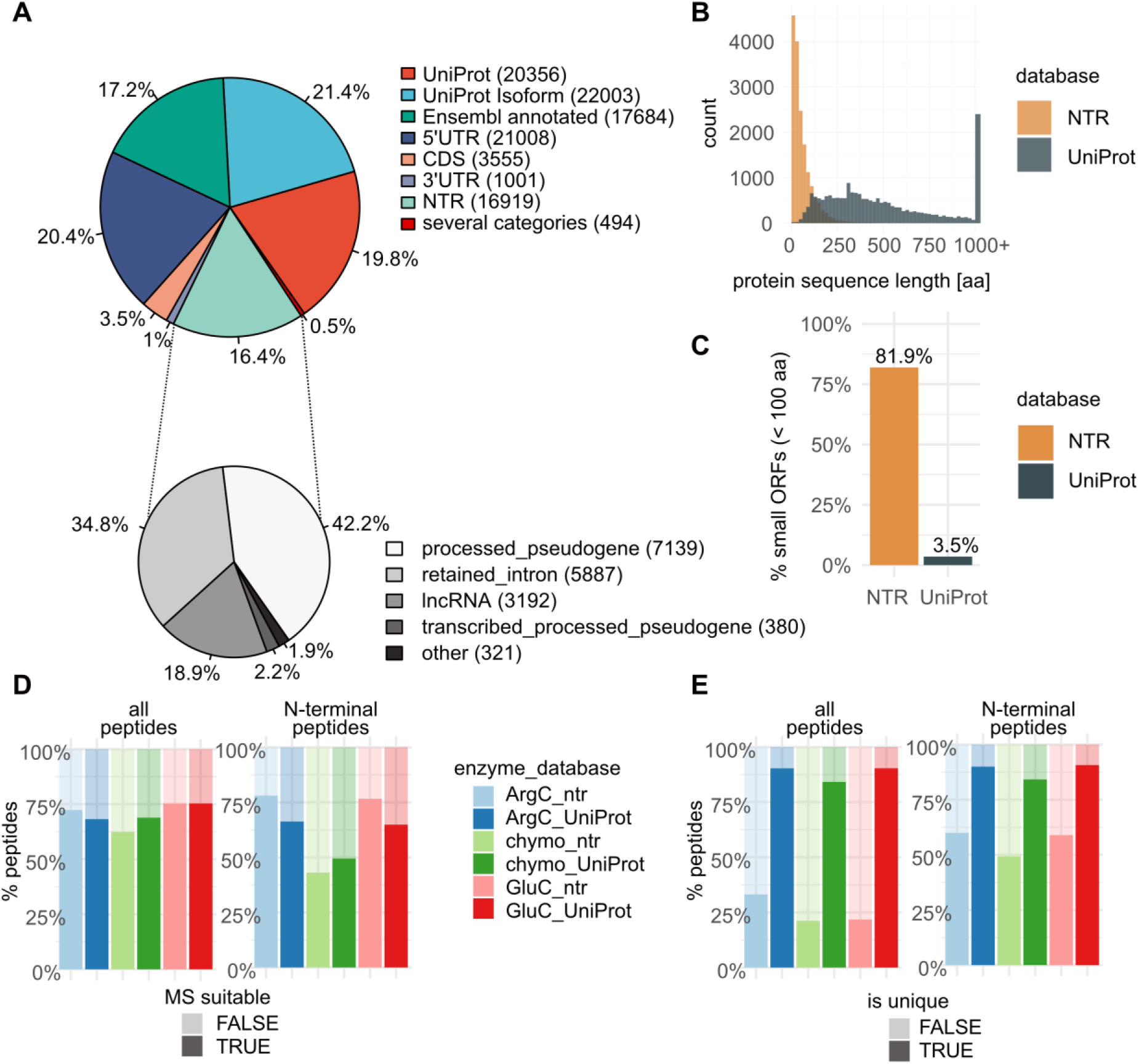
A customized protein sequence database containing NTR and UniProt proteoforms. A) The customized database contains UniProt and Ensembl annotated entries, next to Ribo-seq predicted proteins and proteoforms belonging to several categories. B) NTR proteoforms are significantly shorter than UniProt proteins (median protein length of 40 and 414, respectively; Wilcoxon test p-value < 2.2e-16). C) Most NTR proteoforms are derived from small open reading frames (small ORFs), coding for proteins with lengths less than 100 amino acids. D) The majority of predicted peptides are MS-identifiable, except for chymotrypsin-generated N-terminal peptides. E) When all peptides are considered, NTR peptides are rarely unique compared to those from UniProt proteins. However, when only N-terminal peptides are considered, almost half of the NTR proteins can be uniquely identified by their N-terminal peptide alone. For UniProt proteins, there is almost no difference in the % of unique peptides when considering only N-terminal peptides or all peptides.

### 2. Assessing the MS detectability of proteins

We hypothesized that experimental procedures could be optimized to improve the chances of detecting NTR proteins. Therefore, we calculated and compared NTR and UniProt protein sequence features, such as length, number of protease cleavage sites, MS-identifiable and unique peptides.

NTR proteins were found to be significantly shorter than UniProt proteins (median protein lengths of 40 and 414 respectively, Wilcoxon test p-value < 2.2e-16, **Figure 3B**), with >80% of NTR proteins derived from small ORFs (less than 100 amino acids; **Figure 3C**).

To predict the impact of the protease used for protein digestion and peptide enrichment strategies on protein sequence coverage and NTR protein identification, we performed *in silico* digestion on NTR and UniProt proteins with three different enzymes: endoproteinase ArgC, chymotrypsin and endoproteinase GluC. Note that because lysine side-chains are acetylated in our set-up, trypsin (that will be used for enriching Nt-peptides (see further)) will only cleave C-terminal to arginine, explaining why we studied the effect of endoproteinase ArgC. We considered both shotgun proteomics and N-terminomics approaches and only peptides longer than 6 amino acids, with a maximal charge of 4+ (at a pH of 2), and a mass to charge ratio (*m/z*) ≤ 1500 Th were considered to be MS-idenitfiable. These parameters correspond to the database search outcomes typically obtained in our experiments (see further). The majority of the predicted peptides were found to be MS-identifiable, except for chymotrypsin-generated Nt-peptides which were predicted to have an average peptide length of only 8 AA for NTR proteins and 9 AA for UniProt proteins. Further, the NTR and UniProt proteins produced comparable fractions of MS-identifiable peptides for all conditions considered (**Figure 3D**). However, in contrast to UniProt peptides, NTR peptides were rarely unique (**Figure 3E**), meaning that they matched to more than one protein sequence among all NTR protein sequences. This effect was less pronounced for NTR Nt-peptides (**Figure 3E**).

We next analyzed a theoretical proteome composed of both UniProt and NTR proteoforms, and found that upon digestion, the corresponding peptide mixture is dominated by UniProt peptides and leaves only marginal opportunity to identify unique NTR peptides (1.4 – 3.7%; **Figure 4A**). Note that we do not consider any additional possible disadvantages as for instance caused by lower expression of NTR proteins (15), so the actual detectability of NTR proteins is likely even over-estimated, but enrichment of Nt-peptides seems to increase the likelihood of identifying NTR proteins (**Figure 4B**). Unique peptides offer the strongest evidence of expression of a given proteoform. However, not every proteoform gives rise to unique peptides, inevitably leading to reduced proteome coverage. In contrast to 98% of UniProt proteins that produce at least one unique peptide, up to a third of NTR proteins cannot be uniquely identified (**Figure 4C**). Enrichment of Nt-peptides seems to have an additional negative impact on the number of unique protein identifications, in both UniProt and NTR categories (**Figure 4D**), due to the lack of unique and MS-identifiable N-termini.

**Figure 4:**
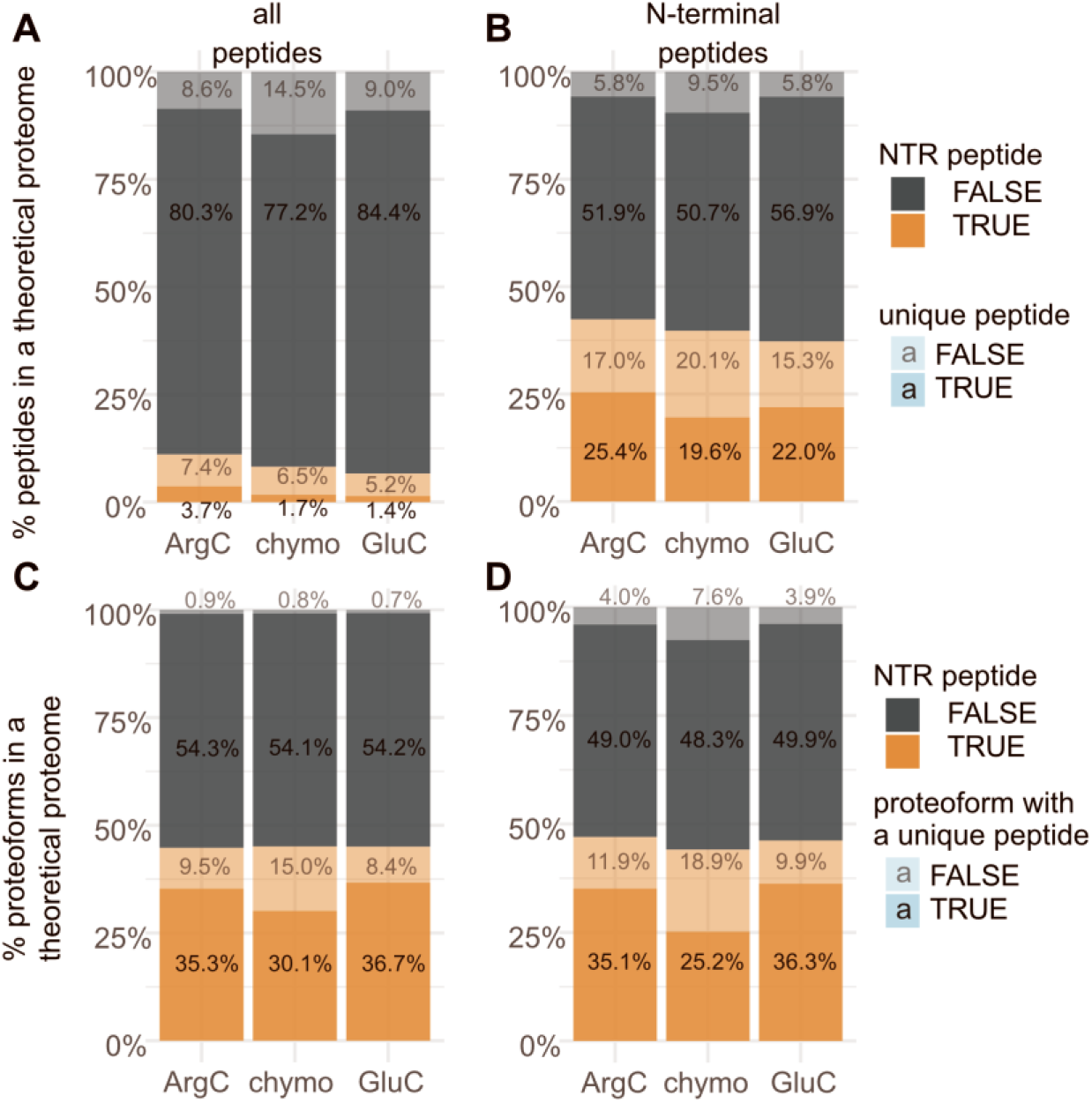
Analysis of *in silico* digests of a theoretical proteome composed of UniProt and NTR sequences. A) In shotgun proteomics, UniProt peptides dominate the digest, leaving marginal chances to identify unique NTR peptides (1.4 – 3.7%, dependent on the protease used). B) Enrichment of N-terminal peptides enhances NTR identification. C) Most UniProt proteins (> 98%) produce at least one unique peptide, whereas > 30% of NTR proteoforms cannot be uniquely identified. D) Enrichment of N-terminal peptides may lead to more inconclusive identifications than using a shotgun proteomics approach.

To conclude this part, it is predicted that by enriching for Nt-peptides, one is more likely to identify NTR proteins, albeit at a potential cost of fewer UniProt annotated proteins.

Next, we evaluated if combining the results after digesting proteomes with different proteases would lead to an overall higher coverage of proteomes. We investigated the proteases mentioned above and found that Nt-peptides enriched after ArgC digestion should capture 89.1% and 95.4% of UniProt and NTR proteins respectively, and both fractions further increase when also including the chymotrypsin and GluC digestion results (**Figure S1**). When considering all MS-identifiable peptides, ArgC provides almost full coverage of UniProt proteins; 99.6% of the proteins generate at least one MS-identifiable peptide. Additionally, this coverage is only increased to a limited extent when including the other two proteases. A somewhat less complete ArgC coverage of NTR proteins (97.4%) in the shotgun proteomics setup can be remedied using complementary digestion strategies.

### 3. Extracting cytosolic proteins

Since decreasing the proteome complexity increases the possibility of obtaining protein evidence from NTR proteins (39) and since 80-90% of all cytosolic proteins are co-translationally acetylated (thus providing translational evidence) (45,47), we decided to isolate cytosolic proteins, enrich their Nt-peptides (59) and perform a highly stringent downstream data analysis. Virotrap (see further), a technology that favors cytosolic proteins as baits, would then be suited to identify potential protein interactors of newly discovered proteins. However, Virotrap is currently limited to HEK293T cells, which explains the selection of these cells.

Cytosolic proteins were enriched after permeabilization of the plasma membrane with 0.02% digitonin, leaving the organellar membranes intact (49). The efficiency of the cytosol isolation was evaluated by Western blot analysis of the cytosolic fraction, the remaining pellet (containing the organelles) and a total cell lysate (as control), using antibodies against several organelle markers (endoplasmic reticulum (ER), cytosol, mitochondrion, cytoskeleton and nucleus) (**Figure S2**). Organelle markers were absent or strongly depleted in the cytosolic fraction, whilst the cytosolic marker was enriched in the cytosolic fraction and depleted in the organelle fraction, indicative of an efficient isolation of cytosolic proteins.

Following COFRADIC, Nt-peptides were analyzed by LC-MS/MS. We also evaluated the quality of cytosol isolation by evaluating Gene Ontology Cellular Component (GOCC) terms associated with the identified proteins for GO terms containing “cytosol” (GO:0005829) (**Table 1**). Among the three samples, a comparable fraction of cytosolic proteins (around 60%) was found. This number is higher than expected when analyzing a total lysate (Fisher’s exact test, p-value < 1e-5), given that The Human Protein Atlas (Cell Atlas) reports that 4,740 proteins (24% of the human proteome) localizes to the cytosol (60). In conclusion, both Western blot and GOCC data indicate that the proteome sample was strongly enriched for cytosolic proteins.

**Table 1:** Overview of proteins that have a GOCC term containing cytosol, lack this term, do not have any GOCC term or no UniProtKB accession at all (which can thus also cannot be matched to a GOCC term). * with ArgC specificity.

|  | Trypsin* digested sample |  | Chymotrypsin digested sample |  | GluC digested sample |  |
| --- | --- | --- | --- | --- | --- | --- |
|  | # | % | # | % | # | % |
| Cytosolic GOCC term | 1,856 | 55.4 | 1,318 | 60.3 | 1,192 | 61.2 |
| Non-cytosolic GOCC term | 1,181 | 35.3 | 708 | 32.4 | 657 | 33.7 |
| No GOCC term | 158 | 4.7 | 123 | 5.6 | 69 | 3.5 |
| No UniProtKB accession | 75 | 4.6 | 37 | 1.7 | 31 | 1.6 |
| Total | 3,270 | 100 | 2,186 | 100 | 1,949 | 100 |

### 4. Increasing proteome coverage by using three proteases and N-terminal peptide enrichment

Prior to digestion, primary amines in the cytosolic proteins were acetylated with an acetyl group carrying stable heavy isotopes, this to distinguish *in vivo* N-terminally acetylated (Nt-acetylated) from *in vivo* free N-termini, and both can serve as a proxy for translation initiation events (61). As mentioned, three different proteases, trypsin (with ArgC specificty), chymotrypsin and endoproteinase GluC, were used in parallel. In the generated peptide mixtures, Nt-peptides are thus N-terminally acetylated (whereas internal peptides are not) and enriched by COFRADIC prior to LC-MS/MS analysis. *In vivo* acetylated peptides will be further referred to as co-translationally modified peptides, while *in vitro* acetylated peptides will be referred to as N-terminally blocked peptides (see **Figure 1B**). The LC-MS/MS data were searched in the above described database, leading to the identification of 10,147, 6,796 and 5,373 unique peptide sequences in the samples digested by trypsin, chymotrypsin and endoproteinase GluC, respectively.

To evaluate the identification gain from using three proteases (see **Figure S1**), we calculated the overlap of the identified unique Nt-peptides and proteins when using each protease. Here, we need to consider that different proteases may generate Nt-peptides with identical start positions yet with different end positions. Therefore, we coupled a peptide start position to its protein entry in the database, creating a unique proxy for a protein’s N-terminus. However, the identified internal and C-terminal peptides may cause an overestimation of protease-specific start positions. Therefore, we applied a rule-based selection strategy to first remove internal and C-terminal peptides and to withhold a list of confidently identified and distinct N-termini per sample. These data were submitted to BioVenn (62) to visualize the overlap (**Figure 5**) and a list of peptides and proteins corresponding to each compartment was generated (**Supplementary table S1**).

**Figure 5:**
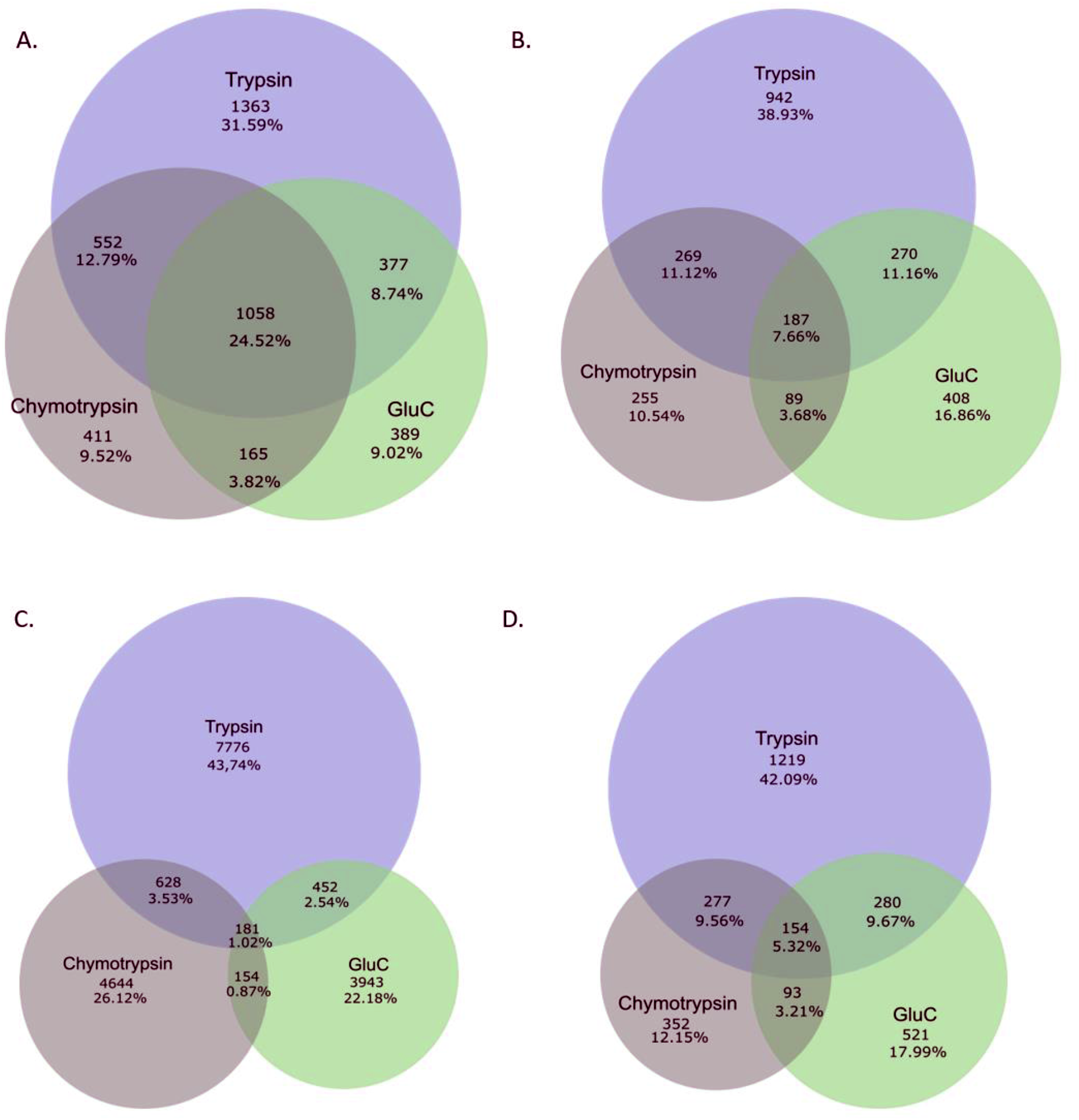
Venn diagrams (generated by http://www.biovenn.nl/) showing the overlap between the samples generated by the different proteases used. Overlap between the different samples based on distinct protein accession (after accession sorting), A) before the selection strategy and B) after the selection strategy, found in each sample (both absolute numbers and fractions are shown). Overlap between the different samples based on distinct peptides found in each sample C) before the filtering strategy (based on all identifications) and D) after the filtering strategy (thus N-terminal peptides).

In total, 2,896 distinct N-termini and 2,420 distinct proteins were identified upon combining the data from the different protease setups. About half of all proteins were identified in at least two datasets (**Figures 5A** and **5B**), this figure drops to 27% when considering matching peptide start sites (**Figures 5C** and **5D**). This drop is explained by different Nt-peptides generated from different Nt-proteoforms that have to the same protein database entry. Trypsin seems to account for the largest part of identified proteins and Nt-peptides (with 38.9 and 42.1% of unique identifications respectively), with chymotrypsin and GluC contributing additional unique sets of proteins (10.5 and 16.9%, respectively) and Nt-peptides (12 and 18%, respectively). Thus, the proteome coverage is indeed increased by using different proteases as each contributes with its own unique set of identified peptides, even exceeding our theoretical predictions (**Figure S1).**

We also evaluated how efficient COFRADIC enriched for N-terminal peptides (**Table 2**). Whereas Nt-peptides normally only account for less than 5% (based on shotgun proteomics data) of all peptides, we found that, following enrichment, 59.2% of the peptides generated by trypsin are Nt-peptides, which agrees with previous reports (36). COFRADIC was most efficient when using trypsin (73.2% of all peptides were N-terminal peptides, when including both protein Nt-peptides and pyroglutamate-starting peptides which are co-enriched by COFRADIC. However, the efficiency of enriching chymotryptic Nt-peptides is much lower (43.6%); one possible explanation for this is due to the fact that this protease recognizes more residues, thus generates a larger pool of (shorter) peptides by which the actual chemical modification step used for sorting in COFRADIC becomes less efficient. In addition, chymotryptic peptides are less basic than tryptic peptides, which might negatively influence their ionization (and thus detection).

**Table 2:** Overview of the numbers of distinct peptide sequences found in the different samples and N-terminal enrichment efficiencies. LC-MS/MS data were searched using Mascot and our custom-build database, and all identifications were stored and retrieved via ms_lims (at a confidence interval of 0.01) (51). Identifications were grouped by peptide sequence. When the same peptide sequence was found several times with different N-terminal modifications, priority was given to in vivo acetylated peptides. The enrichment efficiency was calculated by taking the sum of all peptides expected to be enriched by COFRADIC (these being Nt-peptides and pyroglutamyl-starting peptides) and dividing this by the total number of identified peptides. *with ArgC specificity.

|  | Trypsin* digested sample | Chymotrypsin digested sample | GluC digested sample |
| --- | --- | --- | --- |
| Free N-terminal peptides | 3,820 | 1,332 | 1,644 |
| Co-translationally modified peptides | 2,192 | 1,133 | 1,208 |
| Pyroglutamyl-starting peptides | 1,418 | 495 | 467 |
| Internal peptides | 2,717 | 3,836 | 2,054 |
| **Total** | **10,147** | **6,796** | **5,373** |
| **Fraction of N-terminal peptides** | **59.2%** | **36.3%** | **53.1%** |
| **Enrichment efficiency** | **73.2%** | **43.6%** | **61.8%** |

### 5. Stringent selection of N-terminal peptides

All identified peptides were loaded into a KNIME selection pipeline to select with high stringency Nt-peptides of both known and novel proteins/proteoforms. Our selection was based on the co-translational nature of protein Nt-acetylation, also considering the (possible) removal of the initiator methionine by methionine aminopeptidases, with extra translational evidence provided by Ribo-seq. Our strategy is outlined in **Figure 6** and explained in more detail below. It was applied on the dataset for each used protease, and all results were merged afterwards. In this way, peptides matched to a NTR accession could be traced back to any of the three proteases (**Table 3** and **Figure 7**). More information about the NTR peptides and the proteins retained in each step can be found in **Supplementary table S2**.

**Figure 6:**
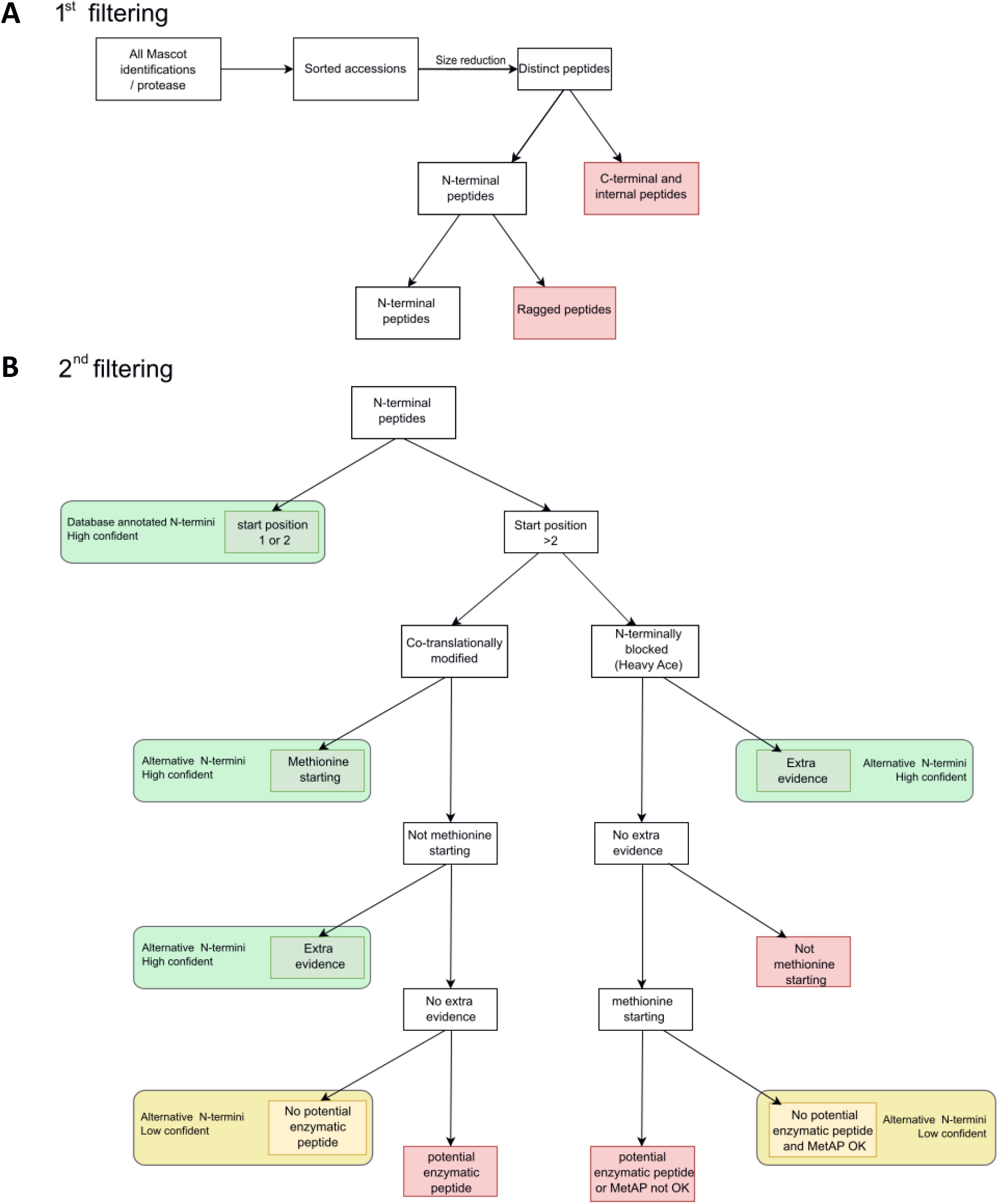
Schematic overview of the selection strategy, which can be divided in two main steps. A) Internal and C-terminal peptides are filtered out in the first step to retain the distinct Nt-peptides, while in the second step (B), the Nt-peptides segregate into categories, being either database annotated N-termini or alternative N-termini pointing to Nt-proteoforms, with a confidence level assigned to them. In the first step, the random picking of protein accessions by Mascot is solved by sorting accessions. This is followed by a reduction of the size of the dataset by removing duplicate sequences to obtain a list of distinct peptides. Next, C-terminal, internal and ragged peptides are removed to obtain a list of unique Nt-peptides, which forms the input for the second step in which Nt-peptides segregate into database annotated N-terminal and alternative N-terminal peptides. To evaluate if a peptide truly points to an alternative N-terminus generated by translation rather than by processing, a stringent selection strategy was used based on co-translational acetylation rules, the presence of an initiator methionine (and its processing) and the absence of a proteolytic cleavage site.

**Table 3:**
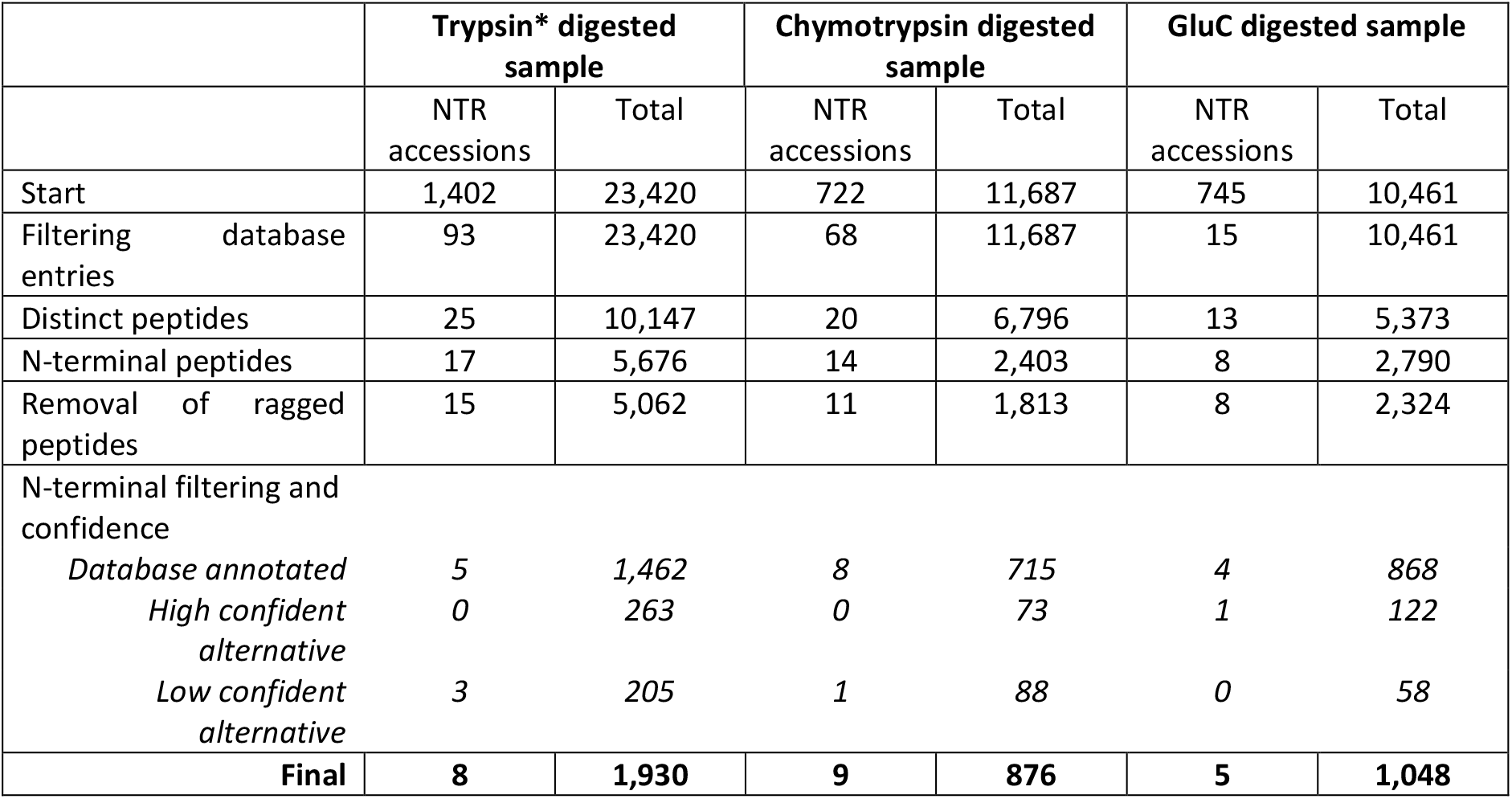
Numbers of peptides with accessions indicating that the peptide originates from an NTR protein during the different steps of the selection procedure. Note that in some steps the total dataset is also reduced in size (reported in the “Total” column). *with ArgC specificity.

**Figure 7:**
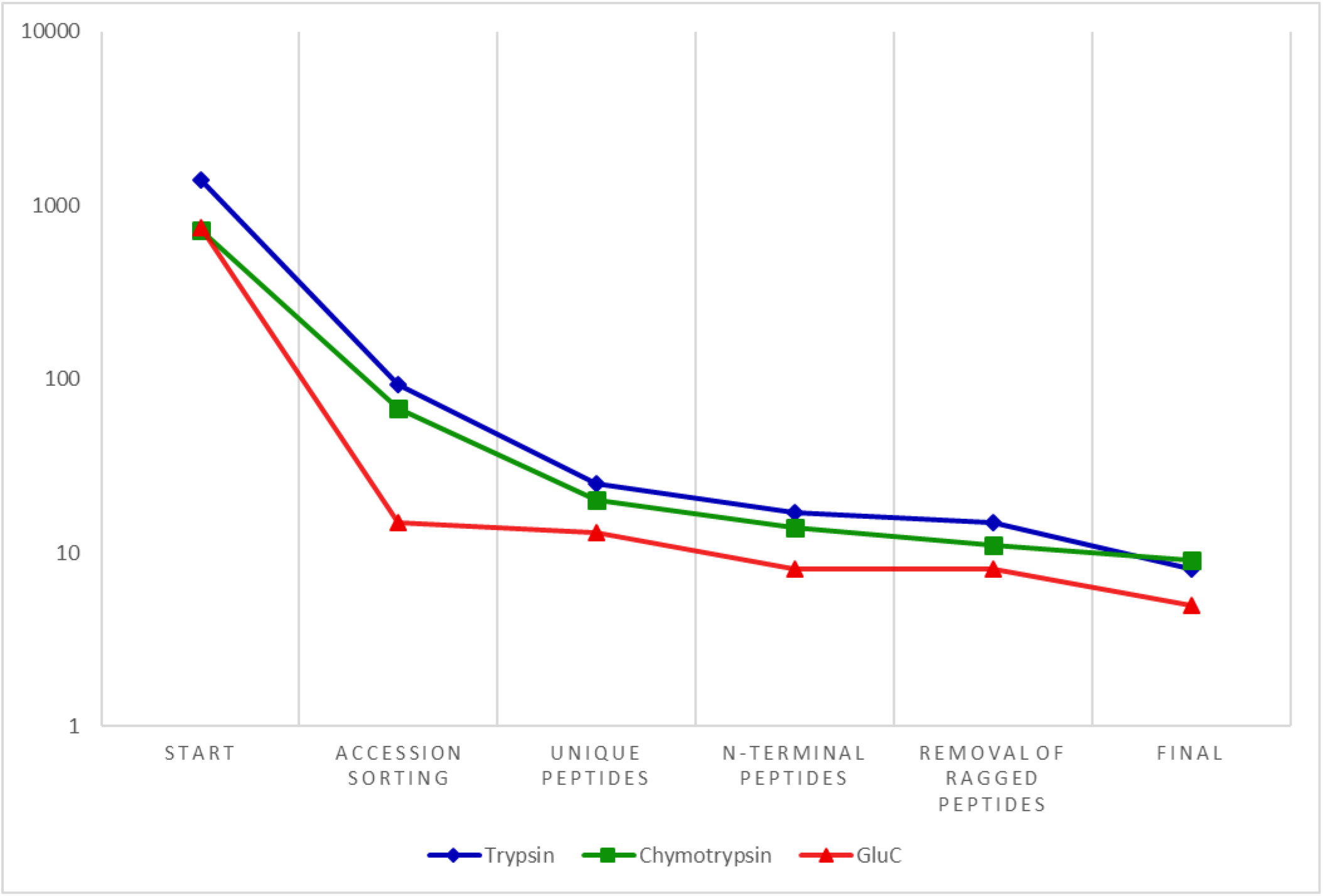
Plot showing the reduction in peptides matched to a NTR accession during the selection strategy, with the number of NTR matches shown on the y-axis (in a logarithmic scale).

#### 5.1 Matching peptides to proteoforms

The assignment of peptides to proteins (protein inference (63)) using protein sequence databases with a high level of subsequence redundancy, such as the database we have used, is challenging. In general, of all the identified peptides, only 3,246 (13.9%), 1,731 (14.8%) and 1,530 (14.6%) (for trypsin, chymotrypsin and GluC respectively) were unique. Peptides often matched to multiple proteins, both annotated and novel, with database entries to be chosen rather at random by Mascot upon identification of such peptides. This is because Mascot needed to be used at the peptide level and not at the protein level where the protein inference problem is better dealt with. To correct for this, we re-ordered all peptide-associated protein entries, prioritizing UniProt entries over UniProt isoform entries, and these over Ensembl entries (coming from the Ribo-seq data) as the UniProt database is by far the most completely annotated and curated of these databases. As for the Ensembl entries, we prioritized the different biotypes as aTIS > CDS > 5’ UTR > 3’UTR > NTR, thus again prioritizing for the most confident or plausible origin of the peptide. Based on these criteria, we assigned each peptide with one main protein entry and listed all other entries in a separate column (the isoform column) as these can contain extra information, such as Ribo-seq evidence for a translation initiation event at a matching position in a transcript. Further, when a peptide was matched to several entries of the same category/biotype, we prioritize the entry holding the lowest start position, thus favoring matches at a protein’s utmost N-terminal position. Finally, for any unresolved cases, we sorted database entries alphabetically, which was the case for 2,299, 1,250 and 1,204 peptides in the trypsin, chymotrypsin and GluC datasets, respectively.

This set of rules ensures that when a peptide was found by more than protease, it was always matched to the same protein entry, simplifying the final merging of the data. In addition, these rules also imply that many peptides that were initially associated with NTR proteins were reassigned as we prioritized matches to known, annotated proteoforms and protein-coding regions. Indeed, before filtering, about 6% of all identifications were matched to an NTR entry (regardless of the protease used). After filtering, only very few NTR entries (0.40%, 0.58% and 0.14% for trypsin, chymotrypsin and GluC, respectively) remained (**Table 3**). Additionally, a huge fraction of the identified NTR peptides were reassigned to UniProt (isoforms), (93.37%, 91.10% and 97.99% for trypsin, chymotrypsin and GluC, respectively). For example the co-translationally acetylated peptide AVNVYSTSVTSDNLSR, identified in the trypsin sample matched to ENST00000522077_8_135625981_ntr_100db1. However, this is also the mature Nt-peptide of a UniProt entry, Q15691 (Microtubule-associated protein RP/EB family member 1) and was therefore matched to this entry.

While checking the influence of the database entry filtering on the overall distribution of the database entries, we noticed that the fraction of UniProt proteins, UniProt isoforms and the different Ribo-seq categories were equally distributed over the three datasets (before and after filtering), and that after database entry filtering, more proteins were matched to UniProt entries (**Supplementary figure S3**). For all remaining NTR entries (just 176 in total), 94 (53.4%) of these were found to be unique. A list of all NTR entries and peptides before and after this first filtering step is shown in **Supplementary table S2**.

#### 5.2 Deduplication and removal of non-N-terminal peptides

In the dataset, now consisting of all identified peptides filtered for database entries, some peptides were present several times as, for instance, they were identified in different fractions of a certain sample (for example: KSAPSTGGVKKPH found in fraction B6 and B9 of the chymotrypsin sample). In the second filtering step, we removed such peptide sequence duplicates. Of note, some peptides were found with different N-terminal modifications. For example, GDVVPKDANAAIATIKTKR (ENST00000530835_11_90283408_ntr_011db2), identified in the trypsin dataset was found both with a free N-terminus and with a co-translationally acetylated one. Peptides were grouped according to their sequence and co-translational acetylation was prioritized over N-terminally blocked, over unmodified (free α-amine group), thereby giving more weight to (translational) evidence that an identified peptide indeed points to the N-terminus of a translation product. When a peptide was identified several times with the same Nt-modification, the highest scoring peptide-to-spectrum match (PSM) was retained. How many times a peptide was identified, the degree of co-translational acetylation and the different Nt-modifications the peptide was identified with (peptideforms) are listed. Thus, for the example above, both modifications (*in vivo* and *in vitro* acetylation) are reported and the acetylation percentage was calculated, being 71% (based on PSM counts). This filtering step reduced the dataset from 23,420, 11,687 and 10,461, to 10,147, 6,796 and 5,373 unique peptide sequences for trypsin, chymotrypsin and GluC, respectively. NTR peptides were also found several times or with different Nt-modifications (see example above), thus the number of NTR matches further reduced to 25, 20 and 13 for trypsin, chymotrypsin and GluC respectively.

Next, from this list of unique peptides, we focused on the Nt-peptides and excluded co-enriched internal and C-terminal peptides. Both peptide classes can be distinguished from Nt-peptides as Nt-peptides are acetylated, either *in vivo* or *in vitro* and internal peptides contain an unmodified N-terminus (free α-amine), whereas the C-terminal peptides are not followed by an amino acid. This allows us to further reduce our dataset to 5,876, 2,403 and 2,790 distinct Nt-peptides for trypsin, chymotrypsin and GluC, respectively.

Some identified NTR peptides were internal or C-terminal peptides as the number of NTR matches further dropped to 16, 13 and 8 for trypsin, chymotrypsin and GluC, respectively. However, as NTR proteins are generally short (**Figure 3B**), we also retained peptides that are both N- and C-terminal and thus cover the complete sequence of the (micro)protein. We identified one such N-terminally blocked peptide in two different datasets (trypsin and chymotrypsin) that covers the complete protein sequence: MKEETKEDAEEKQ. This peptide was found to be a unique peptide belonging to ENST00000486575_22_20127011_ntr_100db1. Note that most of the NTR proteins that were removed in this step were identified by peptides that were neither co-translationally acetylated nor N-terminally blocked. Therefore, they did not contain evidence to be further considered as Nt-peptides.

Among all distinct Nt-peptides, we noticed that many peptides were found to be C-terminally ragged, an artefact previously observed in COFRADIC datasets. Such ragged peptides share the same start position and are linked to the same database entry but are C-terminally shorter. One example is ENST00000556323_14_92026617_ntr_100db1 for which four different peptides were found: EKKEVVEEAENGRDAPAD, EKKEVVEEAENGRDAP, EKKEVVEEAENGR and EKKEVVEEAEN. These peptides were identified in the chymotrypsin and trypsin dataset, but note that the C-terminal ends of the peptides do not comply with chymotrypsin’s specificity for cleavage (Y, W, F, L or M). We grouped such peptides based on their start site and the coupled database entry, and again prioritized for co-translational acetylation. If such peptides held the same Nt-modification, the longest peptide sequence was kept. However, information on all shorter variants was also stored. This step further reduced the dataset and the number of NTR accessions found. For trypsin, chymotrypsin and GluC we now ended with 5,063, 1,813 and 2,324 unique Nt-peptides respectively, and only 15, 11 and 8 Nt-peptides remained matched to a NTR entry.

#### 5.3 Further filtering of N-terminal peptides based on co-translational modifications

The list of distinct Nt-peptides was further filtered for Nt-peptides that are proxies for translation by removing peptides that reported protein processing and peptides that were very unlikely to be Nt-peptides (as explained below and schematically summarized in **Figure 6B**). For all remaining peptides, a confidence level (high or low) and a category, database-annotated or alternative N-termini, were assigned. The latter points to Nt-peptides originating from a (N-terminal) variant of a canonical protein. In this way, a high confident Nt-peptide originating from an NTR protein, combined with the Ribo-seq data that were used to build the database, must provide solid evidence that a transcript from a presumed non-translated region can be translated.

##### 5.3.1 Database-annotated protein N-termini

When a Nt-peptide matched to a protein position one or two (following initiator methionine removal), it was considered a highly confident, database-annotated Nt-peptide. Such peptides may match to a regular UniProt protein, but also to a UniProt isoform or an Ensembl entry (such as an NTR entry). 1,462, 715 and 868 database-annotated N-termini were found for respectively trypsin, chymotrypsin and GluC, respectively. From the 15, 11 and 8 NTR accessions in the trypsin, chymotrypsin and GluC datasets, respectively 5, 8 and 4 of the Nt-peptides were listed under this category.

##### 5.3.2 Alternative protein N-termini

We further filtered Nt-peptides that did not start at positions one or two as these could also originate from signal or transit peptide removal and/or other proteolytic activities in cells or whilst preparing the samples. For trypsin, chymotrypsin and GluC there were respectively 3,600, 1,098 and 1,455 presumed Nt-peptides that started beyond position two. We differentiated between (*in vivo*) co-translationally acetylated peptides and *in vitro* blocked peptides. The former holds more evidence that they originated from translation events, while the latter could also point to protein processing. The majority of N-termini with a start position beyond two were found *in vitro* blocked. For the three proteases, 422, 113 and 214 co-translationally acetylated peptides, and 3,178, 985 and 1,241 blocked peptides were identified for trypsin, chymotrypsin and GluC, respectively. To further check if such peptides originated from translation events we used extra evidence from Ribo-seq and evaluated the presence of an initiator methionine.

###### 5.3.2.1 Co-translationally acetylated peptides

If a peptide was co-translationally acetylated and either started with or was preceded by a methionine, it was classified as a highly confident alternative Nt-peptide. 222, 56 and 109 Nt-peptides belong to this category for trypsin, chymotrypsin and GluC, respectively. Only one peptide matching to an NTR entry was classified as such: MASAASSSSLE (found in the GluC dataset) matching to position 12 of ENST00000403258_6_88276364_ntr_101db1.

Co-translationally acetylated peptides that do not start with nor are preceded by a methionine were also identified (200, 57 and 105 for trypsin, chymotrypsin and GluC, respectively). For such peptides, we first searched for extra evidence (from UniProt isoforms or Ribo-seq) that could indicate that these peptides pointed to alternative starts of translation. For respectively trypsin, chymotrypsin and GluC, we only found three, two and one peptide with extra Ribo-seq evidence and categorized these peptides as highly confident alternative Nt-peptides. Of note, among these peptides, none were from NTR entries. For the remaining peptides without extra Ribo-seq evidence and not starting with methionine we verified if the preceding amino acid was a potential recognition site of the protease used for digestion. If not, such peptides were considered as low confident alternative Nt-peptides, all other peptides were removed.

A missing initiator methionine can be explained by a non-AUG codon that was used for initiating translation, which are generally also translated into a methionine (64). However, in protein databases such as UniProt, non-AUG start codons will be indicated as translated into the amino acid they encode for instead of methionine. As such, when using protein databases, one would not be able to identify the corresponding Nt-peptide. 106, 24 and 38 peptides were found in this category, with three, one and zero Nt-peptides matching to a NTR entry in the trypsin, chymotrypsin and GluC datasets respectively.

###### 5.3.2.2 N-terminally blocked peptides

For the Nt-blocked peptides (3,178, 985 and 1,241 peptides for trypsin, chymotrypsin and GluC, respectively) extra evidence for translation was first evaluated (see above). UniProt isoforms or Ribo-seq corroborated peptides were considered highly confident alternative Nt-peptides. As such 38, 15 and 12 Nt-blocked peptides were assigned as highly confident (for trypsin, chymotrypsin and GluC, respectively). Stringent filtering was then applied for peptides lacking extra evidence. The presence of a methionine at the peptide’s start may provide translational evidence, therefore we scanned if the starting or preceding amino acid was methionine, and if this was not the case, peptides were removed. One example is AVGVIKAVDKKAAGAGKVT, starting at position 27 of ENST00000415278_1_96448151_ntr_010db2. It was found in the chymotrypsin dataset and is not initiated with a methionine. It is thus unlikely that this peptide points to a translation event and it was therefore removed. Note that this step removed the majority of presumed N-terminal peptides (3,009, 886 and 1,201 in the trypsin, chymotrypsin and GluC datasets, respectively).

Peptides that survived this filtering step were further evaluated by checking if the preceding amino acid was a potential cleavage site for the protease used (as some unwanted trans-acetylation acitivity is possible (65)) and if initiator methionine (processing) agreed with the specificity of the MetAPs. Peptides that met these criteria were retained as low confident alternative Nt-peptides, leading to 102, 65 and 21 of such peptides in the trypsin, chymotrypsin and GluC datasets respectively.

Clearly, with this stringent filtering strategy, a large part of peptides that do not confidently point to a protein’s N-terminus, but rather to processing events were removed. In total, we find 263, 73 and 122 highly confident alternative N-termini and 205, 88 and 58 low confident alternative N-termini (for trypsin, chymotrypsin and GluC respectively).

Finally, 1,930, 876 and 1,048 Nt-peptides were identified by COFRADIC in the cytosolic HEK293T proteome in respectively the trypsin, chymotrypsin or GluC digested samples (thus 3,854 in total). Of the peptides that matched to an NTR entry, 8, 9 and 5 peptides (for trypsin, chymotrypsin and GluC, respectively) survived the different filtering steps, thus 22 in total, which is merely 0.57% of all identified Nt-peptides. The majority (17) start at position one or two of the corresponding NTR protein and are thus listed in the highly confident category. For trypsin, the remaining three NTR matches are low confident alternative N-terminal peptides (co-translationally modified, but not starting with methionine). The same is observed for one peptide in the chymotrypsin dataset, while in the GluC dataset, we detected a co-translationally acetylated peptide that starts with methionine and is thus a highly confident alternative N-terminus (**Supplementary table S2**).

##### 5.3.3 Final merging of the data

In the last step, the three different datasets were concatenated. As these proteases generated different ends at a given protein N-terminus, merging based on peptide sequence was not possible. Therefore, we relied on the database entry and the peptide’s start position and retained the longest peptide sequence as this contained the most information, but, as indicated above, also kept all information on shorter forms of this peptide. We also listed the datasets in which a peptide was identified and re-calculated the degree of *in vivo* acetylation based on all identified peptides. This resulted in a final dataset of 2,896 unique and confident Nt-peptides (**Supplementary table S3)**, 19 of which that matched to an NTR entry (**Table 4**).

**Table 4:** List of Nt-peptides matched to an NTR entry. This table list all N-terminal peptides that were identified and matched to an NTR protein. For each peptide, the accession (a detailed explanation of the information contained in the accessions from ENSEMBL can be found in the materials and methods section), the start and end positions, the sequence, the enzyme (this indicates by which protease the peptide was detected), modification (“Mod.”, which indicates the modification found on the protein’s N-terminus), the confidence level assigned to the peptides (“Conf.”, which is either high or low (see main text)), isoform column, transcript information, protein length, “Prec. AA”, which indicates the preceding amino acid, and the spectral counts (#) are listed.

| Accession | Start End | Sequence | Enzyme | Mod. | Conf. | Isoforms | Transcript info | Protein Length | Prec. AA | # |
| --- | --- | --- | --- | --- | --- | --- | --- | --- | --- | --- |
| ENST00000543961_12_25803373_ntr_100db1 | 1-36 | MAEAPNMAVVNEQ<br>QMPEEVPAPAPAE<br>PVQEAPKGR | Chymo | Heavy acetyl | high | ENST00000544060_12_103976970_ntr_100db1 (1-36) | Thymine-DNA glycosylase pseudogene 1, processed pseudogene. | 82 |  | 1 |
| ENST00000439303_10_10174230_ntr_100db1 | 1-33 | MDGEEKTCGGCEGP<br>DAMYVKLISSDGHEFI<br>VKR | Trypsin | Heavy acetyl | high |  | Elongin C pseudogene 3, processed pseudogene | 115 |  | 2 |
| ENST00000486575_22_20127011_ntr_100db1 | 1-13 | MKEETKEDAEKQ | Chymo, Trypsin | Heavy acetyl | high |  | RAN binding protein 1, Retained intron | 13 |  | 2 |
| ENST00000458332_17_19446852_ntr_111db1 | 2-29 | ADDAGAAGGPGGPG<br>GPEMGNRGGFRGGF | GluC | Co-transl. acetyl | high |  | Ribosomal protein S2 pseudogene 46, processed pseudogene | 88 | M | 1 |
| ENST00000454707_6_5972475_ntr_100db1 | 2-9 | ADTFLEHM | Trypsin | Co-transl. acetyl | high | ENST00000564276_15_72219034_ntr_001db3 (2-9) | Pyruvate kinase M1/2 pseudogene 5, processed pseudogene | 157 | M | 2 |
| ENST00000216019<br>_22_38506168_ntr<br>_100db1 | 2-37 | ASATGDSASERESAA<br>PAAAPTAEAPPSVV<br>TRPEPQ | Chymo | Co-transl.<br>acetyl | high |  | DEAD-box<br>helicase 17,<br>retained intron | 463 | M | 3 |
| ENST00000556323<br>_14_92026617_ntr<br>_100db1 | 2-19 | EKKEVVEEAENGRDA<br>PAD | Chymo,<br>Trypsin | Heavy<br>acetyl | high |  | Prothymosin<br>alpha<br>pseudogene,<br>processed<br>pseudogene | 56 | M | 13 |
| ENST00000498385<br>_22_23894847_ntr<br>_001db3 | 2-16 | HSIGKIGGAQNRSYS | GluC | Heavy<br>acetyl | high |  | Macrophage<br>migration<br>inhibitory<br>factor, retained<br>intron | 54 | M | 2 |
| ENST00000415278<br>_1_96448241_ntr<br>_001db3 | 2-13 | KAVDKKAAGAGK | GluC | Heavy<br>acetyl | high |  | Eukaryotic<br>translation<br>elongation<br>factor 1 alpha 1<br>pseudogene 11,<br>processed<br>pseudogene | 25 | M | 1 |
| ENST00000586518<br>_17_75779096_ntr<br>_010db2 | 2-14 | KSAPSTGGVKKPH | Chymo | Heavy<br>acetyl | high |  | H3.3 histone B,<br>retained intron | 17 | M | 2 |
| ENST00000478033<br>_1_159918889_ntr<br>_100db1 | 2-23 | NVIGLQMGTRNGAS<br>QAGMTGYG | GluC | Co-transl.<br>acetyl | high |  | Transgelin 2,<br>processed<br>transcript | 29 | M | 1 |
| ENST00000569492<br>_16_35803171_ntr<br>_100db1 | 2-21 | RKAEGDAKGDKAKVK<br>DEPQR | Chymo | Heavy<br>acetyl | high |  | High mobility<br>group<br>nucleosomal<br>binding domain<br>2 pseudogene<br>41, processed<br>pseudogene | 81 | M | 1 |
| ENST00000555320_14_80822530_ntr_010db2 | 2-17 | RKAEGDAKGDKAKVK D | Chymo | Heavy acetyl | high | ENST00000569492_16_35803171_ntr_100db1 (2-17) | High mobility group nucleosomal binding domain 2 pseudogene, processed pseudogene | 88 | M | 1 |
| ENST00000556323_14_92026566_ntr_100db1 | 2-36 | SDAAVDTSEITTKDL KEKKEVVVEEAENGRD APAD | Chymo, Trypsin | Co-transl. acetyl | high |  | Prothymosin alpha pseudogene, processed pseudogene | 73 | M | 43 |
| ENST00000582213_17_7572835_ntr_100db1 | 9-31 | FPSNWNEIVDSFDD MNLSesLLR | Trypsin | Co-transl. acetyl | low |  | Eukaryotic translation initiation factor 4A1, retained intron | 148 | G | 1 |
| ENST00000403258_6_88276364_ntr_101db1 | 12-22 | MASAASSSSLE | GluC | Co-transl. acetyl | high |  | ACTB pseudogene 8, processed pseudogene | 146 | E | 1 |
| ENST00000402643_6_166064725_ntr_111db1 | 64-83 | ASTGTAKAVGKVIPEL NGKL | Chymo | Co-transl. acetyl | low | ENST00000402643_6_166064722_ntr_001db3 (65-84)<br>ENST00000402643_6_166064704_ntr_100db1 (71-90)<br>ENST00000402643_6_166064650_ntr_110db1 (89-108)<br>ENST00000402643_6_166064509_ntr_010db2 (136-155) | Glyceraldehyde-3-phosphate dehydrogenase pseudogene 72, Transcribed processed pseudogene | 125 | P | 3 |
| ENST00000530835_11_90283408_ntr_011db2 | 70-88 | GDVVPKDANAAIATIKTKR | Trypsin | Co-transl. acetyl | low | ENST00000530835_11_90283141_ntr_001db3 (159-177)<br>ENST00000530835_11_90283060_ntr_001db3 (186-204)<br>ENST00000530835_11_90283018_ntr_001db3 (200-218)<br>ENST00000530835_11_90283009_ntr_101db1 (203-221)<br>ENST00000530835_11_90283006_ntr_010db2 (204-222)<br>ENST00000530835_11_90282775_ntr_001db3 (281-299)<br>ENST00000530835_11_90282763_ntr_010db2 (285-303)<br>ENST00000530835_11_90282676_ntr_100db1 (314-332) | Tubulin alpha pseudogene 2, processed pseudogene | 124 | H | 7 |
| ENST00000530835_11_90283408_ntr_011db2 | 74<br>88 | PKDANAAIATIKTKR | Trypsin | Co-transl. acetyl | low | ENST00000530835_11_90283141_ntr_001db3 (163-177)<br>ENST00000530835_11_90283060_ntr_001db3 (190-204)<br>ENST00000530835_11_90283018_ntr_001db3 (204-218)<br>ENST00000530835_11_90283009_ntr_101db1 (207-221)<br>ENST00000530835_11_90283006_ntr_010db2 (208-222)<br>ENST00000530835_11_90282775_ntr_001db3 (285-299)<br>ENST00000530835_11_90282763_ntr_010db2 (289-303)<br>ENST00000530835_11_90282676_ntr_100db1 (318-332) | Tubulin alpha pseudogene 2, processed pseudogene | 124 | F | 7 |

Several interesting observations can be made for the identified NTR peptides. For example, 12 of these 19 peptides do not end on an amino acid that corresponds to a cleavage site of the protease used. For seven of them, initiator methionine processing did not seem to follow the known MetAPs rules, instead, peptides were identified starting with E, H, K, N or R for which the initiator methionine is normally not removed. 13 peptides are only identified by one or just two PSMs, which points to the low abundance of NTR proteins. On the other hand, the majority of the identified NTR peptides are unique, supporting their NTR origin. For Tubulin alpha pseudogene 2, two peptides were identified. However, one peptide is a shorter variant of the other missing the first four amino acids. Finally, concerning the biotypes of these transcripts in Ensembl, 12 are processed pseudogenes, five have a retained intron, one is a transcribed processed pseudogene and one a processed transcript.

### 6. Further curation of identified NTR proteins by BlastP analysis

After this stringent filtering, an additional curation step was performed, similar as described by Zhu *et al.* (66). The identified peptides from the NTR proteins were searched for homologous sequences using the BlastP algorithm (using standard settings, automatically adjusted for short sequences and an e-value of 200,000) against the human UniProtKB/Swiss-Prot database.

Strikingly, nine out of the 19 peptides had an exact match to a UniProt protein sequence (**Supplementary table 4**). This can be explained by the semi-protease settings (semi-ArgC, semi-GluC and semi-chymotrypsin) that were used during the database search, which imply that one end of the peptide (either C- or N-terminal, not both) was allowed not to comply with the protease’s specificity rules, and such settings are required to identify alternative start positions (37). However, in the cases explained below, the peptides’ ends do not comply with the specificity of the protease used, while they were matched to a start position of a NTR protein. As such, a match against NTR proteins appears “forced” over a match to an internal peptide of a UniProt protein. For example, the peptide ADTFLEHM is found in the trypsin digested sample. As the peptide does not end on R, its N-terminal amino acid should follow a trypsin (acting as ArgC) cleavage site or be a start position. The peptide was matched to the N-terminus (position 2, preceded by a methionine) of ENST00000454707 (pyruvate kinase M1/2 pseudogene 5, processed pseudogene). However, following BlastP analysis this peptide was found to match to a peptide starting at position 22 of pyruvate kinase PKM (UniProtKB accession P14618). Here, this peptide is preceded by a methionine, which is not a potential trypsin cleavage site but is likely an internal start site of P14618. Thus, similar to Zhu *et al.* (67), we assume that for all peptides with an exact match after BlastP analysis, the semi-setting caused them to match against an NTR protein, whereas they likely originated from an annotated protein.

Of note, by removing such peptides, we lost peptides of which the initiator methionine was removed though not in agreement with the specificity of the MetAPs. However, one case remains, being the removal of methionine leading to the EKKEVVEEAENGRDAPAD peptide.

The remaining ten out of 19 peptides have a strong homology match to a UniProt entry with just one (or two) amino acids that are different (**Table 5**). There are two hits for Prothymosin alpha and Tubulin alpha, this because for both cases, two peptides were identified that differ at their N-terminus but are linked to the same UniProt protein and contain the same single amino acid variation (SAAV). Interestingly, nine of the ten peptides just had a single amino acid difference with a UniProt reference protein, which for most of them could be explained by a potential SNP. For example, for ADDAGAAGGPGGPGGP**<u>E</u>**MGNRGGFRGGF (Ribosomal protein S2 pseudogene 46) and ADDAGAAGGPGGPGGP**<u>G</u>**MGNRGGFRGGF (40S ribosomal protein S2), both proteins were found co-translationally acetylated and for both, the peptides were matched to the second position in the protein sequence. In the nucleotide sequence of the UniProt protein, glycine is encoded by GGG, which only differs one base from the GAG codon that encodes glutamic acid in the pseudogene. For other peptides, any straightforward explanation is less clear. For example, EKKEVVEEAENGRDAPA**<u>D</u>** (protymosin alpha pseudogene) and EKKEVVEEAENGRDAPA**<u>N</u>** (prothymosin alpha). The peptide was identified as being heavy acetylated and, in the pseudogene, preceded by a methionine (i.e. a potential Nt-peptide). However, it is unlikely that the initiator methionine is removed when the second amino acid is glutamic acid (43,44). As for the UniProt protein, the peptide is preceded by lysine which is not a trypsin (here acting as ArgC) nor chymotrypsin cleavage site however, the lysine is encoded as AAG and is surrounded by a Kozak-like sequence, which could point to a non-AUG alternative translation start site. Nonetheless, both cases appear unlikely, and the actual difference (Asn versus Asp) can be explained by a SNP (GAC instead of AAC).

**Table 5:** List of all peptides that matched to an NTR entry and for which a strong homology match to a UniProt entry was found. For each peptide, the accessions (containing both the ENSEMBL accession and transcript information) are provided, the column “match” contains the peptide sequence of the identified peptide matched to the NTR and the corresponding sequence in the UniProt protein it was matched to, with the positions of the peptide within the NTR and UniProt proteins. The UniProt accession to which the NTR peptide is matched to, the enzyme (protease) the peptide was identified with and some extra information are listed. Abbreviations: SNP = single nucleotide polymorphism.

| Accession | Match | UniProt entry | Enzyme | Extra info |
| --- | --- | --- | --- | --- |
| ENST00000439303<br>_10_10174230_ntr_100db1<br>Elongin C pseudogene 3,<br>processed pseudogene | 1 MDGEEKTCGGCEGPDAMYVKLISSDGHEFIVKR 33<br>MDGEEKT GGCEGPDAMYVKLISSDGHEFIVKR<br>1 MDGEEKTYGGCEGPDAMYVKLISSDGHEFIVKR 33 | Q15369<br>Elongin-C | Trypsin | 1 AA difference<br>Possible SNP? Yes |
| ENST00000486575<br>_22_20127011_ntr_100db1<br>RAN binding protein 1,<br>Retained intron | 1 MKEETKEDAEKQ 13<br>KEETKEDAEKQ<br>189 VKEETKEDAEKQ 201 | P43487<br>Ran-specific GTPase-activating<br>protein (RANBP1) | Trypsin,<br>chymotrypsin | 1 AA difference<br>Possible SNP? No |
| ENST00000458332<br>_17_19446852_ntr_111db1<br>Ribosomal protein S2<br>pseudogene 46, processed<br>pseudogene | 2 ADDAGAAGGPGGPGGPEMGNRGGFRGGF 29<br>ADDAGAAGGPGGPGGP MGNRGGFRGGF<br>2 ADDAGAAGGPGGPGGPGMGNRGGFRGGF 29 | P15880<br>40S ribosomal protein S2 | GluC | 1 AA difference<br>Possible SNP? Yes |
| ENST00000556323<br>_14_92026617_ntr_100db1<br>Prothymosin alpha<br>pseudogene, processed<br>pseudogene | 2 EKKEVVVEEAENGRDAPAD 19<br>EKKEVVVEEAENGRDAPA+<br>19 EKKEVVVEEAENGRDAPAN 36 | P06454 Prothymosin alpha | Trypsin,<br>chymotrypsin | 1 AA difference<br>Possible SNP? Yes |
| ENST00000556323<br>_14_92026566_ntr_100db1 | 2 SDAAVDTSSEITTKDLKEKKEVVVEEAENGRDAPAD 36<br>SDAAVDTSSEITTKDLKEKKEVVVEEAENGRDAPA+<br>2 SDAAVDTSSEITTKDLKEKKEVVVEEAENGRDAPAN 36 | P06454 Prothymosin alpha | Trypsin,<br>chymotrypsin | 1 AA difference<br>Possible SNP? Yes |
| Prothymosin alpha<br>pseudogene, processed<br>pseudogene |  |  |  |  |
| ENST00000582213<br>_17_7572835_ntr_100db1<br>Eukaryotic translation<br>initiation factor 4A1, retained<br>intron | 9 FPSNWNEIVDSFDDMNLSESLLR 31<br>SNWNEIVDSFDDMNLSESLLR<br>23 IESNWNEIVDSFDDMNLSESLLR 45 | P60842 Eukaryotic initiation<br>factor 4A-I | Trypsin | 2 AA difference<br>Possible SNP? No |
| ENST00000403258<br>_6_88276364_ntr_101db1<br>ACTB pseudogene 8,<br>processed pseudogene | Match to Q6S8J3<br>12 MASAASSSSLE 22<br>MA AASSSSLE<br>927 MATAASSSSLE 937<br>Match to P68032 (and other actin variants)<br>12 MASAASSSSLE 22<br>MA AASSSSLE<br>229 MATAASSSSLE 239 | Q6S8J3 POTE ankyrin domain<br>family member E and several<br>Actin variants (P68032, P62736,<br>P68133, ...) | GluC | 1 AA difference<br>Possible SNP? No |
| ENST00000402643<br>_6_166064725_ntr_111db1<br>Glyceraldehyde-3-phosphate<br>dehydrogenase pseudogene<br>72, Transcribed processed<br>pseudogene | 2 ASTGTAKAVGKVIPELNGKL 37<br>ASTG AKAVGKVIPELNGKL<br>209 ASTGAAKAVGKVIPELNGKL 228 | P04406 Glyceraldehyde-3-phosphate dehydrogenase | Trypsin | 1 AA difference<br>Possible SNP? Yes |
| ENST00000530835<br>_11_90283408_ntr_011db2<br>Tubulin alpha pseudogene 2,<br>processed pseudogene | 70 GDVVPKDANAIAIATIKTKR 88<br>GDVVPKD NAAIATIKTKR<br>321 GDVVPKDVNAIAIATIKTKR 339 | Q71U36<br>Tubulin alpha-1A chain and<br>other tubulin alpha chains:<br>P68363 P0DPH7, Q6PEY2,<br>Q9BQE3 | Trypsin | 1 AA difference<br>Possible SNP? Yes |
| ENST00000530835<br>_11_90283408_ntr_011db2<br>Tubulin alpha pseudogene 2,<br>processed pseudogene | 74 PKDANAIAIATIKTKR 89<br>PKD NAAIATIKTKR<br>325 PKDVNAIAIATIKTKR 339 | Q71U36<br>Tubulin alpha-1A chain<br>P68363 Tubulin alpha-1B chain<br>and other tubulin alpha chains:<br>P0DPH7, Q6PEY2, Q9BQE3 | Trypsin | 1 AA difference<br>Possible SNP? Yes |

### 7. Inspection of MS/MS spectra

Misidentification of MS/MS spectra is a considerable threat in MS-driven proteomics. This is especially true when considering unexpected and hence unaccounted for modifications, which can yield isobaric peptides with similar fragmentation patterns. In a recent proteogenomics study that also applied a highly stringent workflow, peptides with single amino acid substitutions were removed (68). However, similarly to Zhu *et al*. (2018) we considered that inspection of MS/MS-spectra by experts facilitates differentiation between correct and incorrect single amino acid variants as called by database searching. In the abovementioned study, SpectrumAI was used to perform this task at a large scale (67). Here, the inspection was done manually as only ten spectra required examination. Upon inspection, two peptides (EKKEVVEEAENGRDAPAD and SDAAVDTSSEITTKDLKEKKEVVEEAENGRDAPAD), both pointing to the same NTR (prothymosin alpha), were removed, the other eight were evaluated correct and thus retained. For the removed peptides, the mass difference was just 1 Da, which equals the mass difference between the peptide from the NTR and UniProt protein (the difference is the last amino acid, D in the NTR protein and N in the UniProt protein). Besides this, the internal Asn-Gly motif makes the inspection of the spectra more difficult as this motif is prone to deamidation to Asp-Gly (69).

For all other NTR proteins, synthetic peptides were used to compare and validate the identified spectral matches. If for a peptide the same precursor ion (same m/z) was found, this ion was selected and the top ten fragment ions of the synthetic peptides were selected and compared with the ranking of the same fragment ions of the fragmented peptide ion that was identified in our COFRADIC samples. For MDGEEKTCGGCEGPDAMYVKLISSDGHEFIVKR we experienced issues with the synthesis of the synthetic peptide and cannot draw conclusions for this case. The ranking of the fragment ions of ASTGTAKAVGKVIPELNGKL identified in our sample was too different from the ranking of the synthetic peptide (**Figure S4**). Therefore, ASTGTAKAVGKVIPELNGKL linked to Glyceraldehyde-3-phosphate dehydrogenase pseudogene 72 (ENST00000402643_6_166064725_ntr_111db1) was removed, further reducing the amount of confident Nt-peptides of NTRs to seven. For all other NTR linked Nt-peptides the ranking of their fragment ions was (highly) similar between the peptides identified in our COFRADIC samples and the synthetic peptides, and therefore, these peptides were retained. For example, the fragment ions of MASAASSSSLE where highly comparable (**Figure 8**, all the others can be found in **Figure S4**).

**Figure 8:**
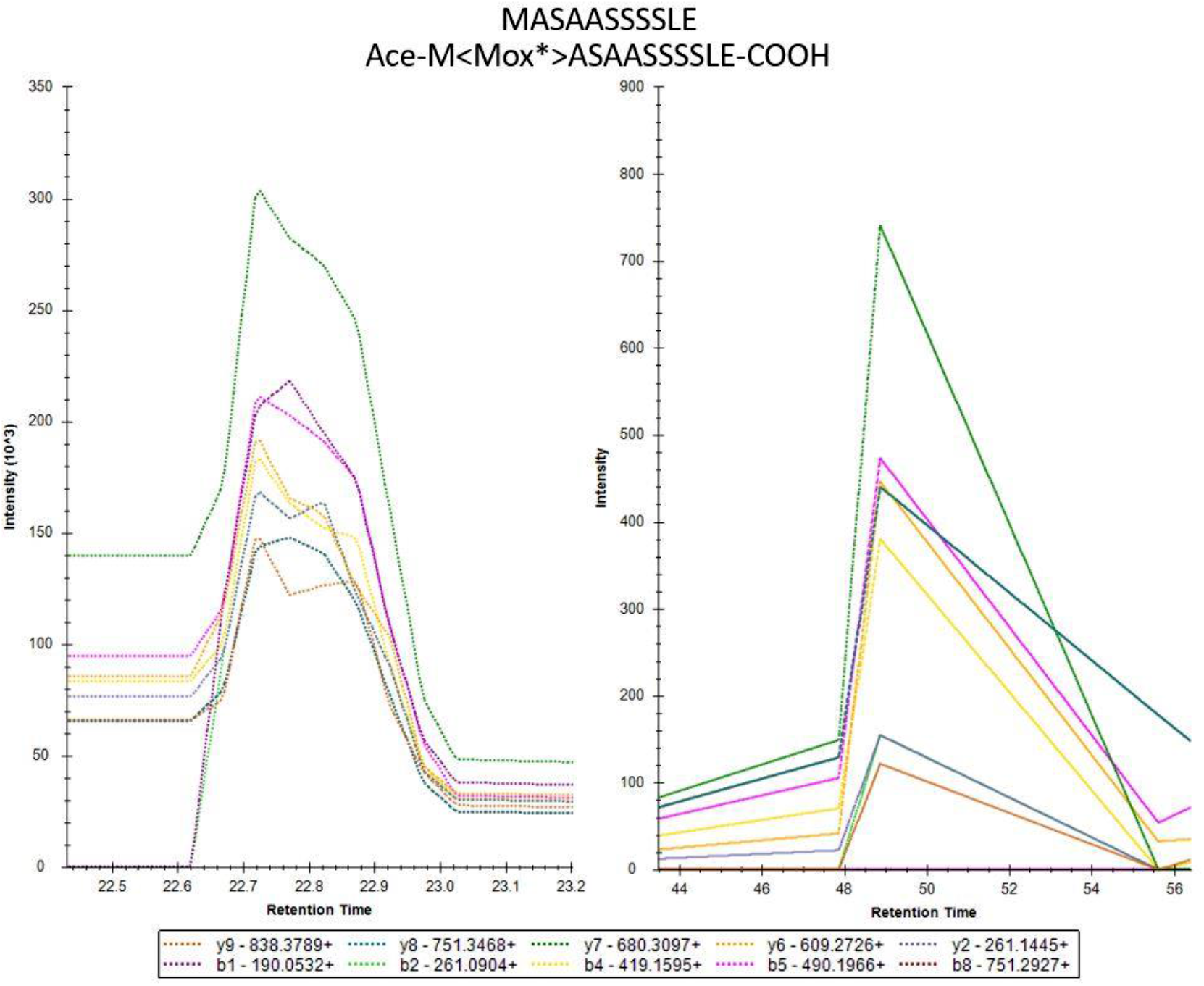
Comparison based on the ranking of the top 10 fragment ions of the synthetic peptide and the peptide identified in our COFRADIC samples of MASAASSSSLE. The modified peptide sequence is indicated at the top of the spectra. The top ten fragment ions (transitions), indicated with different colors at the bottom of the spectrum were used as comparison between the synthetic (left) and the identified (right) peptide.

### 8. Inspection of sequencing data

We used Ribo-seq data to determine splice boundaries and exact nucleotide sequences of exons at homologous genomic loci, to verify the expression of NTR-specific transcripts and variants in HEK293T cells. Using transcript coordinates of custom database entries, we developed an approach to map identified peptides to their genomic positions. Subsequently, we visualized peptides next to Ribo-seq sequencing reads using an Integrative Genome Viewer (IGV) (70). Inspection of the seven most-confident NTR peptides, lacking an exact BlastP match to UniProt entries and retained after inspection of MS/MS spectra (**Table 6**), revealed sequencing evidence supporting four NTRs (**Figure 9** and **Figure S5**). More specifically, we found NTR-specific, alternatively spliced reads from a retained intron transcript matching the FPSNWNEIVDSFDDMNLSESLLR peptide of Eukaryotic translation initiation factor 4A1 (EIF4A1) (**Figure 9**). Interestingly, translation of this peptide is initiated at exon 1 in a different reading frame compared to the canonical proteoform of the same gene, for which the N-terminus was also found. From exon 1, NTR continues translation directly to exon 3, thereby restoring the canonical reading frame. Splicing of RANBP1 NTR transcript responsible for the MKEETKEDAEEKQ peptide was also confirmed (**Figure S5**). From the remaining inspected cases, ribosome-protected fragments carrying NTR-specific non-synonymous and synonymous variant in peptide MDGEEKTCGGCEGPDAMYVKLISSDGHEFIVKR (ELOCP3), next to synonymous variants in ADDAGAAGGPGGPGGPEMGNRGGFRGGF (RPS2P46) and MASAASSSSLE (ACTBP8) were found (indicated in green, see **Figure S5**). However, many NTR-specific nucleotide variants were not supported. Instead, Ribo-seq reads at these positions were missing or mapped with a mismatch (indicated in red). To test if NTR peptides can be explained by known SNPs, NTR nucleotide sequences (encompassing the peptide) were compared to their closest annotated protein-coding match. None of the NTR-specific, non-synonymous sequence variations were previously reported in the canonical genes by the dbSNP database (build 142 hg38) (71).

**Table 6:**
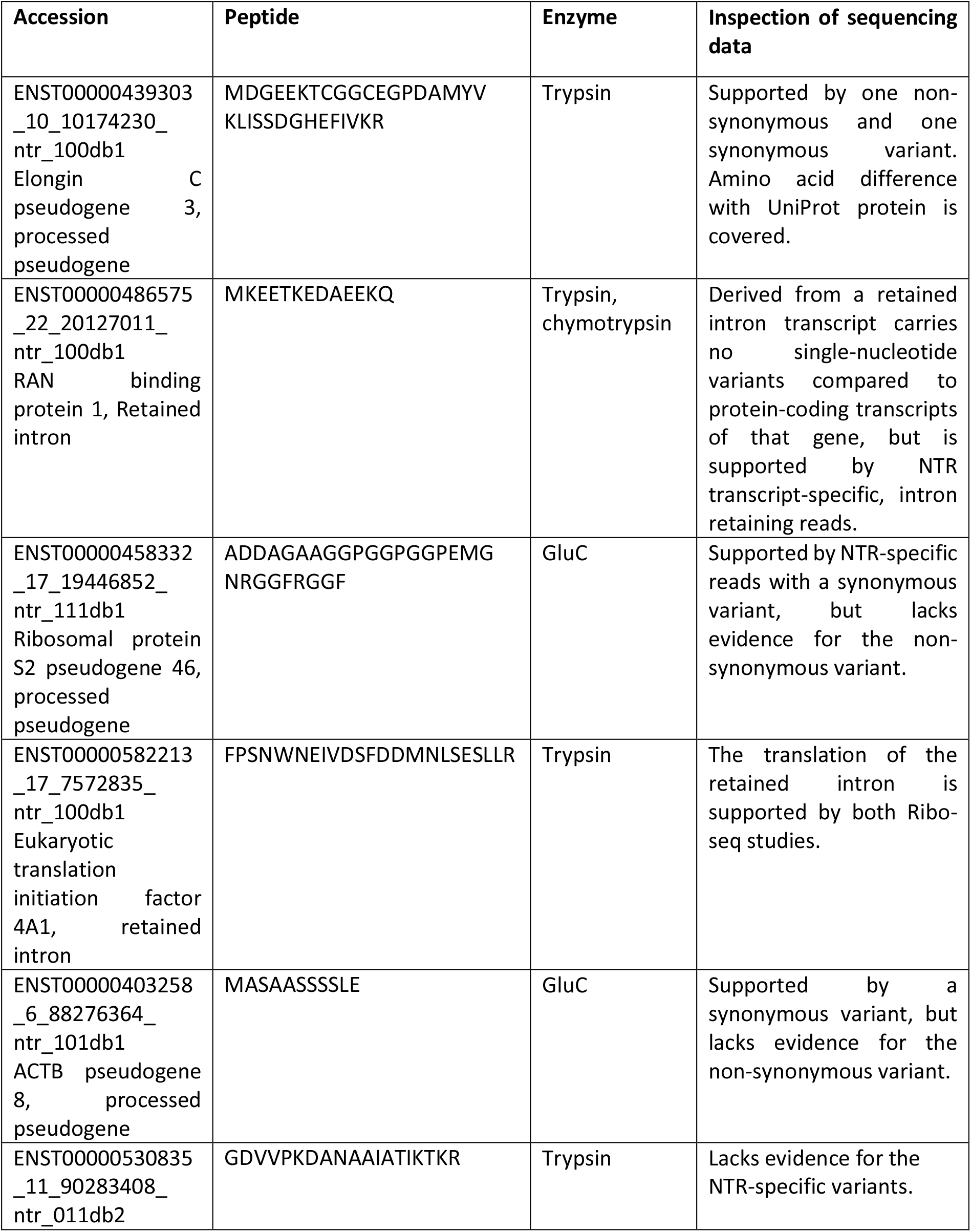
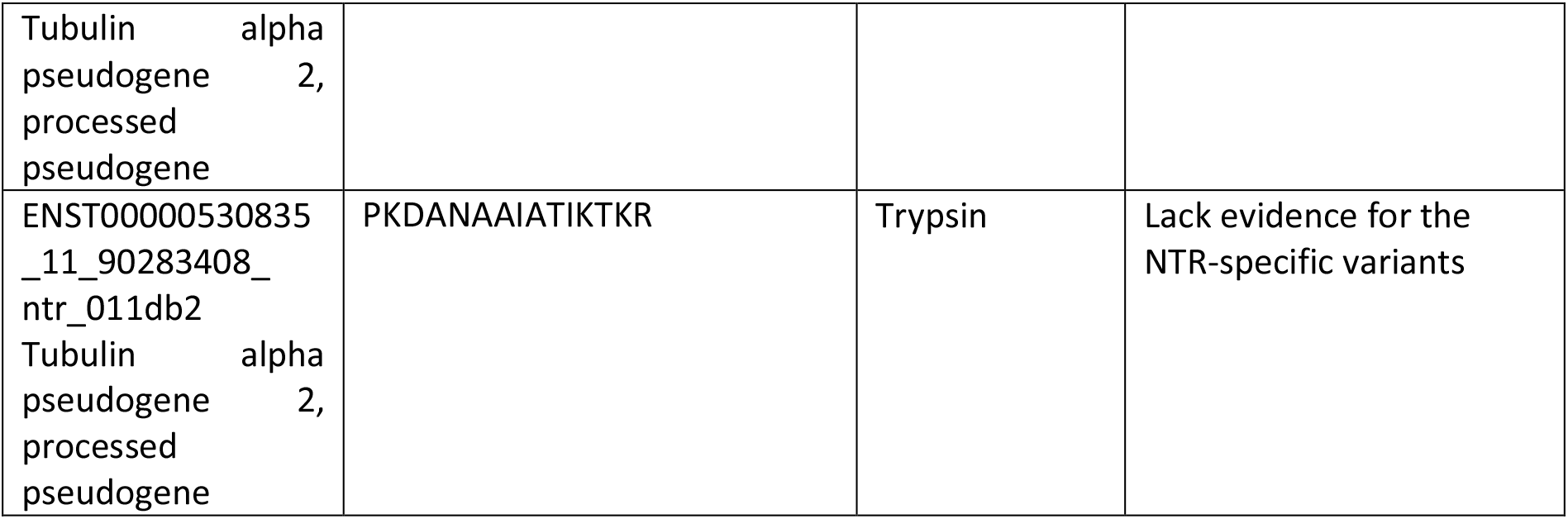
List of all peptides that matched to an NTR entry which were retained after inspection of MS/MS data. For each peptide, the accessions (containing both the ENSMBL accession and transcript information) are provided along with the identified peptide sequence and the enzyme (protease) the peptide was identified with.

**Figure 9:**
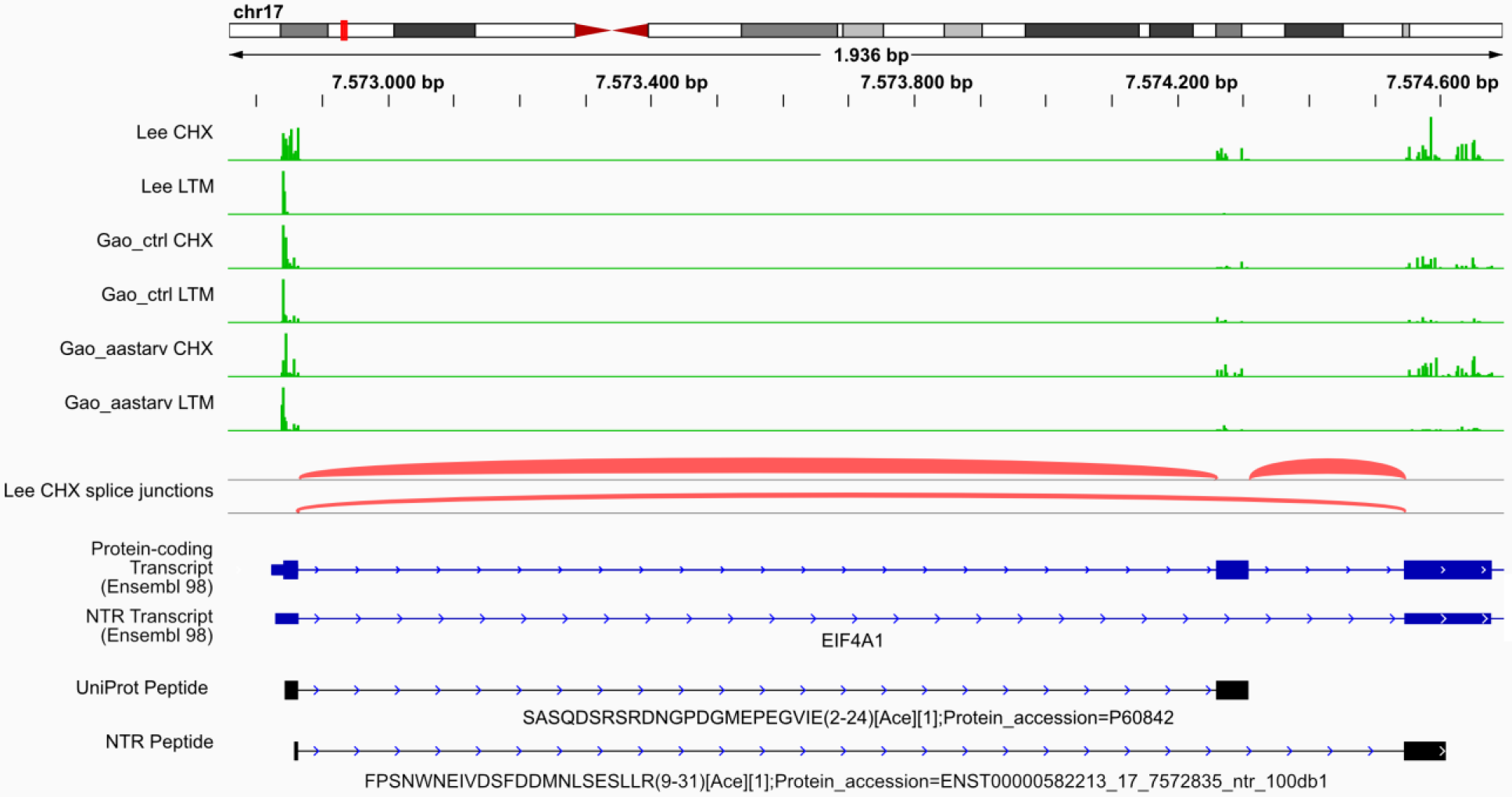
Omics evidence for an EIF4A1 proteoform visualized using a genome browser. FPSNWNEIVDSFDDMNLSESLLR peptide of the Eukaryotic translation initiation factor 4A1 (EIF4A1) derived from a retained intron transcript is supported by NTR-specific, alternatively spliced reads. The six top tracks represent ribosome profiling evidence of translation in Watson (green) or Crick orientation (red). We used two published studies ((4,34)) as source of data for elongating ribosomes treated with cycloheximide (CHX) and initiating ribosomes treated with lactimidomycin (LTM), harvested from HEK cells under normal conditions (“Lee” and “Gao_ctrl”) or under amino acid deprivation (“Gao_aastarv”). The red tracks display the experimentally verified splice junctions. Next, transcripts (from Ensembl annotation) and genome-mapped peptides (from our study) are shown. Increasing line thickness represents introns, exons and CDS, respectively and arrows mark the direction of translation. The peptide name consists of peptide sequence, start - end position, N-terminal modification, spectral count and matching protein accession.

### 9. Virotrap data of a selected NTR protein

To further investigate the functionality of NTR proteins we selected one NTR protein to study its interactome by Virotrap. A Nt-peptide pointing to an Nt-proteoform (missing the first 11 AA) of ACTB pseudogene 8 (ENST00000403258_6_88276364_ntr_101db1) was identified and we decided to select both forms (full length and proteoform) for interactome analysis. The proteoforms were coupled to HIV-1 GAG protein and expressed in HEK293T cells. Expression of the fusion protein initiates budding of viral-like particles (VLPs) from the cells, which contain the bait protein and its interactions partners. The VLPs can thereafter be purified from the growth medium as a genetic fusion protein of the vesicular stomatitis virus G protein (VSV-G) coupled to a Flag-tag is co-transfected and expressed on the surface of the VLPs (41). We performed triplicate Virotrap experiments for both NTR proteoforms to obtain specific interaction partners for the NTR protein. *E. coli* dihydrofolate reductase (eDHFR) fused to GAG was used as a negative control. Mass spectrometry of the VLPS revealed that both proteoforms were successfully expressed and pairwise comparisons between either the full length or the Nt-proteoform with the eDHFR control samples revealed several potential interaction partners of the NTR protein (11 for the full length and 10 for the proteoform, not taking the bait itself into account, **Figures 10A** and **B**). Four common interaction partners were identified: CTNNA1, SNX2, HGS and CTBP1/CTBP2. All significant proteins (adjusted p-value <0.01) in at least one comparison were also visualized in a heatmap (**Figure 10C**), revealing 54 significant proteins in the control samples and 18 significant proteins in the bait samples. The NTR proteins seem to interact with several proteins that are found to localize at membranes (IRS4, TMEM219, CTNNA1, MAT2B, PI4KA and CMTM6) and/or function in vesicles and protein transport (TFG, SNX2, HGS, RER1 and CTMT6) thus providing molecular functions to this novel protein.

**Figure 10:**
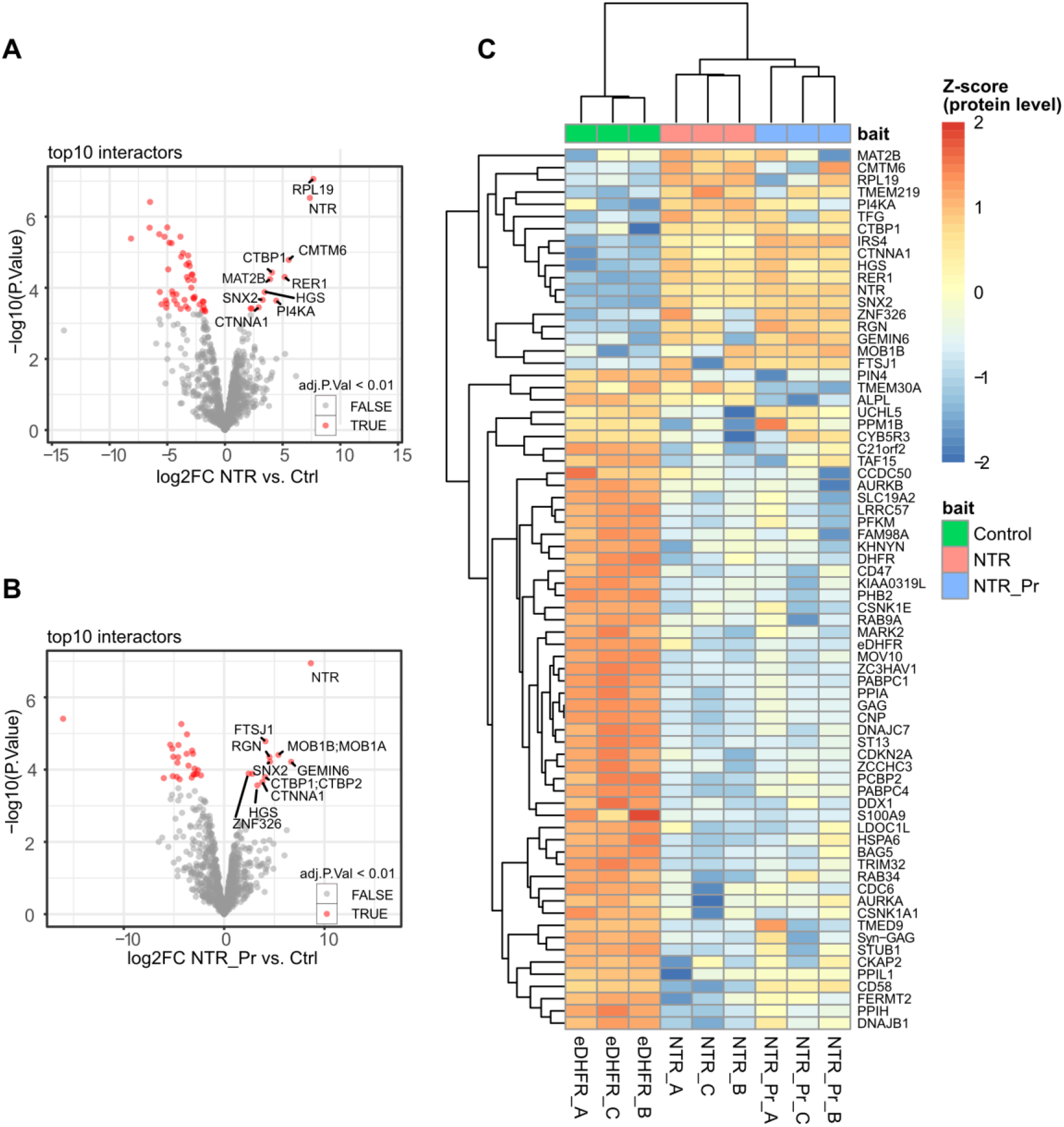
Virotrap interactome analysis of a novel protein and its N-terminal proteoform. Virotrap screens were performed in HEK293T cells using two proteoforms of the ACTB pseudogene 8 (annotated as NTR in the figures) as baits. *Escherichia coli* dihydrofolate reductase (eDHFR) fused to GAG was used as a negative control. A and B) Volcano plots showing the interactions partners of A) the full-length protein and B) its N-terminal, 11 amino acids shorter proteoform. Proteins with significantly altered levels are indicated in red and were defined through a pairwise t-test (FDR < 0.01). The x-axis shows the log_2_ fold change (FC) of the proteins in the ACTB pseudogene 8 Virotrap studies compared to the negative control, while the y-axis shows the –log_10_ of the adjusted p-values. C) Heatmap of all proteins found with significantly altered levels in at least one of the pairwise comparisons between eDHFR and the ACTB pseudogene 8 proteoforms. The scale shows Z-scored site intensity values.

## Discussion

The discrepancy between the number of NTR proteins predicted from RNA analyses (such as Ribo-seq and RNA sequencing) (3,5,7,8,12–14) and the number of unambiguously detected NTR protein products using mass spectrometry-based proteomics was the main motive for performing our study. We used an experimental setup that combined a reduced sample complexity with an extended database search space to improve the possibility of identifying NTR proteins. To cope with the increased search space, we introduced a rigorous workflow for data analysis and curation of the results.

In 2018, Na *et al*. (72) used a combination of N-terminal peptide enrichment, a Ribo-seq-based search database and downstream filtering in their search for protein evidence of translation starting at non-canonical translation initiation sites. Similarly to their study, we reduced the sample complexity by focusing on protein Nt-peptides only, as these peptides are proxies for translation events and can provide direct evidence for NTR proteins. However, restricting ourselves to Nt-peptides comes with a cost. Indeed, by not using shotgun proteomics data, we will have missed identification of several NTR proteins. In addition, protein identification in N-terminomics studies is typically based on a single peptide, with not all of these peptides being identifiable. To counter this effect we used three different proteases for proteome digestion as these generate different N-terminal peptides. Our bioinformatic analysis revealed that our approach should, in theory, greatly improve identification of peptides from NTR protein products however, less than 1% of the identified Nt-peptides was found to originate from such proteins.

With complex databases such as the one used in our study, the protein inference problem is highly prevalent. Therefore, our filtering approach started with the re-assignment of protein entries to identified peptides and, when peptides were matched to database entries with different levels of evidence for protein expression and protein (functional) annotation, we gave priority to the best annotated ones (UniProtKB entries). As expected, this led to the highest reduction in the apparent novel proteins identified (a decrease from 6% of all identifications to <0.6 %) as most (> 91%) of the NTR peptides were now re-matched to UniProt protein entries. Several other studies have used similar or slightly different strategies (for example, filtering out all peptides originating from known proteins or filtering out all non-unique proteins) to find novel proteins in their proteogenomics workflows (12,18,32,61,73,74). Filtering of our results continued with the deduplication and removal of non-N-terminal peptides, reducing the number of NTR proteins identified to just 39. The remaining peptides were expected to originate either from translation or from protein processing. The former start with an initiator methionine, which can be co-translationally removed by MetAPs. Co-translational Nα-acetylation of the initiator methionine or the exposed second amino acid, is a frequent modification occurring on eukaryotic intracellular proteins. Both features were used to remove peptides that most likely originated from protein processing, followed by a final merge of the results from the three different proteases used. Ultimately, this lead to translational evidence for only 19 NTR proteins (or 0.78% of all identified proteins).

To account for the fact that isobaric amino acids (or amino acid combinations) and amino acid modifications might have influenced correct identification of MS/MS spectra, a BlastP analysis of these 19 NTR peptides was performed. This revealed nine exact matches to (high abundant) annotated proteins. The remaining ten were found to be highly similar to annotated proteins as well, only differing by one or two amino acids. When evaluating the PSMs of these peptides, two more peptides (pointing to the same NTR protein) were removed. Additional comparisons of the fragment ions of the peptides identified in our sample with synthetic peptides resulted in the removal of an extra peptide. To verify the coverage of NTR-specific transcript variants present in the identified Nt-peptides using Ribo-seq data, we visualized their genomic locations with the corresponding Ribo-seq reads, which showed that many of the NTR-specific nucleotide variants were not unambiguously covered by Ribo-seq reads. In fact, only four of seven NTR proteins were highly supported.

It must be clear that all of the above indicates that stringent curation and inspection of database search results and proteomics data are essential to report identification of NTR proteins as several shortcomings inherent to MS data analysis wrongly assigned peptides to NTR proteins. In this respect, Kim *et al*. (16) reported several peptides pointing to translation of non-coding RNA’s, but upon checking some of these peptides by BlastP, we found an exact match to UniProt proteins. For instance, the peptide VLGSAPPPFTPSLLEQEVR was linked to a non-coding RNA (LOC113230), while BlastP revealed an exact match to MISP3 (starting at position 140, UniProt accession: Q96FF7)). In fact, of the nine proteins reported to originate from novel protein coding regions (more specifically non-coding RNA’s) we found for six of them that the identified peptide(s) had an exact match to a UniProt protein (**Supplementary Table S4**). This might point to misannotation of long non-coding RNA’s.

Several databases such as OpenProt (74), sORFs (75) and smPROT (76) hold sequences of novel proteins or proteins originating from NTRs. When available, these databases also report MS evidence for such predicted proteins. Considering the NTR proteins identified in our study, often (slightly) different protein sequences are reported from the same transcripts. When evaluating our ten most confident NTR proteins in these databases, surprisingly, one for one of them MS evidence was found in one database. For MASAASSSSLE we found a similar protein in the OpenProt database (accession: IP_591792, reported as an alternative protein), which is N-terminally 7 AA longer (MCDIKEK) compared to the NTR protein (ENST00000403258) we identified here. OpenProt reports several IP_591792 peptide matches found by MS, among these is EMASAASSSSLEK, which holds the Nt-peptide we have identified (MASAASSSSLE).

We also evaluated if Nt-peptides from Nt-proteoforms generated by alternative translation initiation or alternative splicing, or from translation of small upstream ORFs located in the 5’UTR of a regular CDS were identified. Indeed, we detected 31 N-termini originating from an annotated start site in ENSMBL (aTIS), nine N-termini pointing to proteoforms located inside an annotated coding sequence in ENSMBL (CDS), and 22 proteoforms located in the 5’UTR region. However, we were not able to detect protein products from 3’UTR regions, in line with previous data that translation from 5’UTRs is more frequent than from 3’UTRs [20]. Further, we identified 92 Nt-peptides that uniquely match to a UniProt isoform and 689 Nt-peptides that match to an internal position of a UniProt protein and thus point to possible Nt-proteoforms (**Supplementary Table S5**).

A possible explanation for the discrepancy between the reported number of translated NTRs by Ribo-seq and the number of NTR proteins for which peptide products were identified by mass spectrometry is a potential overestimation by Ribo-seq. Several papers have raised concerns about a need for standardization as biases in both sample preparation (due to the RNase used, use of antibiotics, method and buffer for cell lysis, …) and data processing (library assembly, normalization, quality cut-off thresholds, handling of non-unique reads and mismatches, …) can greatly influence the translational evidence reported (77–82).

Besides Ribo-seq and proteogenomics, other methods exist to study or monitor protein synthesis. Some proteomics-based methods rely on the incorporation of an azidohomoalanine (a bio-orthogonal methionine analog), which can then be used for affinity based purification (83,84). However, such methods were only able to identify a few hundreds of proteins. A more recent method called PUNCH-P recovers ribosome-nascent chain complexes from the cells by ultracentrifugation followed by labeling with biotin-puromycin and affinity purification before LC-MS/MS analysis. With this method, thousands of proteins could be detected (85). However, PUNCH-P is unable to evaluate degradation, protein stability and post-translational modifications. Hence, proteins found by this method might not be stable or functional.

Another possible reason for the difference between Ribo-seq and MS data is that the expression of (non-coding) genes is tissue-specific (86,87) (and likely also depends on the cell cycle phase and stimuli). We thus likely miss several NTR proteins by only analyzing HEK293T proteomes under normal, ideal growth conditions. However, from the Ribo-seq reported NTR proteins that were assumed to be translated in HEK293T cells (16,919 reported in our database) we only find protein evidence for 0.11% of them. Clearly, by restricting to cytosolic proteins, we will also have missed proteins present at other subcellular localizations. As shown recently, it might be interesting to focus on the immunopeptidome as Cuevas *et al*. (2021) showed that proteins originating from non-coding genes are more likely to be detected in the immunopeptideome compared to canonical proteins, hinting to the fact that they might be non-functional (24). Similar as in our study they only found translational evidence (by MS) for 0.44% of the non-canonical proteins reported by their RNA-seq and Ribo-seq experiments.

We performed an interactome analysis using Virotrap to evaluate apparently stable NTR proteins using one NTR protein that survived our filtering steps, ACTB pseudogene 8 (ENST00000403258_6_88276364_ntr_101db1). When used as a Virotrap bait, we found that this protein had 18 potential interaction partners that mainly function in vesicle/protein transport and/or were found to localize at membranes, thus assigning functionalities to this particular NTR protein.

In summary, our data let us suspect that the majority of currently annotated non-coding regions are indeed non-coding however, the few that are translated and give rise to stable proteins might be functional.

## Data Availability

The mass spectrometry proteomics data have been deposited to the ProteomeXchange Consortium via the PRIDE (88) partner repository with the following dataset identifiers:

- PXD030601(cytosolic N-terminal COFRADIC data from the three different proteases), data is private before publishing and can be accessed with the following login credentials: username: Password: 2V9lL6fi
- PXD030216 (Virotrap data), data is private before publishing and can be accessed with the following login credentials: username: Password: KkY1Ycjn.

The mass spectrometry data of the synthetic peptides and the comparison with the peptides identified in our samples has been deposited to the ProteomeXchange Consortium via the PanormaPublic repository with the dataset identifier PXD030285 (data is private before publishing and can be accessed with the following link: https://panoramaweb.org/pakkBx.url and using the following login credentials: username: and password: mmZcpgPd).

Other relevant data are included in the paper and accompanying Supplementary Data or are available from the corresponding author upon reasonable request.

## Supplementary data

Supplementary Data are available at NAR online.

## Acknowledgment

We thank Dr. Patrick Willems, Jarne Pauwels and Jade Hawksworth of the Gevaert lab for their suggestions and helpful discussions. A.B. performed all experiments and wrote the manuscript. D.F created the custom build database, performed the peptide detectability analysis, set up the data filtering pipeline and inspected the sequencing data together with A.B. A.S. assisted the COFRADIC experiments and analyzed the synthetic peptides. T.V.d.S. generated the NTR clones and performed the Virotrap experiment. H.D. generated synthetic peptides. K.G. with A.B. managed the project and wrote and edited the manuscript.

## REFERENCES

1. Smith, L.M., Kelleher, N.L. and Consortium for Top Down, P. (2013) Proteoform: a single term describing protein complexity. Nat Methods, 10, 186–187.

2. Bogaert, A., Fernandez, E. and Gevaert, K. (2020) N-Terminal Proteoforms in Human Disease. Trends Biochem Sci, 45, 308–320.

3. Ingolia, N.T., Lareau, L.F. and Weissman, J.S. (2011) Ribosome profiling of mouse embryonic stem cells reveals the complexity and dynamics of mammalian proteomes. Cell, 147, 789–802.

4. Lee, S., Liu, B., Huang, S.X., Shen, B. and Qian, S.B. (2012) Global mapping of translation initiation sites in mammalian cells at single-nucleotide resolution. Proc Natl Acad Sci U S A, 109, E2424–2432.

5. Ingolia, N.T., Brar, G.A., Stern-Ginossar, N., Harris, M.S., Talhouarne, G.J., Jackson, S.E., Wills, M.R. and Weissman, J.S. (2014) Ribosome profiling reveals pervasive translation outside of annotated protein-coding genes. Cell Rep, 8, 1365–1379.

6. Mouilleron, H., Delcourt, V. and Roucou, X. (2015) Death of a dogma: eukaryotic mRNAs can code for more than one protein. Nucleic Acids Res.

7. Slavoff, S.A., Mitchell, A.J., Schwaid, A.G., Cabili, M.N., Ma, J., Levin, J.Z., Karger, A.D., Budnik, B.A., Rinn, J.L. and Saghatelian, A. (2013) Peptidomic discovery of short open reading frame-encoded peptides in human cells. Nat Chem Biol, 9, 59–64.

8. Samandi, S., Roy, A.V., Delcourt, V., Lucier, J.F., Gagnon, J., Beaudoin, M.C., Vanderperre, B., Breton, M.A., Motard, J., Jacques, J.F. et al. (2017) Deep transcriptome annotation enables the discovery and functional characterization of cryptic small proteins. Elife, 6.

9. Delcourt, V., Staskevicius, A., Salzet, M., Fournier, I. and Roucou, X. (2018) Small Proteins Encoded by Unannotated ORFs are Rising Stars of the Proteome, Confirming Shortcomings in Genome Annotations and Current Vision of an mRNA. Proteomics, 18, e1700058.

10. Brunet, M.A., Leblanc, S. and Roucou, X. (2020) Reconsidering proteomic diversity with functional investigation of small ORFs and alternative ORFs. Exp Cell Res, 393, 112057.

11. Gibb, E.A., Brown, C.J. and Lam, W.L. (2011) The functional role of long non-coding RNA in human carcinomas. Mol Cancer, 10, 38.

12. Brunet, M.A., Brunelle, M., Lucier, J.F., Delcourt, V., Levesque, M., Grenier, F., Samandi, S., Leblanc, S., Aguilar, J.D., Dufour, P. et al. (2019) OpenProt: a more comprehensive guide to explore eukaryotic coding potential and proteomes. Nucleic Acids Res, 47, D403–D410.

13. Bazzini, A.A., Johnstone, T.G., Christiano, R., Mackowiak, S.D., Obermayer, B., Fleming, E.S., Vejnar, C.E., Lee, M.T., Rajewsky, N., Walther, T.C. et al. (2014) Identification of small ORFs in vertebrates using ribosome footprinting and evolutionary conservation. EMBO J, 33, 981–993.

14. Frith, M.C., Forrest, A.R., Nourbakhsh, E., Pang, K.C., Kai, C., Kawai, J., Carninci, P., Hayashizaki, Y., Bailey, T.L. and Grimmond, S.M. (2006) The abundance of short proteins in the mammalian proteome. PLoS Genet, 2, e52.

15. Verheggen, K., Volders, P.J., Mestdagh, P., Menschaert, G., Van Damme, P., Gevaert, K., Martens, L. and Vandesompele, J. (2017) Noncoding after All: Biases in Proteomics Data Do Not Explain Observed Absence of lncRNA Translation Products. J Proteome Res, 16, 2508–2515.

16. Kim, M.S., Pinto, S.M., Getnet, D., Nirujogi, R.S., Manda, S.S., Chaerkady, R., Madugundu, A.K., Kelkar, D.S., Isserlin, R., Jain, S. et al. (2014) A draft map of the human proteome. Nature, 509, 575–581.

17. Crappe, J., Ndah, E., Koch, A., Steyaert, S., Gawron, D., De Keulenaer, S., De Meester, E., De Meyer, T., Van Criekinge, W., Van Damme, P. et al. (2015) PROTEOFORMER: deep proteome coverage through ribosome profiling and MS integration. Nucleic Acids Res, 43, e29.

18. Koch, A., Gawron, D., Steyaert, S., Ndah, E., Crappe, J., De Keulenaer, S., De Meester, E., Ma, M., Shen, B., Gevaert, K. et al. (2014) A proteogenomics approach integrating proteomics and ribosome profiling increases the efficiency of protein identification and enables the discovery of alternative translation start sites. Proteomics, 14, 2688–2698.

19. Ma, J., Ward, C.C., Jungreis, I., Slavoff, S.A., Schwaid, A.G., Neveu, J., Budnik, B.A., Kellis, M. and Saghatelian, A. (2014) Discovery of human sORF-encoded polypeptides (SEPs) in cell lines and tissue. J Proteome Res, 13, 1757–1765.

20. Schwaid, A.G., Shannon, D.A., Ma, J., Slavoff, S.A., Levin, J.Z., Weerapana, E. and Saghatelian, A. (2013) Chemoproteomic discovery of cysteine-containing human short open reading frames. J Am Chem Soc, 135, 16750–16753.

21. Pauli, A., Valen, E. and Schier, A.F. (2015) Identifying (non-)coding RNAs and small peptides: challenges and opportunities. Bioessays, 37, 103–112.

22. Uszczynska-Ratajczak, B., Lagarde, J., Frankish, A., Guigo, R. and Johnson, R. (2018) Towards a complete map of the human long non-coding RNA transcriptome. Nat Rev Genet, 19, 535–548.

23. Gawron, D., Gevaert, K. and Van Damme, P. (2014) The proteome under translational control. Proteomics, 14, 2647–2662.

24. Ruiz Cuevas, M.V., Hardy, M.P., Holly, J., Bonneil, E., Durette, C., Courcelles, M., Lanoix, J., Cote, C., Staudt, L.M., Lemieux, S., et al. (2021) Most non-canonical proteins uniquely populate the proteome or immunopeptidome. Cell Rep, 34, 108815.

25. Johnstone, T.G., Bazzini, A.A. and Giraldez, A.J. (2016) Upstream ORFs are prevalent translational repressors in vertebrates. EMBO J, 35, 706–723.

26. Chew, G.L., Pauli, A. and Schier, A.F. (2016) Conservation of uORF repressiveness and sequence features in mouse, human and zebrafish. Nat Commun, 7, 11663.

27. Anderson, D.M., Anderson, K.M., Chang, C.L., Makarewich, C.A., Nelson, B.R., McAnally, J.R., Kasaragod, P., Shelton, J.M., Liou, J., Bassel-Duby, R. et al. (2015) A micropeptide encoded by a putative long noncoding RNA regulates muscle performance. Cell, 160, 595–606.

28. Slavoff, S.A., Heo, J., Budnik, B.A., Hanakahi, L.A. and Saghatelian, A. (2014) A human short open reading frame (sORF)-encoded polypeptide that stimulates DNA end joining. J Biol Chem, 289, 10950–10957.

29. Rathore, A., Chu, Q., Tan, D., Martinez, T.F., Donaldson, C.J., Diedrich, J.K., Yates, J.R., 3rd and Saghatelian, A. (2018) MIEF1 Microprotein Regulates Mitochondrial Translation. Biochemistry, 57, 5564–5575.

30. Jackson, R., Kroehling, L., Khitun, A., Bailis, W., Jarret, A., York, A.G., Khan, O.M., Brewer, J.R., Skadow, M.H., Duizer, C. et al. (2018) The translation of non-canonical open reading frames controls mucosal immunity. Nature, 564, 434–438.

31. Nelson, B.R., Makarewich, C.A., Anderson, D.M., Winders, B.R., Troupes, C.D., Wu, F., Reese, A.L., McAnally, J.R., Chen, X., Kavalali, E.T. et al. (2016) A peptide encoded by a transcript annotated as long noncoding RNA enhances SERCA activity in muscle. Science, 351, 271–275.

32. Chen, J., Brunner, A.D., Cogan, J.Z., Nunez, J.K., Fields, A.P., Adamson, B., Itzhak, D.N., Li, J.Y., Mann, M., Leonetti, M.D. et al. (2020) Pervasive functional translation of noncanonical human open reading frames. Science, 367, 1140–1146.

33. UniProt, C. (2021) UniProt: the universal protein knowledgebase in 2021. Nucleic Acids Res, 49, D480–D489.

34. Gao, X., Wan, J., Liu, B., Ma, M., Shen, B. and Qian, S.B. (2015) Quantitative profiling of initiating ribosomes in vivo. Nat Methods, 12, 147–153.

35. Verbruggen, S., Ndah, E., Van Criekinge, W., Gessulat, S., Kuster, B., Wilhelm, M., Van Damme, P. and Menschaert, G. (2019) PROTEOFORMER 2.0: Further Developments in the Ribosome Profiling-assisted Proteogenomic Hunt for New Proteoforms. Mol Cell Proteomics, 18, S126–S140.

36. Staes, A., Van Damme, P., Helsens, K., Demol, H., Vandekerckhove, J. and Gevaert, K. (2008) Improved recovery of proteome-informative, protein N-terminal peptides by combined fractional diagonal chromatography (COFRADIC). Proteomics, 8, 1362–1370.

37. Willems, P., Ndah, E., Jonckheere, V., Stael, S., Sticker, A., Martens, L., Van Breusegem, F., Gevaert, K. and Van Damme, P. (2017) N-terminal Proteomics Assisted Profiling of the Unexplored Translation Initiation Landscape in Arabidopsis thaliana. Mol Cell Proteomics, 16, 1064–1080.

38. McDonald, L., Robertson, D.H., Hurst, J.L. and Beynon, R.J. (2005) Positional proteomics: selective recovery and analysis of N-terminal proteolytic peptides. Nat Methods, 2, 955–957.

39. Yeom, J., Ju, S., Choi, Y., Paek, E. and Lee, C. (2017) Comprehensive analysis of human protein N-termini enables assessment of various protein forms. Sci Rep, 7, 6599.

40. Kaulich, P.T., Cassidy, L., Bartel, J., Schmitz, R.A. and Tholey, A. (2021) Multi-protease Approach for the Improved Identification and Molecular Characterization of Small Proteins and Short Open Reading Frame-Encoded Peptides. J Proteome Res, 20, 2895–2903.

41. Eyckerman, S., Titeca, K., Van Quickelberghe, E., Cloots, E., Verhee, A., Samyn, N., De Ceuninck, L., Timmerman, E., De Sutter, D., Lievens, S. et al. (2016) Trapping mammalian protein complexes in viral particles. Nat Commun, 7, 11416.

42. Alberts, B. (2008) Molecular biology of the cell. 5th ed. Garland Science, New York.

43. Frottin, F., Martinez, A., Peynot, P., Mitra, S., Holz, R.C., Giglione, C. and Meinnel, T. (2006) The proteomics of N-terminal methionine cleavage. Mol Cell Proteomics, 5, 2336–2349.

44. Bradshaw, R.A., Brickey, W.W. and Walker, K.W. (1998) N-terminal processing: the methionine aminopeptidase and N alpha-acetyl transferase families. Trends Biochem Sci, 23, 263–267.

45. Arnesen, T., Van Damme, P., Polevoda, B., Helsens, K., Evjenth, R., Colaert, N., Varhaug, J.E., Vandekerckhove, J., Lillehaug, J.R., Sherman, F. et al. (2009) Proteomics analyses reveal the evolutionary conservation and divergence of N-terminal acetyltransferases from yeast and humans. Proc Natl Acad Sci U S A, 106, 8157–8162.

46. Varland, S., Osberg, C. and Arnesen, T. (2015) N-terminal modifications of cellular proteins: The enzymes involved, their substrate specificities and biological effects. Proteomics, 15, 2385–2401.

47. Aksnes, H., Drazic, A., Marie, M. and Arnesen, T. (2016) First Things First: Vital Protein Marks by N-Terminal Acetyltransferases. Trends Biochem Sci, 41, 746–760.

48. Demir, F., Niedermaier, S., Kizhakkedathu, J.N. and Huesgen, P.F. (2017) Profiling of Protein N-Termini and Their Modifications in Complex Samples. Methods Mol Biol, 1574, 35–50.

49. van Loo, G., Schotte, P., van Gurp, M., Demol, H., Hoorelbeke, B., Gevaert, K., Rodriguez, I., Ruiz-Carrillo, A., Vandekerckhove, J., Declercq, W. et al. (2001) Endonuclease G: a mitochondrial protein released in apoptosis and involved in caspase-independent DNA degradation. Cell Death Differ, 8, 1136–1142.

50. Staes, A., Van Damme, P., Timmerman, E., Ruttens, B., Stes, E., Gevaert, K. and Impens, F. (2017) Protease Substrate Profiling by N-Terminal COFRADIC. Methods Mol Biol, 1574, 51–76.

51. Helsens, K., Colaert, N., Barsnes, H., Muth, T., Flikka, K., Staes, A., Timmerman, E., Wortelkamp, S., Sickmann, A., Vandekerckhove, J. et al. (2010) ms_lims, a simple yet powerful open source laboratory information management system for MS-driven proteomics. Proteomics, 10, 1261–1264.

52. MacLean, B., Tomazela, D.M., Shulman, N., Chambers, M., Finney, G.L., Frewen, B., Kern, R., Tabb, D.L., Liebler, D.C. and MacCoss, M.J. (2010) Skyline: an open source document editor for creating and analyzing targeted proteomics experiments. Bioinformatics, 26, 966–968.

53. Ning, Z., Seebun, D., Hawley, B., Chiang, C.K. and Figeys, D. (2013) From cells to peptides: “one-stop” integrated proteomic processing using amphipols. J Proteome Res, 12, 1512–1519.

54. Ritchie, M.E., Phipson, B., Wu, D., Hu, Y., Law, C.W., Shi, W. and Smyth, G.K. (2015) limma powers differential expression analyses for RNA-sequencing and microarray studies. Nucleic Acids Res, 43, e47.

55. Ingolia, N.T., Ghaemmaghami, S., Newman, J.R. and Weissman, J.S. (2009) Genome-wide analysis in vivo of translation with nucleotide resolution using ribosome profiling. Science, 324, 218–223.

56. Koch, A., Gawron, D., Steyaert, S., Ndah, E., Crappe, J., De Keulenaer, S., De Meester, E., Ma, M., Shen, B., Gevaert, K. et al. (2014) A proteogenomics approach integrating proteomics and ribosome profiling increases the efficiency of protein identification and enables the discovery of alternative translation start sites. Proteomics, 14, 2688–2698.

57. Crappe, J., Ndah, E., Koch, A., Steyaert, S., Gawron, D., De Keulenaer, S., De Meester, E., De Meyer, T., Van Criekinge, W., Van Damme, P. et al. (2015) PROTEOFORMER: deep proteome coverage through ribosome profiling and MS integration. Nucleic Acids Research, 43.

58. Fijalkowska, D., Verbruggen, S., Ndah, E., Jonckheere, V., Menschaert, G. and Van Damme, P. (2017) eIF1 modulates the recognition of suboptimal translation initiation sites and steers gene expression via uORFs. Nucleic Acids Res.

59. Gevaert, K., Goethals, M., Martens, L., Van Damme, J., Staes, A., Thomas, G.R. and Vandekerckhove, J. (2003) Exploring proteomes and analyzing protein processing by mass spectrometric identification of sorted N-terminal peptides. Nat Biotechnol, 21, 566–569.

60. Thul, P.J., Akesson, L., Wiking, M., Mahdessian, D., Geladaki, A., Ait Blal, H., Alm, T., Asplund, A., Bjork, L., Breckels, L.M. et al. (2017) A subcellular map of the human proteome. Science, 356.

61. Van Damme, P., Gawron, D., Van Criekinge, W. and Menschaert, G. (2014) N-terminal proteomics and ribosome profiling provide a comprehensive view of the alternative translation initiation landscape in mice and men. Mol Cell Proteomics, 13, 1245–1261.

62. Hulsen, T., de Vlieg, J. and Alkema, W. (2008) BioVenn - a web application for the comparison and visualization of biological lists using area-proportional Venn diagrams. BMC Genomics, 9, 488.

63. Nesvizhskii, A.I. and Aebersold, R. (2005) Interpretation of shotgun proteomic data: the protein inference problem. Mol Cell Proteomics, 4, 1419–1440.

64. Kearse, M.G. and Wilusz, J.E. (2017) Non-AUG translation: a new start for protein synthesis in eukaryotes. Genes Dev, 31, 1717–1731.

65. Boyer, J.B., Dedieu, A., Armengaud, J., Verdie, P., Subra, G., Martinez, J. and Enjalbal, C. (2014) N- and O-acetylation of threonine residues in the context of proteomics. J Proteomics, 108, 369–372.

66. Zhu, Y., Orre, L.M., Johansson, H.J., Huss, M., Boekel, J., Vesterlund, M., Fernandez-Woodbridge, A., Branca, R.M.M. and Lehtio, J. (2018) Publisher Correction: Discovery of coding regions in the human genome by integrated proteogenomics analysis workflow. Nat Commun, 9, 1852.

67. Zhu, Y., Orre, L.M., Johansson, H.J., Huss, M., Boekel, J., Vesterlund, M., Fernandez-Woodbridge, A., Branca, R.M.M. and Lehtio, J. (2018) Discovery of coding regions in the human genome by integrated proteogenomics analysis workflow. Nat Commun, 9, 903.

68. Wright, J.C., Mudge, J., Weisser, H., Barzine, M.P., Gonzalez, J.M., Brazma, A., Choudhary, J.S. and Harrow, J. (2016) Improving GENCODE reference gene annotation using a high-stringency proteogenomics workflow. Nat Commun, 7, 11778.

69. Meinwald, Y.C., Stimson, E.R. and Scheraga, H.A. (1986) Deamidation of the asparaginyl-glycyl sequence. Int J Pept Protein Res, 28, 79–84.

70. Robinson, J.T., Thorvaldsdottir, H., Winckler, W., Guttman, M., Lander, E.S., Getz, G. and Mesirov, J.P. (2011) Integrative genomics viewer. Nat Biotechnol, 29, 24–26.

71. Sherry, S.T., Ward, M.H., Kholodov, M., Baker, J., Phan, L., Smigielski, E.M. and Sirotkin, K. (2001) dbSNP: the NCBI database of genetic variation. Nucleic Acids Res, 29, 308–311.

72. Na, C.H., Barbhuiya, M.A., Kim, M.S., Verbruggen, S., Eacker, S.M., Pletnikova, O., Troncoso, J.C., Halushka, M.K., Menschaert, G., Overall, C.M. et al. (2018) Discovery of noncanonical translation initiation sites through mass spectrometric analysis of protein N termini. Genome research, 28, 25–36.

73. Menschaert, G., Van Criekinge, W., Notelaers, T., Koch, A., Crappe, J., Gevaert, K. and Van Damme, P. (2013) Deep proteome coverage based on ribosome profiling aids mass spectrometry-based protein and peptide discovery and provides evidence of alternative translation products and near-cognate translation initiation events. Mol Cell Proteomics, 12, 1780–1790.

74. Brunet, M.A., Lucier, J.F., Levesque, M., Leblanc, S., Jacques, J.F., Al-Saedi, H.R.H., Guilloy, N., Grenier, F., Avino, M., Fournier, I. et al. (2021) OpenProt 2021: deeper functional annotation of the coding potential of eukaryotic genomes. Nucleic Acids Res, 49, D380–D388.

75. Olexiouk, V., Van Criekinge, W. and Menschaert, G. (2018) An update on sORFs.org: a repository of small ORFs identified by ribosome profiling. Nucleic Acids Res, 46, D497–D502.

76. Hao, Y., Zhang, L., Niu, Y., Cai, T., Luo, J., He, S., Zhang, B., Zhang, D., Qin, Y., Yang, F. et al. (2018) SmProt: a database of small proteins encoded by annotated coding and non-coding RNA loci. Brief Bioinform, 19, 636–643.

77. Gerashchenko, M.V. and Gladyshev, V.N. (2014) Translation inhibitors cause abnormalities in ribosome profiling experiments. Nucleic Acids Res, 42, e134.

78. Gerashchenko, M.V. and Gladyshev, V.N. (2017) Ribonuclease selection for ribosome profiling. Nucleic Acids Res, 45, e6.

79. Bartholomaus, A., Del Campo, C. and Ignatova, Z. (2016) Mapping the non-standardized biases of ribosome profiling. Biol Chem, 397, 23–35.

80. Santos, D.A., Shi, L., Tu, B.P. and Weissman, J.S. (2019) Cycloheximide can distort measurements of mRNA levels and translation efficiency. Nucleic Acids Res, 47, 4974–4985.

81. Sharma, P., Wu, J., Nilges, B.S. and Leidel, S.A. (2021) Humans and other commonly used model organisms are resistant to cycloheximide-mediated biases in ribosome profiling experiments. Nat Commun, 12, 5094.

82. Glaub, A., Huptas, C., Neuhaus, K. and Ardern, Z. (2020) Recommendations for bacterial ribosome profiling experiments based on bioinformatic evaluation of published data. J Biol Chem, 295, 8999–9011.

83. Dieterich, D.C., Link, A.J., Graumann, J., Tirrell, D.A. and Schuman, E.M. (2006) Selective identification of newly synthesized proteins in mammalian cells using bioorthogonal noncanonical amino acid tagging (BONCAT). Proc Natl Acad Sci U S A, 103, 9482–9487.

84. Howden, A.J., Geoghegan, V., Katsch, K., Efstathiou, G., Bhushan, B., Boutureira, O., Thomas, B., Trudgian, D.C., Kessler, B.M., Dieterich, D.C. et al. (2013) QuaNCAT: quantitating proteome dynamics in primary cells. Nat Methods, 10, 343–346.

85. Aviner, R., Geiger, T. and Elroy-Stein, O. (2014) Genome-wide identification and quantification of protein synthesis in cultured cells and whole tissues by puromycin-associated nascent chain proteomics (PUNCH-P). Nat Protoc, 9, 751–760.

86. Derrien, T., Johnson, R., Bussotti, G., Tanzer, A., Djebali, S., Tilgner, H., Guernec, G., Martin, D., Merkel, A., Knowles, D.G. et al. (2012) The GENCODE v7 catalog of human long noncoding RNAs: analysis of their gene structure, evolution, and expression. Genome Res, 22, 1775–1789.

87. Cabili, M.N., Trapnell, C., Goff, L., Koziol, M., Tazon-Vega, B., Regev, A. and Rinn, J.L. (2011) Integrative annotation of human large intergenic noncoding RNAs reveals global properties and specific subclasses. Genes Dev, 25, 1915–1927.

88. Perez-Riverol, Y., Csordas, A., Bai, J., Bernal-Llinares, M., Hewapathirana, S., Kundu, D.J., Inuganti, A., Griss, J., Mayer, G., Eisenacher, M. et al. (2019) The PRIDE database and related tools and resources in 2019: improving support for quantification data. Nucleic Acids Res, 47, D442–D450.

