## Supplementary Figures S1-S5 for "Limited, but potentially functional translation of non-coding transcripts in the HEK293T cellular cytosol"

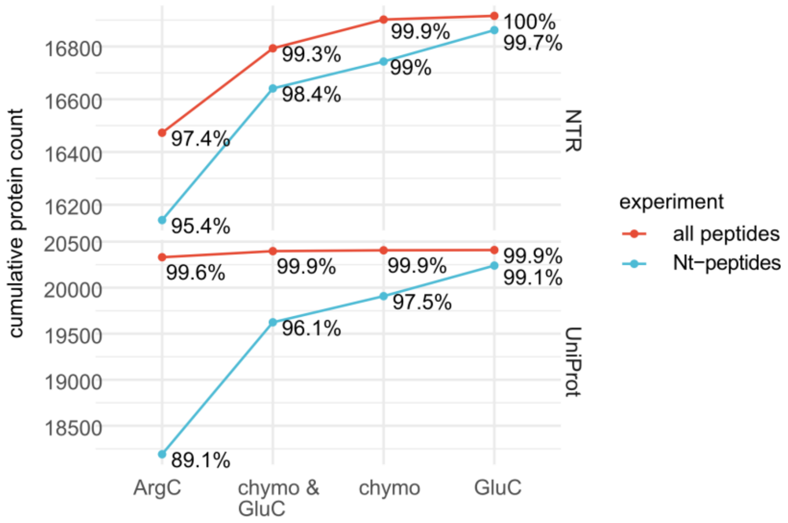


**Figure S1: Advantages of using three proteases for covering NTR and UniProt proteins.** Digestion with ArgC is estimated to give a quite complete proteome coverage (89.1% and 95.4% of UniProt and NTR proteoforms) but almost full coverage can be achieved by parallel processing samples with chymotrypsin and GluC.  Note that “chymo” or “GluC” point to protein identifications exclusively gained by using chymotrypsin or GluC, and to “chymo & GluC” indicates protein identifications gained in both conditions.

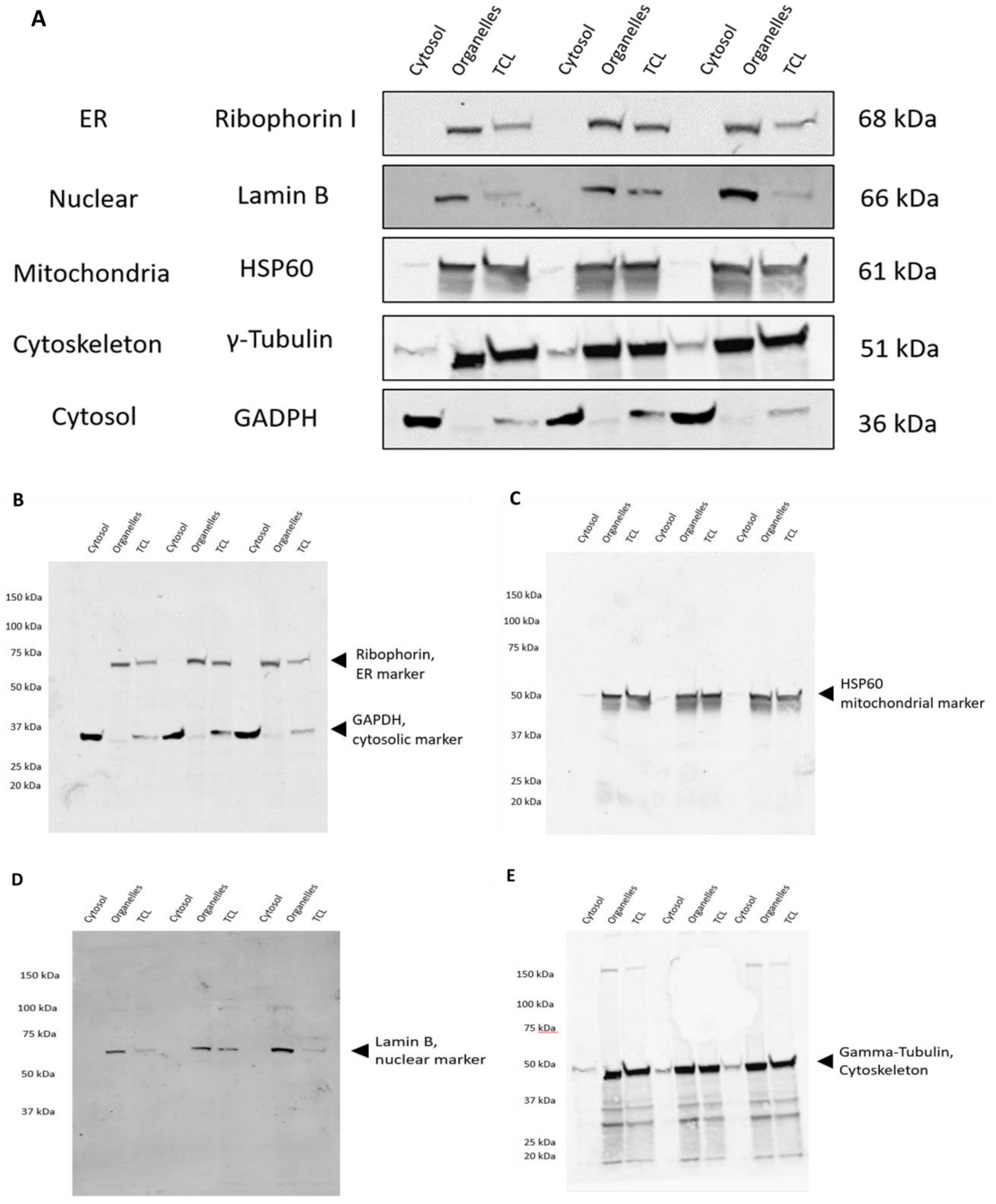


**Figure S2: Efficiency of the use of digitonin for extracting cytosolic proteins from HEK293T cells.** $10^7$ cells were collected and lysed in 1.25 ml of CFS buffer containing 0.02% digitonin. Following centrifugation, the supernatant was collected and the cell pellet was further extracted in RIPA lysis buffer. A total cell lysate was made by lysis of $10^7$ HEK293T cells in RIPA lysis buffer. An aliquot of each extract was subjected to 4-12% SDS-PAGE followed by Western blot (on PVDF membrane) using an anti-GAPDH antibody as a marker of the cytosol, an anti-HSP60 antibody as a marker for the mitochondria, an anti-LaminB antibody as a marker for the nucleus, anti-γ-Tubulin as a marker for the cytoskeleton and an antibody against Ribophorin I as marker for the endoplasmic reticulum. A) ER and nuclear markers were absent in the cytosolic fraction, whilst clearly visible in the organelle fraction and the total cell lysate. For HSP60 (mitochondrial marker) and γ-Tubulin (cytoskeleton marker), faint staining was observed in the cytosolic fraction and intense staining in the organelle fraction and the TC total cell lysate. The cytosol marker (GAPDH) is not found in the organelle fraction, while intense in the cytosolic fraction, and this intensity is clearly higher compared to that in the total cell lysate B-E) uncropped Western Blot results.

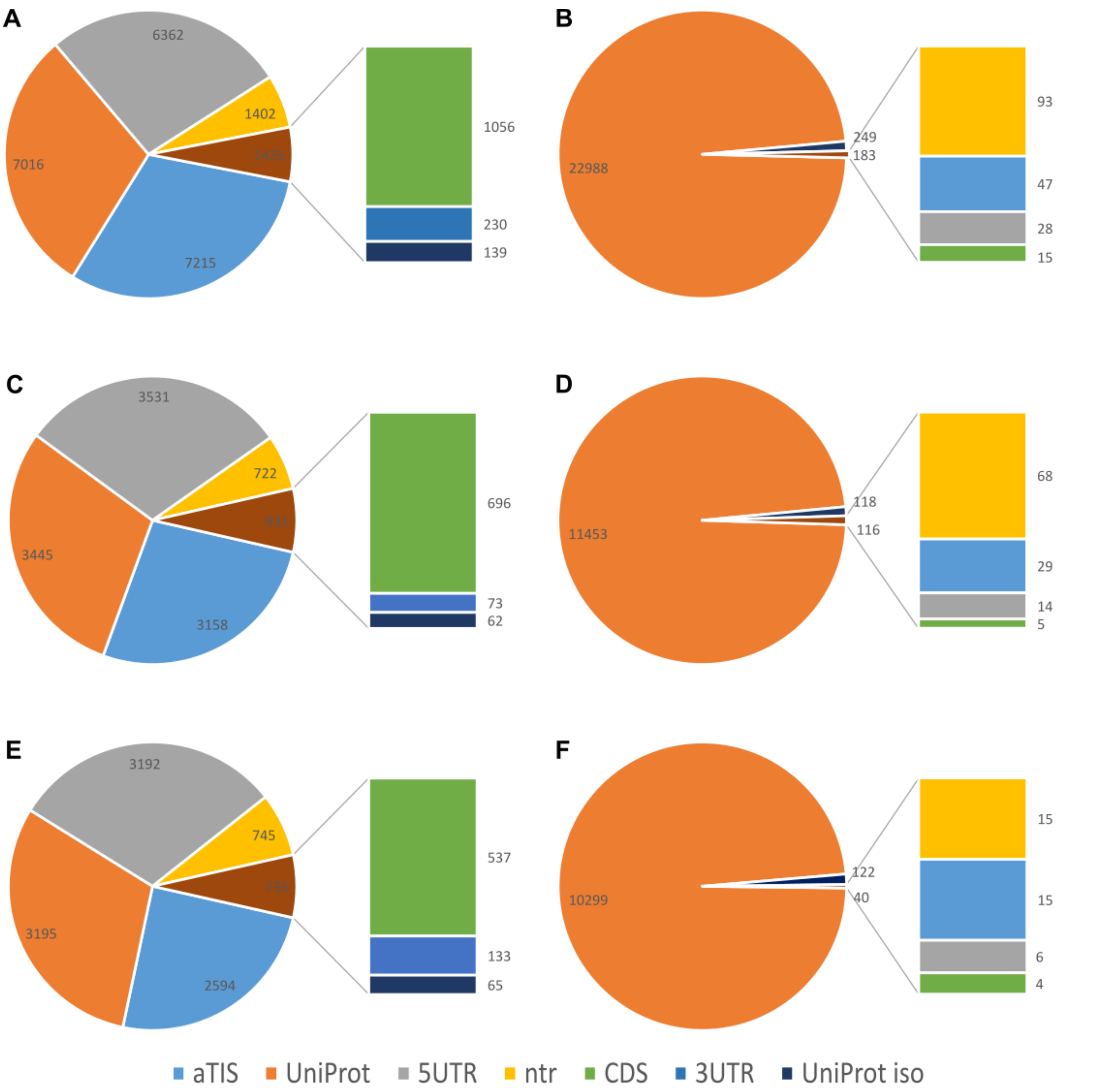


**Figure S3: Pie chart showing the distribution of all types of accession** for the trypsin digested sample before (A) and after accession sorting (B), for the chymotrypsin digested sample before (C) and after accession sorting (D) and for the GluC digested sample before (E) and after (F) accession sorting.

**A**

ADDAGAAGGPGGPGGPEMGNRGGFRGGF

Ace-ADDAGAAGGPGGPGGPEM<Mox*>GNRGGFRGGF-COOH

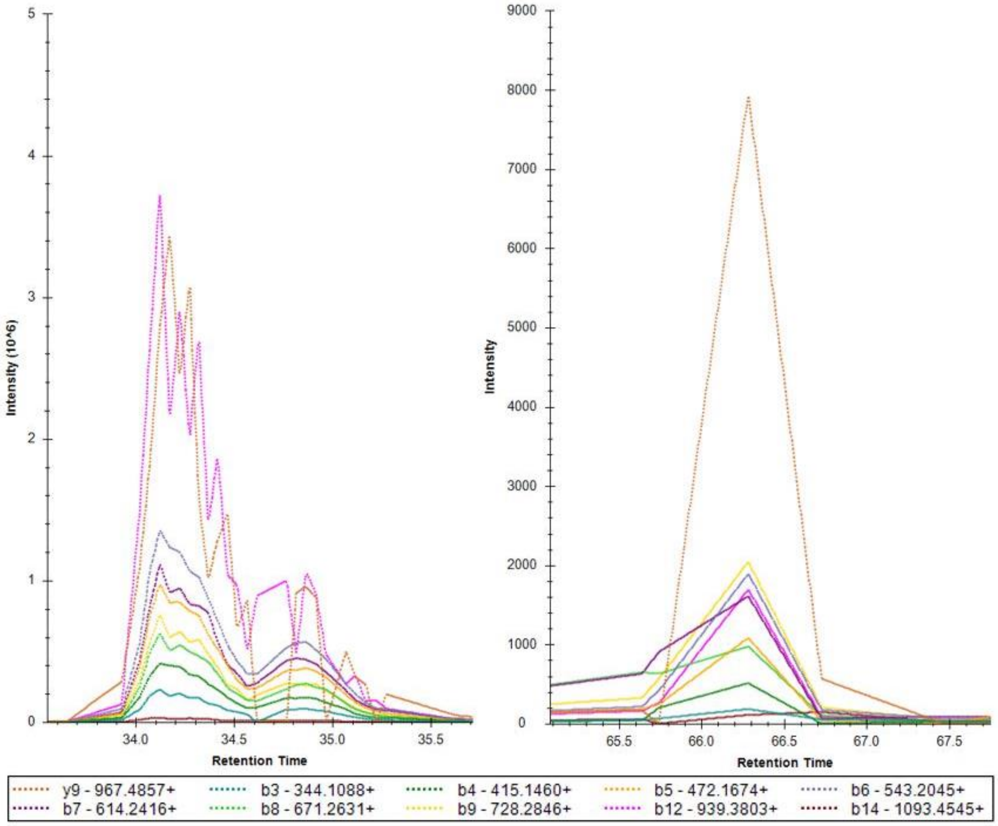


**B**

ASTGTAKAVGKVIPELNGKL
Ace-ASTGTAK<AcD4*>AVGK<AcD4*>VIPELNGK<AcD4*>L-COOH

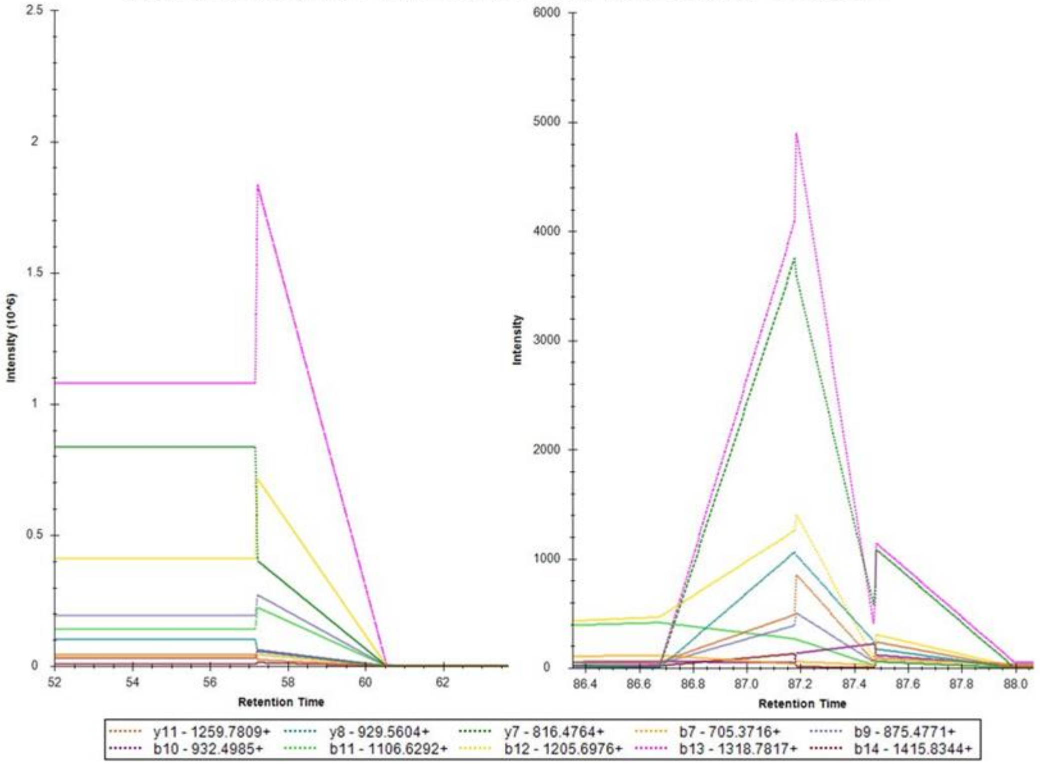


C

FPSNWNEIVDSFDDMNLSESLLR
Ace-FPSNWNEIVDSFDDM<Mox*>NLSESLLR-COOH

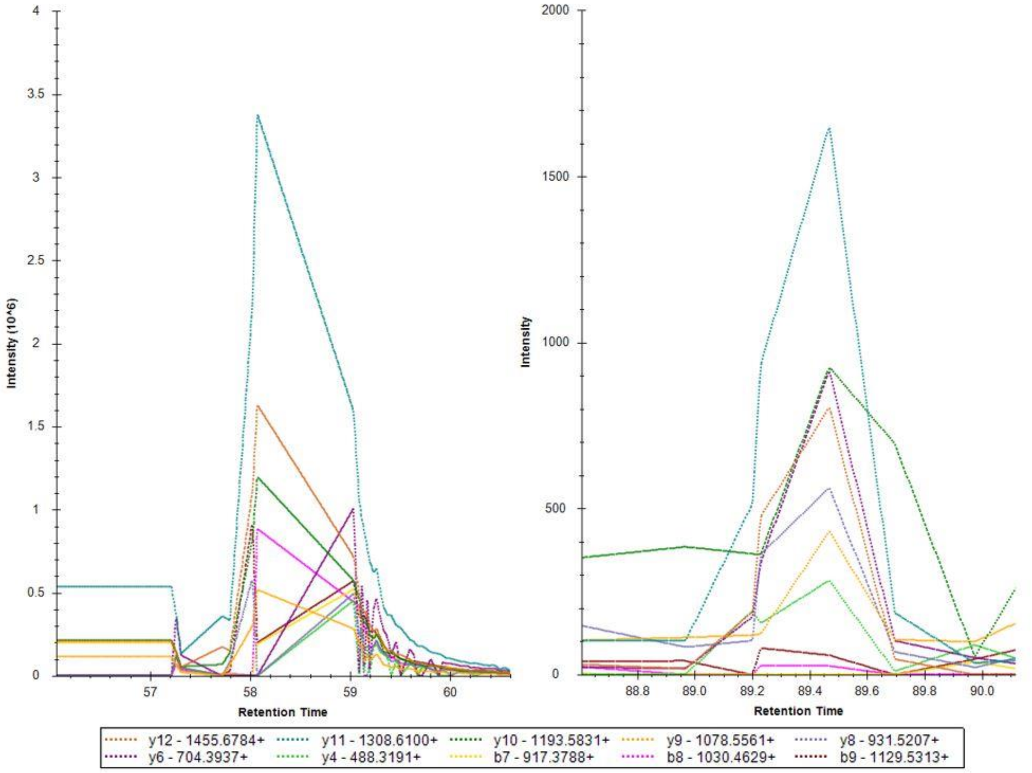

**D**

GDVVPKDANAAIATIKTKR

Ace-GDVVPK<AcD4*>DANAAIATIK<AcD4*>TK<AcD4*>R-COOH

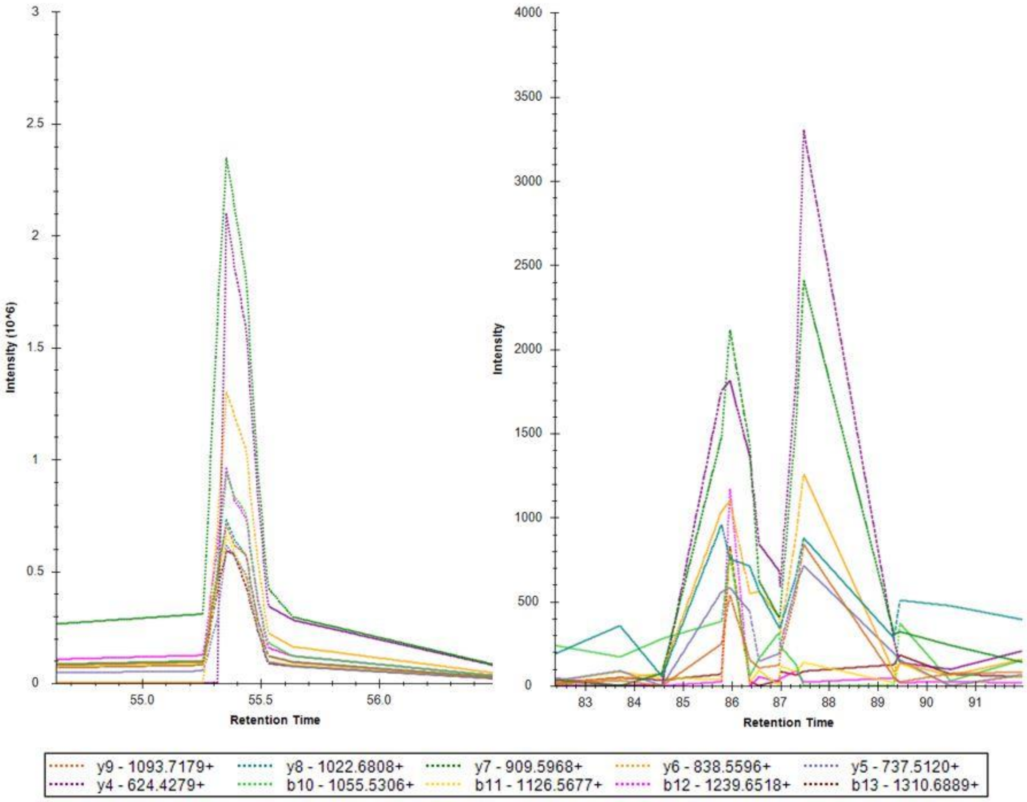

**E**

PKDANAAIATIKTKR

Ace-PK<AcD4*>DANAAIATIK<AcD4*>TK<AcD4*>R-COOH

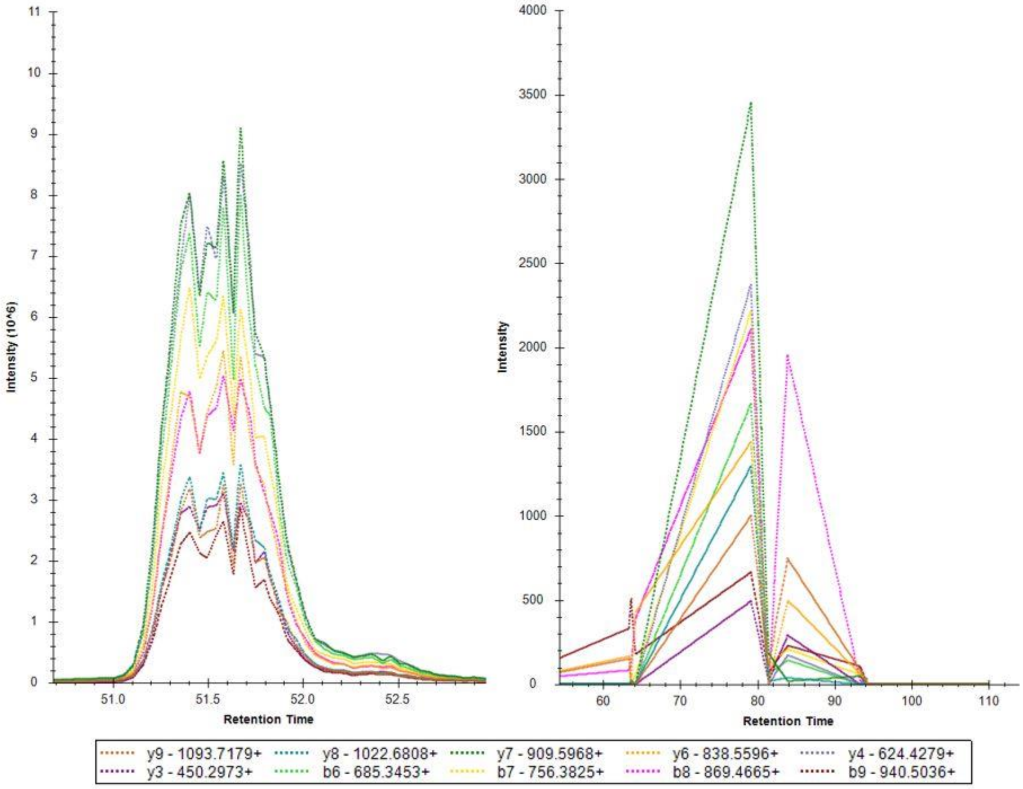

**F**

MKEETKEDAEEKQ

AcD4-M<Mox*>K<AcD4*>EETK<AcD4*>EDAEEK<AcD4*>Q-COOH

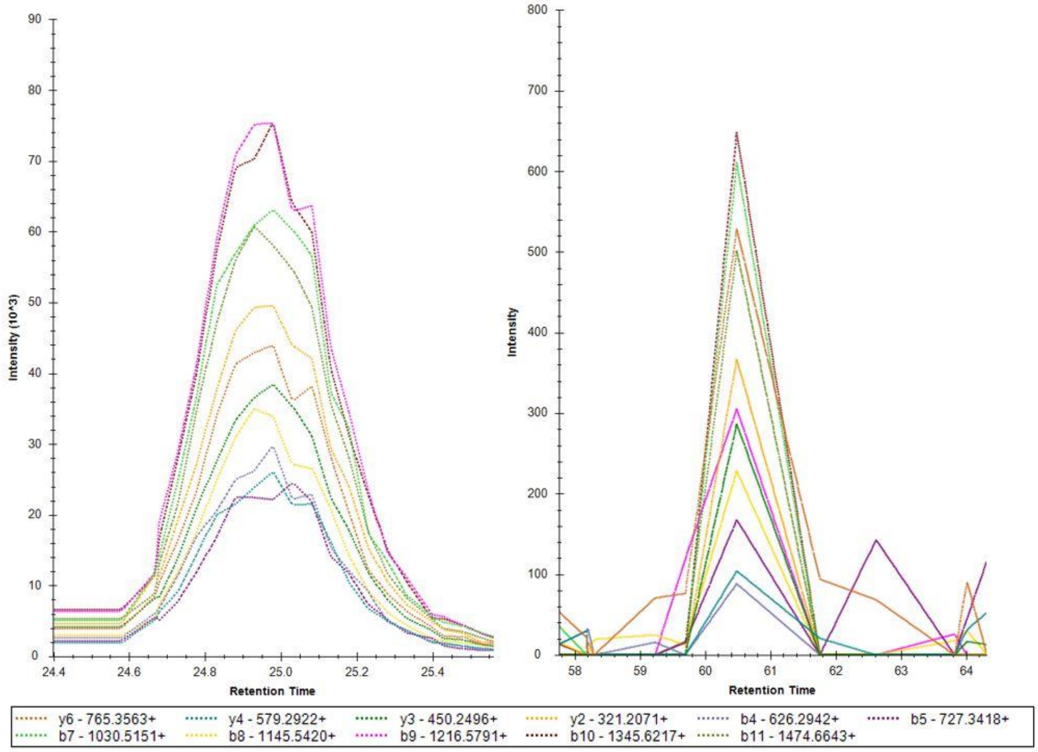


Figure S4: Comparison based on the ranking of the top 10 fragment ions of the synthetic peptide and the NTR-derived peptide identified in our COFRADIC samples. A) For ADDAGAAGGPGGPGGPEMGNRGGFRGGF, B) ASTGTAKAVGKVIPELNGKL, C) FPSNWNEIVDSFDDMNLSESLLR, D) GDVVPKDANAAIATIKTKR, E) PKDANAAIATIKTKR and F) MKEETKEDAEEKQ. The modified peptide sequence is indicated at the top of each spectrum. The top ten fragment ions (or transitions), indicated with different colors at the bottom of the spectrum, were used as comparison between the synthetic (left) and identified (right) peptide.

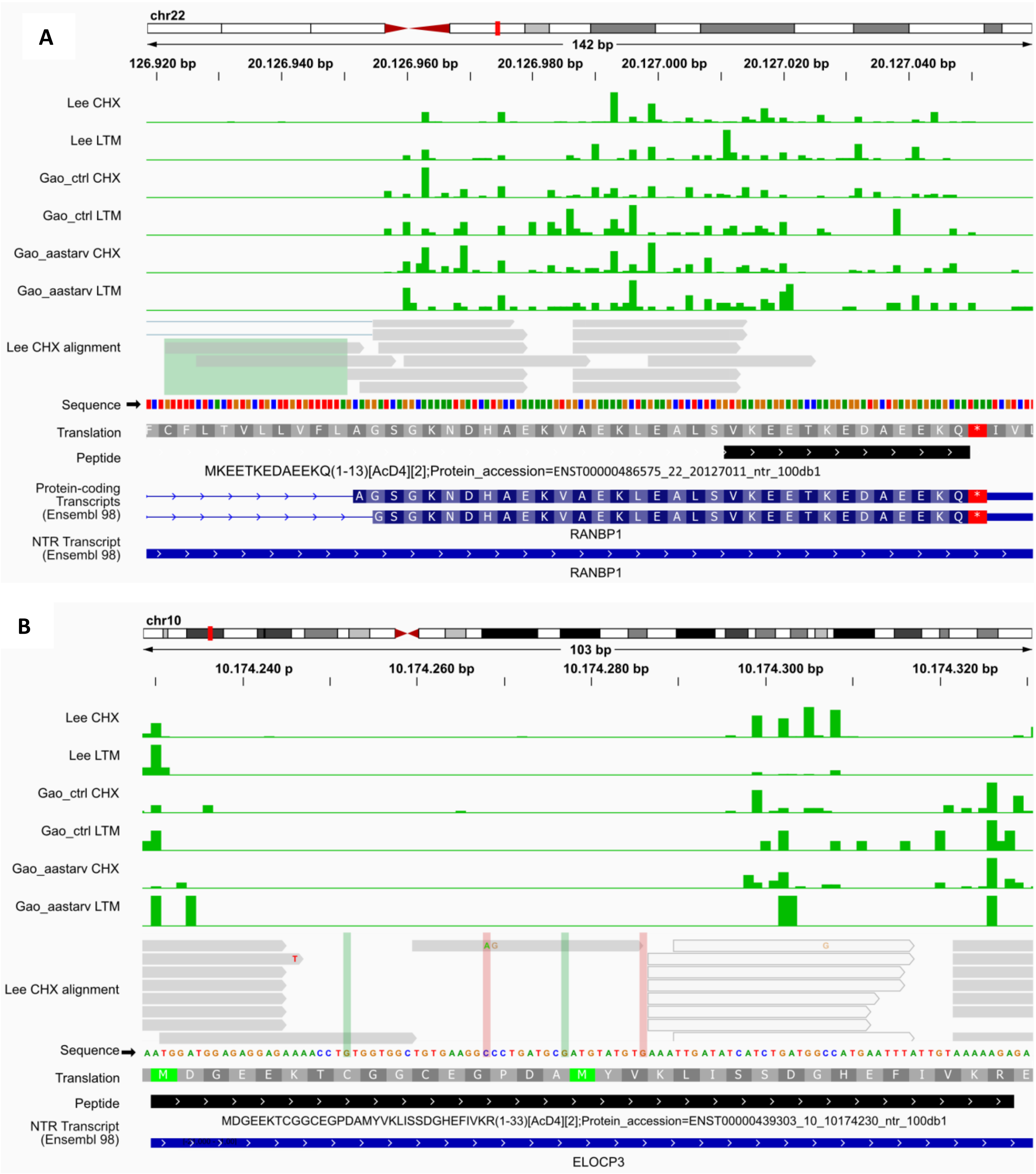
A
chr22
142 bp
126.920 bp
20.126.940 bp
20.126.960 bp
20.126.980 bp
20.127.000 bp
20.127.020 bp
20.127.040 bp
Lee CHX
Lee LTM
Gao_ctrl CHX
Gao_ctrl LTM
Gao_aastarv CHX
Gao_aastarv LTM
Lee CHX alignment
Sequence
Translation
Peptide
MKEETKEDAEEKQ(1-13)[AcD4][2];Protein_accession=ENST00000486575_22_20127011_ntr_100db1
Protein-coding Transcripts (Ensembl 98)
RANBP1
NTR Transcript (Ensembl 98)
RANBP1
B
chr10
103 bp
10.174.240 p
10.174.260 bp
10.174.280 bp
10.174.300 bp
10.174.320 bp
Lee CHX
Lee LTM
Gao_ctrl CHX
Gao_ctrl LTM
Gao_aastarv CHX
Gao_aastarv LTM
Lee CHX alignment
Sequence
AATGGATGGAGAGGAGAAAACCTGTGGTGGCTGTGAAGGCCCTGATGCGATGTATGTGAAATTGATATCATCTGATGGCCATGAATTTATTGTAAAAAGAGA
Translation
M D G E E K T C G G C E G P D A M Y V K L I S S D G H E F I V K R E
Peptide
MDGEEKTCGGCEGPDAMYVKLISSDGHEFIVKR(1-33)[AcD4][2];Protein_accession=ENST00000439303_10_10174230_ntr_100db1
NTR Transcript (Ensembl 98)
ELOCP3

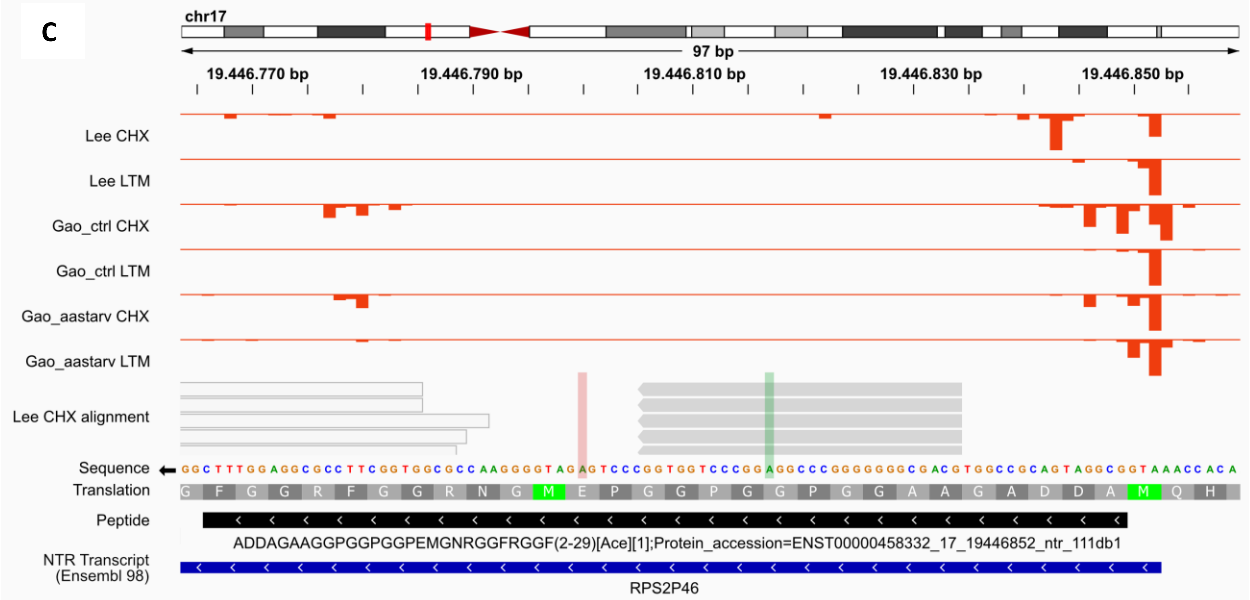
C
chr17
97 bp
19.446.770 bp
19.446.790 bp
19.446.810 bp
19.446.830 bp
19.446.850 bp
Lee CHX
Lee LTM
Gao_ctrl CHX
Gao_ctrl LTM
Gao_aastarv CHX
Gao_aastarv LTM
Lee CHX alignment
Sequence
Translation
Peptide
ADDAGAAGGPGGPGGPEMGNRGGFRGGF(2-29)[Ace][1];Protein_accession=ENST00000458332_17_19446852_ntr_111db1
NTR Transcript (Ensembl 98)
RPS2P46

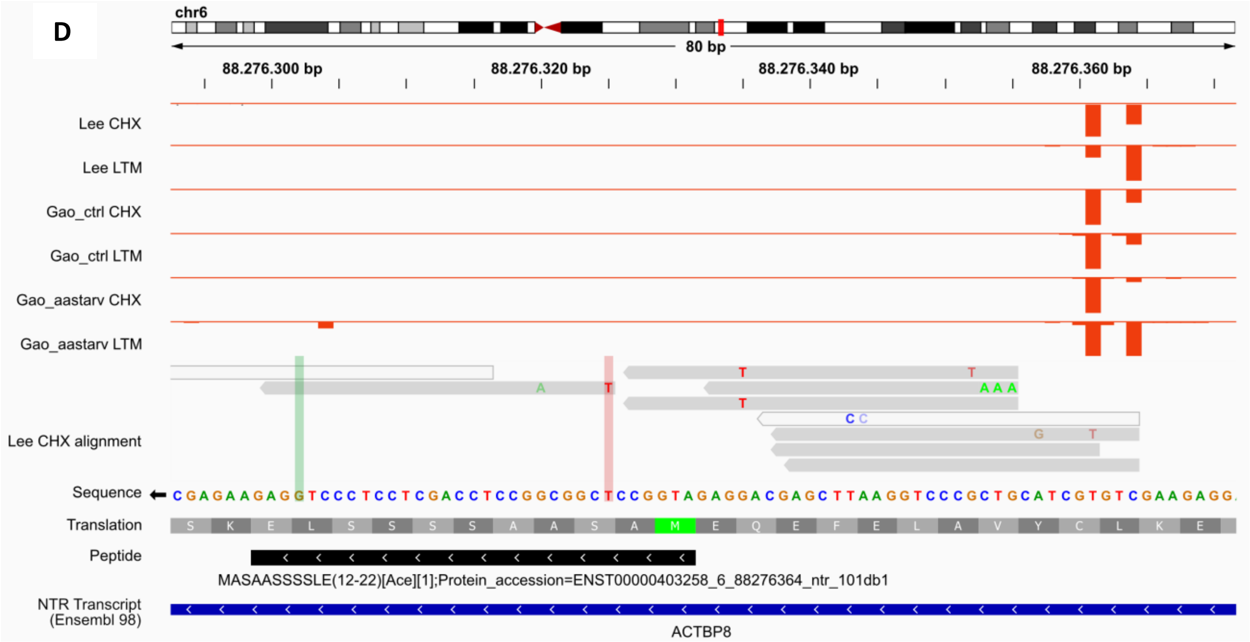
D
chr6
80 bp
88.276.300 bp
88.276.320 bp
88.276.340 bp
88.276.360 bp
Lee CHX
Lee LTM
Gao_ctrl CHX
Gao_ctrl LTM
Gao_aastarv CHX
Gao_aastarv LTM
Lee CHX alignment
Sequence
Translation
Peptide
MASAASSSLE(12-22)[Ace][1];Protein_accession=ENST00000403258_6_88276364_ntr_101db1
NTR Transcript (Ensembl 98)
ACTBP8

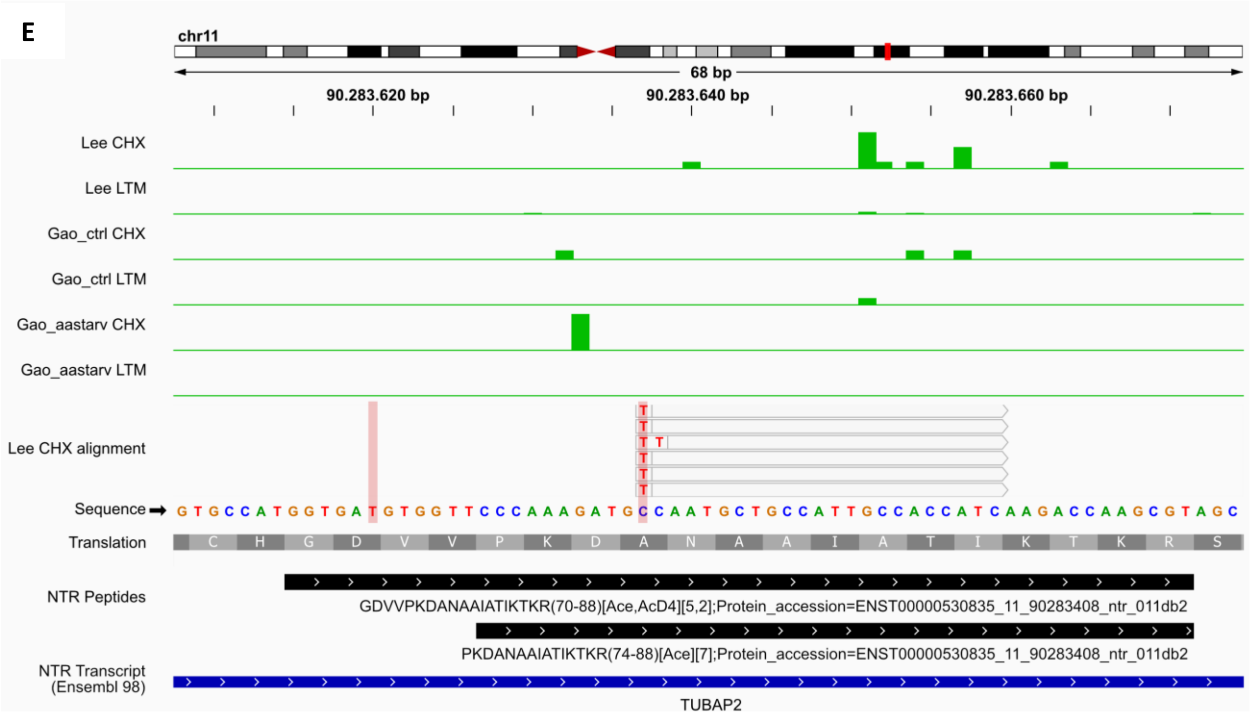


**Figure S5. Omics evidence for NTR proteoform expression visualized using a genome browser.** The six top tracks represent ribosome profiling evidence of translation in Watson (green) or Crick orientation (red). We used two published studies (Lee *et al*. (4) and Gao *et al*. (34)) as source of data for elongating ribosomes treated with cycloheximide (CHX) and initiating ribosomes treated with lactimidomycin (LTM), harvested from HEK cells under normal conditions ("Lee" and "Gao_ctrl") or under amino acid deprivation ("Gao_aastarv"). The following track displays Ribo-seq read alignment and/or the experimentally verified splice junctions. NTR-specific Ribo-seq evidence was displayed onto the alignment track using colored rectangles: green (to highlight NTR-specific variants covered by reads) or red (to mark unsupported positions). Next, transcripts (from Ensembl annotation) and genome-mapped peptides (from current study) are shown. Increasing line thickness represents introns, exons and CDS, respectively and arrows mark the direction of translation. Peptide name consists of peptide sequence, start - end position, N-terminal modification, spectral count and matching protein accession. **A.** MKEETKEDAEEKQ peptide of the RAN binding protein 1 (RANBP1) derived from a retained intron transcript carries no single-nucleotide variants compared to protein-coding transcripts of that gene, but is supported by NTR transcript-specific, intron retaining reads. **B.** MDGEEKTCGGCEGPDAMYVKLISSDGHEFIVKR peptide of the Elongin C pseudogene 3 (ELOCP3) derived from a processed pseudogene transcript is supported by one non-synonymous and one synonymous variant. **C.** ADDAGAAGGPGGPGGPEMGNRGGFRGGF peptide of the Ribosomal protein S2 pseudogene 46 (RPS2P46) derived from a processed pseudogene transcript is supported by NTR-specific reads with a synonymous variant, but lacks evidence for the non-synonymous variant. **D.** MASAASSSLE peptide of the ACTB pseudogene 8 (ACTBP8) derived from a processed pseudogene transcript is supported by a synonymous variant, but lacks evidence for the non-synonymous variant. **E.** GDVVPKDANAAIATIKTKR and PKDANAAIATIKTKR peptides of the Tubulin alpha pseudogene 2 (TUBAP2) derived from a processed pseudogene transcript lack evidence for the NTR-specific variants
